# Optimized photochemistry and enzymology enable efficient analysis of RNA structures and interactions in cells and virus infections

**DOI:** 10.1101/2020.04.30.071167

**Authors:** Minjie Zhang, Kongpan Li, Willem A. Velema, Jianhui Bai, Chengqing Yu, Ryan van Damme, Wilson H. Lee, Maia L. Corpuz, Jian-Fu Chen, Zhipeng Lu

**Affiliations:** Department of Pharmacology and Pharmaceutical Sciences, School of Pharmacy, University of Southern California, Los Angeles, CA 90033; Institute for Molecules and Materials, Radboud University Nijmegen, The Netherlands; Center for Craniofacial Molecular Biology, University of Southern California, Los Angeles, CA 90033, USA

## Abstract

Direct determination of RNA structures and interactions in living cells is critical for understanding their functions. Current crosslinking and proximity-ligation approaches are fundamentally limited due to inefficient RNA crosslinking, purification and high-level photochemical damages. Here we present PARIS2 (psoralen analysis of RNA interactions and structures, second generation), a re-invented method for capturing RNA duplexes in cells with three orders of magnitude improved efficiency. PARIS2 captures ribosome small subunit (SSU) binding sites on mRNAs, reporting translation status on a transcriptome wide scale, and captures spliceosomal snRNP binding sites on various RNA targets. We determine the RNA genome structure of enterovirus D68, a re-emerging viral pathogen associated with severe neurological symptoms, and discover alternative conformations in the internal ribosome entry site (IRES) that controls translation initiation. Together, these results reveal new aspects of RNA photochemistry and enzymology, and enable highly efficient interrogation of the RNA structurome and interactome in cells.

## INTRODUCTION

RNA structures and interactions play important roles in many cellular processes, ranging from carrying genetic information, catalysis, regulation of gene expression, and beyond^1^. However, the vast majority of RNA molecules are too large and flexible for structure analysis using the physical methods such as X-ray crystallography, NMR and cryo-EM^2^. Base pair stacking is the dominant force in RNA structures and RNA-RNA interactions; therefore, direct determination of base pairs is a critical step towards decoding the structural basis of RNA-mediated regulation in cells.

Recently, we and others developed new approaches to determine RNA base pairs, based on the principle of crosslinking, proximity ligation and high throughput sequencing^3–6^. These methods, including PARIS, SPLASH, LIGR-seq and COMRADES, allowed direct analysis of RNA duplexes at the transcriptome level, achieving single molecule accuracy and near base pair resolution. Application of these methods has led to new insights into the mechanisms and functions of cellular and viral RNAs, such as modular architectures of long noncoding RNAs, and dynamic structures/interactions of RNA virus genomes^3, 4^.

Despite over 50 years of research on the nucleic acid crosslinking, our understanding of the physical, chemical, and enzymatic properties of “crosslink-ligation” methods remain limited. The solubility of the commonly used psoralen AMT (aminomethyl trioxalen) is low, limiting its crosslinking efficiency. Ultraviolet (UV) crosslinking and reversal induce severe damage to RNA^7^, impeding reverse transcription. We discovered that crosslinked RNA cannot be recovered efficiently from cells using the classical AGPC (acid guanidine thiocyanate phenol chloroform) aqueous-organic phase separation method (commercially known as TRIzol, QIAzol, etc.) or silica-based solid phase extraction methods^8, 9^. Given the low efficiency, several methods have been developed to enrich crosslinked fragments, including native-denatured two-dimension (ND2D) gel, biotin-tagging and RNase R treatment, however, these approaches are often expensive and inefficient^2^.

We have now systematically investigated the basic physics and chemistry of major steps in the crosslink-ligation methods for RNA duplex discovery; and develop a new generation of the PARIS method (PARIS2). In particular, we identify amotosalen as an efficient and significantly improved crosslinker compared to the commonly used psoralen AMT, due to its higher solubility. We discover that crosslinking increases RNA hydrophobicity, and design a new method, TNA, to purify crosslinked RNA, enabling targeted analysis of RNAs using antisense enrichment. We develop a denatured-denatured 2D (DD2D) gel system for isolation of pure crosslinked RNA without the need for tagging the crosslinker. We introduce new chemical and enzymatic approaches to prevent and bypass photochemical damages to RNA. Together, these optimizations in PARIS2 resulted in >4000-fold increased efficiency, and importantly, the individual improvement will also find broad use in RNA research. Applying PARIS2, we discover that crosslinked RNA fragments can report translation status of mRNAs, and profile global snRNP binding sites. We also use PARIS2 to determine the genome architecture of enterovirus EV-D68, an important re-emerging pathogen associated with severe neurological symptoms, and discover novel structure conformations in enterovirus D68. The new PARIS2 method will enable more rapid and facile analysis of structural basis of RNA functions in various biological systems.

## RESULTS

### Overview of the PARIS2 strategy and major improvements

The crosslinking and proximity ligation-based principle for RNA secondary structure and interaction analysis relies on the successful completion of multiple reaction and extraction steps (**Fig. 1a**). The process starts with psoralen crosslinking in live cells, followed by RNA extraction and fragmentation, isolation of crosslinked from non-crosslinked, proximity ligation, crosslink reversal, adapter ligation, reverse transcription and finally cDNA amplification. In this study, we performed a systematic analysis of each step and make improvements based on the newly discovered physical, chemical and enzymatic properties of RNA reactions and extractions (summarized in **Fig. 1b**). The new improvements include (1) high solubility and high efficiency crosslinker amotosalen, (2) complete extraction of crosslinked RNA, (3) simplified RNA fragmentation using RNase III, (4) DD2D gel selection of crosslinked RNA, (5) optimized adapter ligation, and (6) prevention and bypass of photochemical RNA damage. Together these changes lead to >4000-fold improvement in the efficiency for PARIS2 (**Supplementary Tables 1,2**). Major optimizations are presented below, whereas additional details are in Supplementary Notes.

**Figure 1.**
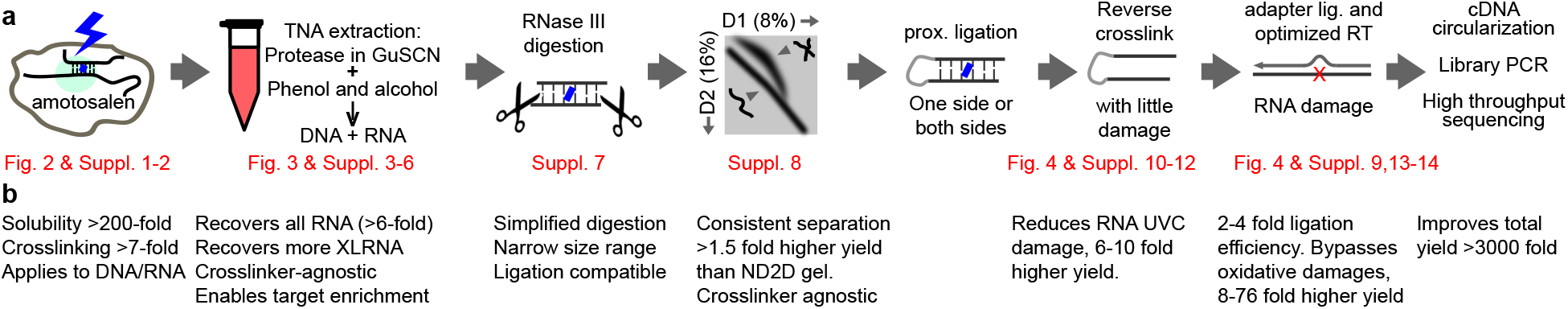
PARIS2 outline and summary of improvements. a, Outline of the PARIS method and major improvements. Details of the improvements are presented in subsequent figures, supplementary figures and tables. **b**, Notes of the major advantages of the new method and and fold improvement for each step. Amotosalen used at 5mg/ml (10 fold higher than AMT), achieves >7 fold higher efficiency. Improvement of DD2D gel efficiency over biotin-tag purification is even higher. The prevention of UVC and bypass of PUVA damages depends on RNA length, and the improvement is more remarkable for longer RNAs.

### Highly soluble psoralen amotosalen increases RNA crosslinking efficiency

The most commonly used psoralen AMT is only soluble at 1mg/ml in aqueous solutions, limiting its crosslinking efficiency^10^ (**Fig. 2a**). Amotosalen, a previously reported psoralen derivative, is soluble at above 50mg/ml, and its activity in virus inactivation is similar to AMT at the same concentration^11^ (**Fig. 2a)**. We designed a new 3-step method to synthesize amotosalen-HCl and found it soluble at 230mg per ml in water and >100mg per ml in phosphate buffered saline (PBS) (**Supplementary Fig. 1**). AMT and amotosalen crosslink DNA oligo duplexes in vitro with similar efficiency at the same concentration (**Supplementary Fig. 2**). At higher concentrations, amotosalen is more efficient.

**Figure 2.**
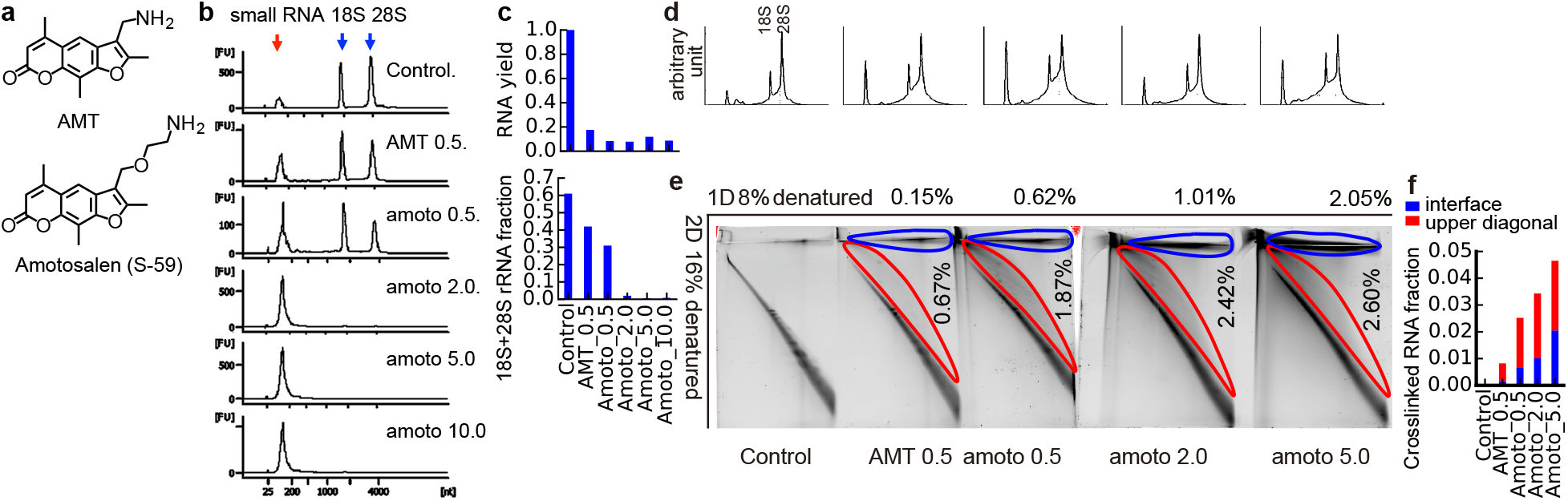
High solubility psoralen amotosalen increases crosslinking efficiency. **a**, Structures of two psoralen derivatives, AMT and amotosalen (also known as S-59). **b**, Higher concentrations of crosslinkers increases crosslinking efficiency. HEK 293 cells crosslinked with increasing concentrations of psoralens (mg/ml) are extracted using the standard AGPC (TRIzol) method to recover RNA from the aqueous phase of the TRIzol-chloroform mixture. Purified RNA was analyzed using Bioanalyzer. small RNA: RNA in the range of 50-300nt. **c**, Total RNA yield (upper panel) and percentage of 18S+28S rRNA (lower panel) from the experiments shown in panel b. **d-e**, Total RNA extracted from HEK293 cells using the TNA method are analyzed using TapeStation (**d**) and DD2D gel (**e**). Percentages of crosslinked RNA extracted from the upper diagonal (red outlines) and the 1D-2D interface (blue outlines) are indicated. **f**, Percentages of RNA from the upper diagonal the in the second dimension gel and the 1D-2D interface.

We discovered that crosslinking repartitions large RNA from the aqueous phase to the interphase during standard AGPC (TRIzol) extraction (**Fig. 2b**). The migration of large RNAs from aqueous to the interphase serves as an indicator for the crosslinking efficiency. We crosslinked cells with AMT or amotosalen at various concentrations and extracted RNA from the aqueous phase in TRIzol-chloroform mixture. Both total RNA yield and percentage of 18S+28S rRNAs were reduced with higher concentrations of psoralens, suggesting higher crosslinking efficiency (**Fig. 2c**).

To accurately measure crosslinking efficiency, we recovered crosslinked intact RNA using the newly invented TNA method (**Fig. 2d**), fragmented RNA using an optimized RNase III protocol, and extracted crosslinked but not monoadduct fragments using the new DD2D gel system (see details later). Total crosslinked fragments (including both 1D-2D interface and 2D upper diagonal) increased from 0.82% to 4.65%, roughly 5.7-fold, after a 10-fold increase in psoralen concentration (AMT 0.5 vs. amotosalen 5). Even at the same concentration (0.5mg/ml), amotosalen is more efficient than AMT. The RNA duplexes captured by AMT and amotosalen are similar (**Supplementary Fig. 2c-e**). Therefore, we identified amotosalen as a more efficient nucleic acid crosslinker.

### Phase partition and extraction of crosslinked RNA

In the classical AGPC method for RNA extraction, the mixture of guanidine thiocyanate (GuSCN), phenol and chloroform forms two phases^8^. RNA partitions to the aqueous phase at pH below 5, proteins partition to the inter and organic phase, while DNA partitions to the interphase (**Fig. 3a-b, Supplementary Note 1**). On the other hand, the standard PCI (phenol, chloroform and isoamyl alcohol) extraction of DNA employs higher pH (~8.0) to bring both DNA and RNA to the aqueous phase. While applying the AGPC method (commercial name TRIzol, etc.), we noticed that crosslinked cells cannot be completely dissolved, and RNA yield was greatly reduced^9^ (**Supplementary Fig. 3a**). Proteinase K (PK) treatment was necessary but insufficient for improving RNA yield (by only ~10%). We used PK and RNase digestion in lysate to recover crosslinked RNA^9^, however, recovery was still incomplete. Furthermore, fragmentation prior to purification makes it difficult to enrich specific RNAs using antisense oligos.

**Figure 3.**
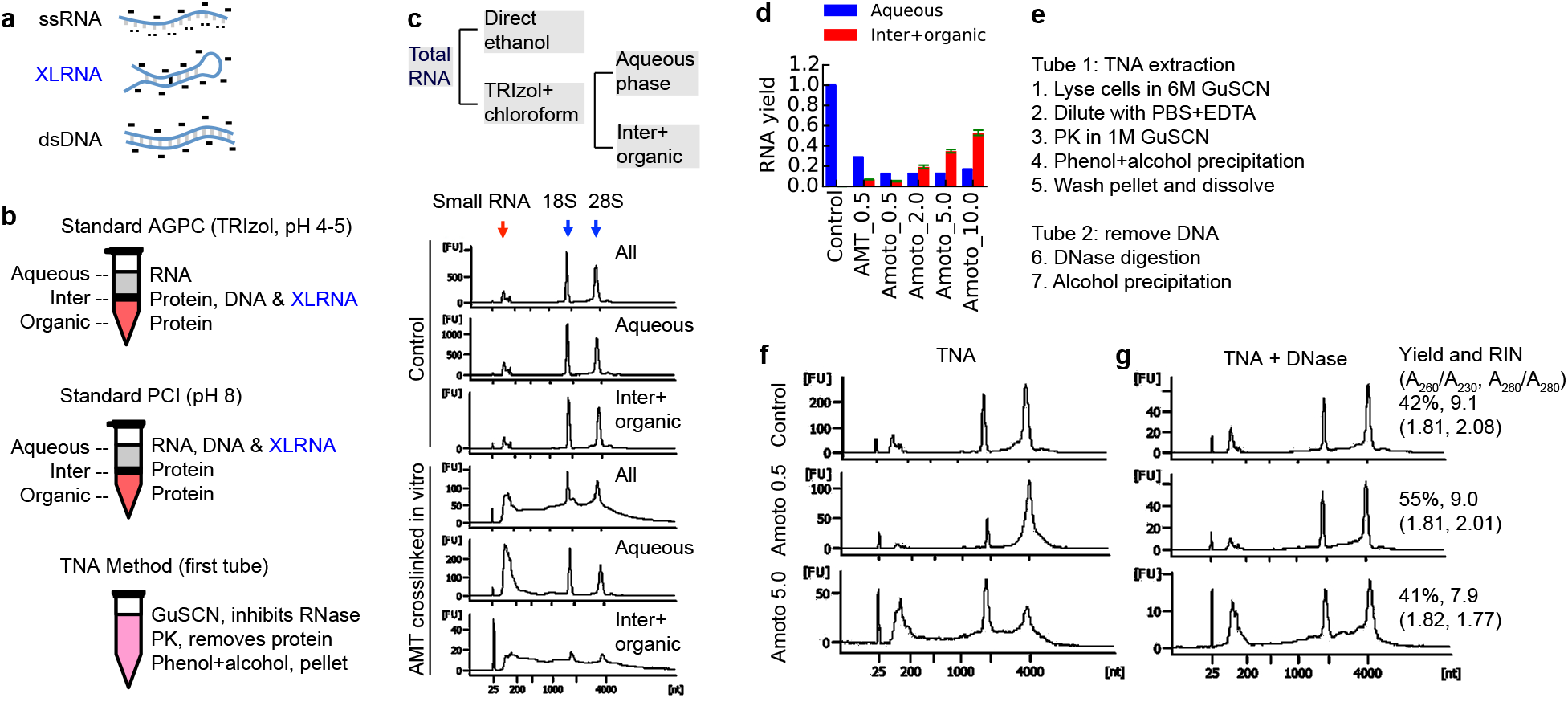
TNA: a new method recovers crosslinked RNA from cells. **a**, Charge and hydrogen bonding of RNA and DNA molecules in standard AGPC (TRIzol) purification. The minus sign ‘-’ indicates negative charges on DNA and RNA. The ‘..’ indicates the exposed hydrophilic bases involved in hydrogen bonding. **b**, Phase partition of DNA, RNA and protein in standard AGPC and PCI (phenol chloroform isopropanol) extraction of nucleic acids, and comparison to the one-phase method TNA. c, Repartition of crosslinked RNA to the interphase during Trizol extraction. Purified total RNA was treated with or without 365nm UV plus AMT, and then either directly precipitated using ethanol and sodium acetate, or extracted using the TRIzol method. RNA from the aqueous or inter+organic phases are precipitated with ethanol and sodium acetage. Small RNA: RNAs in the range of 50-300nt, including tRNA, snRNA, 5S and 5.8S rRNAs etc. **d**, Recovery of crosslinked RNA from the aqueous and inter+organic phases using the TNA method. **e**, Outline of the TNA method. HEK293 cells were crosslinked using 0.5 or 5mg/ml amotosalen and then extracted using the TNA method. Size distributions were analyzed before (f) and after (g) DNase treatment. Yield (RNA/TNA), RNA integrity numbers (RIN) and two indicators of RNA quality, A_260_/A_230_ and A_260_/A_280_, were indicated on the right. In vivo crosslinking is less efficient than in vitro, hence the lower smear compared to panel **c**.

We suspected that crosslinked RNA may be more hydrophobic and partitioned to the interphase. To test this possibility, we crosslinked pure total RNA with AMT and then extracted RNA using direct ethanol precipitation (**Fig. 3c**). Alternatively, we used the AGPC method (TRIzol), where RNA from the aqueous, inter and organic phases was precipitated separately. Crosslinking induced a broad smear that spans beyond the largest 28S peak. While direct ethanol precipitation recovered all RNA, the aqueous phase in TRIzol-chloroform mixture contains only RNA in the 50-300nt range (e.g. tRNAs, snRNAs, snoRNAs), and sharp non-crosslinked 18S and 28S peaks (**Fig. 3c-d**). These results suggested that crosslinking increased RNA hydrophobicity, making it difficult to extract using the classical AGPC method. In most previous studies, the inefficient recovery of crosslinked RNA likely has resulted in significant bias because larger and heavier crosslinked RNA molecules are lost.

To recover total RNA after crosslinking, we developed a new method termed total nucleic acid extraction (TNA). Briefly, cells are first lysed in 6M GuSCN to completely inhibits nucleases. The lysate is diluted to reduce GuSCN concentration, buffered, treated with EDTA to chelate divalent cations, and digested with PK to remove proteins, which also inactivates nucleases. Total nucleic acids are then precipitated using phenol and alcohol. Phenol keeps residual proteins in solution so that only nucleic acids precipitate. DNA that makes up about 50% of all nucleic acids is removed using DNase, leaving intact RNA (**Supplementary Fig. 3c**). Carmustine and chlorambucil, two cancer chemotherapy drugs that crosslink nucleic acids, also promoted the partition of RNA to the interphase, and crosslinked RNA was successfully recovered using the TNA method (**Supplementary Figs. 4-5**). TNA also outperforms both AGPC and solid-phase methods by at least 6-fold (**Supplementary Fig. 6, Supplementary Tables 1,2**). These results suggest that crosslink-induced hydrophobicity is a general property of crosslinked RNA. Therefore, TNA method is generally applicable to crosslinking studies and enables targeted antisense enrichment from intact total RNA (see details later).

### Efficient isolation of crosslinked RNA using a DD2D gel system

To obtain short crosslinked RNA fragments, we developed a simplified one-step RNase III protocol that takes advantage of the digestion kinetics (**Supplementary Fig. 7, Supplementary Note 2**). Given the low efficiency of psoralens, crosslinked fragments need to be enriched for sequencing. Biotin-conjugated psoralens have been used to enrich RNA after crosslinking, but these methods also enrich monoadducts, which are more abundant than crosslinks^4, 6^. Tag-based purification requires custom synthesis and cannot be quickly adapted for other crosslinkers. RNase R depletion of non-crosslinked RNA is also impeded by monoadducts^5^. We initially used a ND2D gel system to isolate crosslinked RNA without monoadduct contamination ^9, 12^, however, this method suffers from low resolution and low yield (**Supplementary Fig. 8**). The diagonal in the second dimension is broad, partially masking crosslinked fragments above the diagonal. We developed a new DD2D gel system takes advantage of the differential migration of crosslinked RNA vs. non-crosslinked at different gel concentrations during electrophoresis (**Supplementary Fig. 8** and **Supplementary Note 3**). The DD2D method has higher resolution and consistency, recovers more crosslinked fragments (>1.5 fold vs. ND2D), does not rely on the base pairing of the crosslinked RNA for separation. RNA duplexes crosslinked with other compounds can be separated from noncrosslinked, suggesting that DD2D is a broadly applicable method (**Supplementary Figs. 4-5**).

### Prevention of RNA against UVC induced photochemical damages

Photochemical crosslinking (psoralen + UVA, or 365nm) and reversal (UVC, 254nm) enable in vivo analysis of RNA duplexes, but also cause many types of damage. Together with the low efficiency proximity ligation, the damages block reverse transcription and reduce both the total cDNA yield and percentage of gapped reads (**Fig. 4a**). UVC irradiation induces pyrimidine dimers and other damages via the singlet excited state, even after very short exposure^7, 13^ (**Fig. 4b**). Earlier studies showed that UVC induced DNA damage can be prevented by singlet state quenchers, but such approaches did not work well for RNA^14, 15^. To prevent, repair or bypass UVC-induced RNA damages, we systematically screened a variety of conditions, including intercalating dyes and solvents that act as singlet quenchers (**Fig. 4c, Supplementary Figs. 9-11** and **Supplementary Note 4**). Superscript IV (SSIV) reverse transcriptase outperforms other enzymes on UVC damaged RNA, increasing yield by 7-fold over SSIII (**Supplementary Figs. 9**). Acridine orange (AO) and ethidium bromide (EB) at high concentrations can protect both normal and psoralen crosslinked RNA from UVC irradiation. AO effectively protects non-crosslinked RNA even after 30min UVC irradiation (at 4mW per cm^2^), after which 30% RNA remain intact, vs. 0.5% in the absence of AO (**Fig. 4c,** upper panel). For crosslinked RNA, there is simultaneous UVC-induced reversal and damage, yet AO still protects RNA effectively (**Fig. 4c**, lower panel). Importantly, the singlet quenchers did not block crosslink reversal, making it possible to apply them in PARIS-like experiments (**Supplementary Fig. 10i-l**). Together these studies demonstrate for the first time that UVC induced RNA damage can be largely prevented by high concentrations of singlet quenchers. After proximity ligation and UVC reversal of crosslinks, the RNA samples are then ligated with adapters for reverse transcription and library preparation. We also optimized the adapter ligation step using synthetic oligos (**Supplementary Fig. 12**).

**Figure 4.**
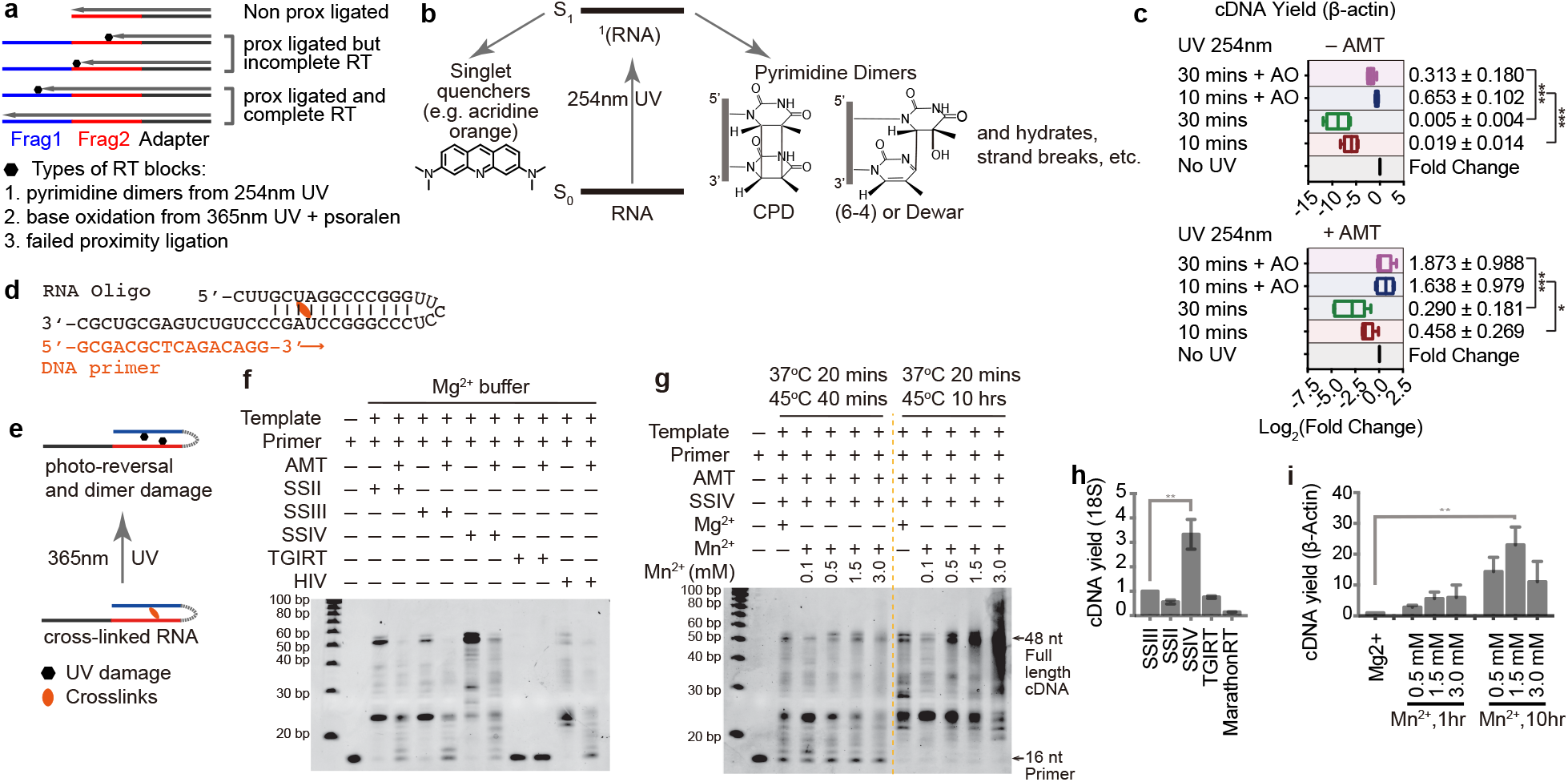
Prevention and bypass of photochemical RNA damage. **a**, Diagram showing the effects of UV damage-induced RT stops and failed proximity ligations (non prox ligated) on the yield of gapped reads. **b**, UVC (254nm) excited RNA form various products, such as pyrimidine dimers, hydrates and strand breaks. Alternatively the energy can be transferred to singlet quenchers like acridine orange. S_0_ and S_1_: ground state and excited singlet state. **c**. Yield of cDNA from RNA irradiated with 254nm UV, with or without acridine orange protection. cDNA yield is normalized to non-photo-reversal sample. Top: Non-crosslinked RNA sample. Bottom: AMT crosslinked RNA sample before 254nm treatment. **d,** Sequence of the 48nt RNA oligo template and DNA primer used for primer extension assay. **e**, UVA (365nm) reversal of crosslink simultaneously induce oxidative damages. **f,** Electrophoresis of primer extensions by different reverse transcriptases. Full extension of the primer results in a 48 nt product. Reverse transcriptases used include the Superscript series SSII, SSIII and SSIV, TGIRT-III, and recombinant HIV RT. **g**, The effect of divalent cations and incubation time on full-length cDNA synthesis by SSIV. **h**, cDNA yield obtained in RT-qPCR experiments for PUVA damaged 18s-rRNA, normalized to SSIII. ** p < 0.01. **i**, cDNA yield for β-Actin using SSIV in different reaction buffers and different incubation time, normalized to a standard Mg^2+^ condition. ** p < 0.01.

### Bypass of oxidative damages in reverse transcription

In addition to crosslinking pyrimidines, photosensitized psoralens also induce oxidative damage to RNA, primarily affecting guanines through direct electron transfer and elicitation of oxygen^16^ (**Supplementary Fig. 13**). In order to minimize the adverse effect of these damages, we systematically screened conditions to prevent, repair or bypass them (**Supplementary Figs. 13-15, and Supplementary Note 5**). Certain oxidant scavengers, such as vitamin C, Tiron and MnTBAP^17, 18^, reduced PUVA-induced oxidation, but also blocked crosslinking, due to their common energetic precursors (**Supplementary Fig. 14**). RNA damage impedes reverse transcription by trapping the enzyme in an inactive state^19, 20^. We reasoned that conditions that promote enzyme conformation dynamics, or longer incubation time, may overcome such barriers. Indeed, several conditions, including SSIV, cofactor Mn^2+^ and longer incubation time dramatically increased cDNA yield both alone or in combinations, in both primer extension assays and qRT-PCRs (**Fig. 4d-i, Supplementary Fig. 15**). SSIV outperforms all other reverse transcriptases (**Fig. 4f, h** and **Supplementary Fig. 15f**). Mn^2+^ is better than Mg^2+^ for reverse transcription (**Fig. 4g, i** and **Supplementary Fig. 15h-i**). The longer reaction time is particularly effective in promoting the bypass of damages bases (**Fig. 4i and Supplementary Fig. 15h-i**). Together, these conditions for SSIV improved the bypass of PUVA-induced damages by 8-70 folds over SSIII (**Supplementary Tables 1-2**).

### PARIS2 enables highly efficient and sensitive detection of RNA duplexes in cells

After optimizing all individual steps, we tested their performance in new PARIS2 workflow. Starting from the same number of cells, the 5mg/ml amotosalen crosslinking, TNA extraction and DD2D gel isolation improved the yield of crosslinked RNA fragments by ~60-fold over the standard AMT-TRIzol-ND2D protocol. Starting from the same amount of crosslinked RNA fragments (after DD2D gel step), the DNA library yield is improved ~76 fold (**Fig. 1, Supplementary Tables 1-2**). Together the improvements resulted in a total of >4000-fold increase in efficiency. We applied PARIS2 with oligo(dT) enrichment of cellular RNA from crosslinked HEK293 cells and mouse brain tissues, and with antisense enrichment of viral RNAs (**Figs. 5, 6**). For oligo(dT) enriched RNA, we were able to model structures of abundant mRNAs even with only ~1M gapped reads (**Supplementary Fig. 16**).

**Figure 5.**
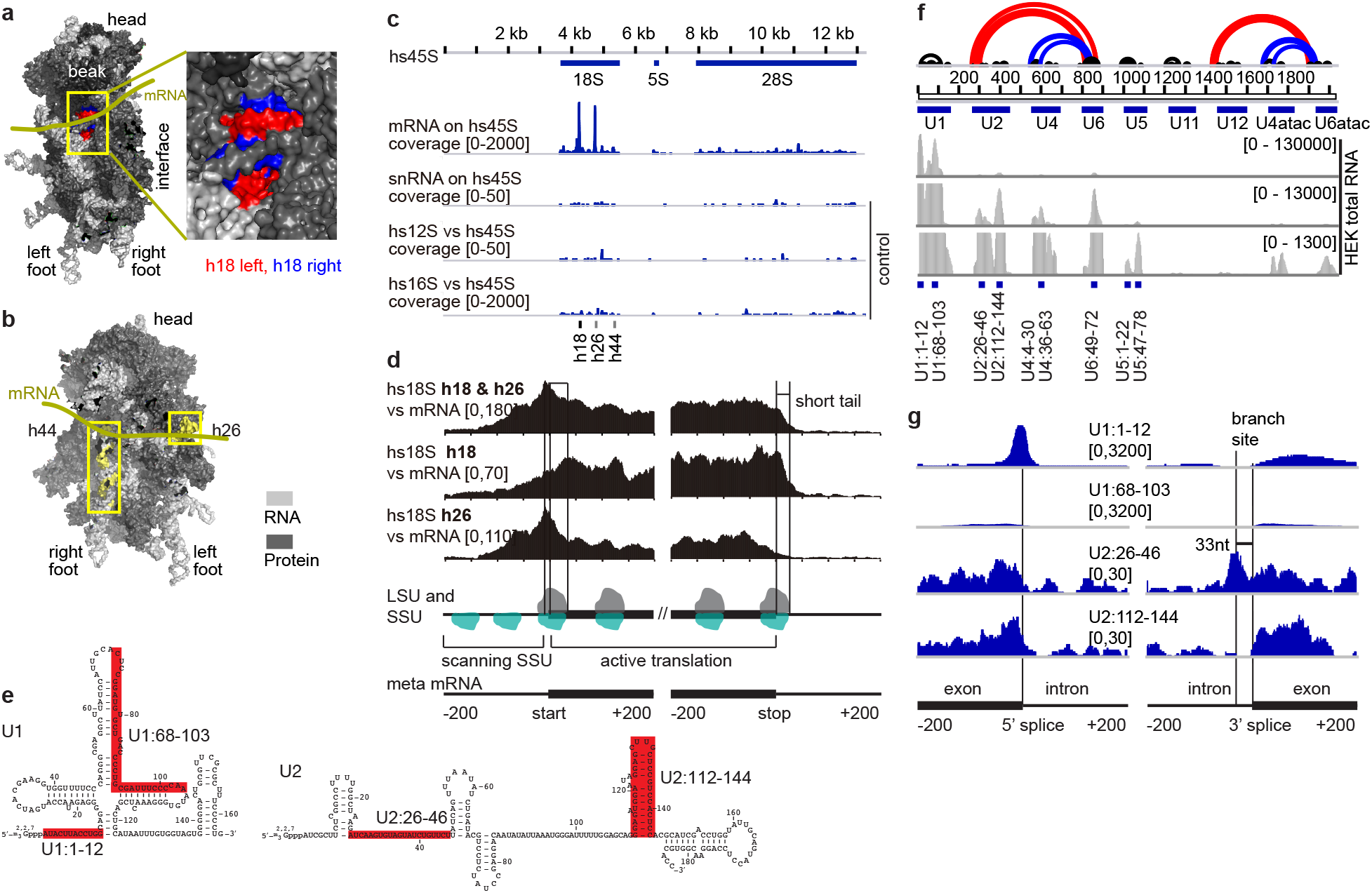
PARIS2 enables global profiling of ribosome SSU and spliceosomal snRNP binding sites. **a**, The 40S ribosome small subunit (SSU), showing the beak side view, highlighting the 18S h18 helix exposed in the mRNA channel. rRNA in light gray, RPS in dark gray, except h18 left arm: 595-620, right arm: 621-641. b, Rotated 40S, showing h26 and h44 in yellow. **c**, The binding sites of mRNAs snRNAs and mitochondrial rRNAs (12S and 16S) on the 45S unit based on HEK293 mRNA PARIS2 data. **d**, The binding sites of h18 and h26 on the meta mRNA. The tapering off signal on the 5’UTR is likely due to its limited length. The sharp drop off after the stop codon (20-30nt short tail) is due to irregular RNA fragment length. **e**, Secondary structure model of human snRNAs U1 and U2, highlighting the pre-mRNA binding regions U1:1-12 and U2:26-46, which bind the 5' splice site and the branch site respectively. U1:68-103 and U2:112-144 serve as controls. **f**, Regions in the 9 snRNAs that interact with other RNAs, based on PARIS data from total RNA, shown in 3 scales. The black and colored arcs represent intramolecular structures and intermolecular interactions. **g**, U1 and U2 binding sites on a meta pre-mRNA. U1 binding site is right at the exon-intron junction.

**Figure 6.**
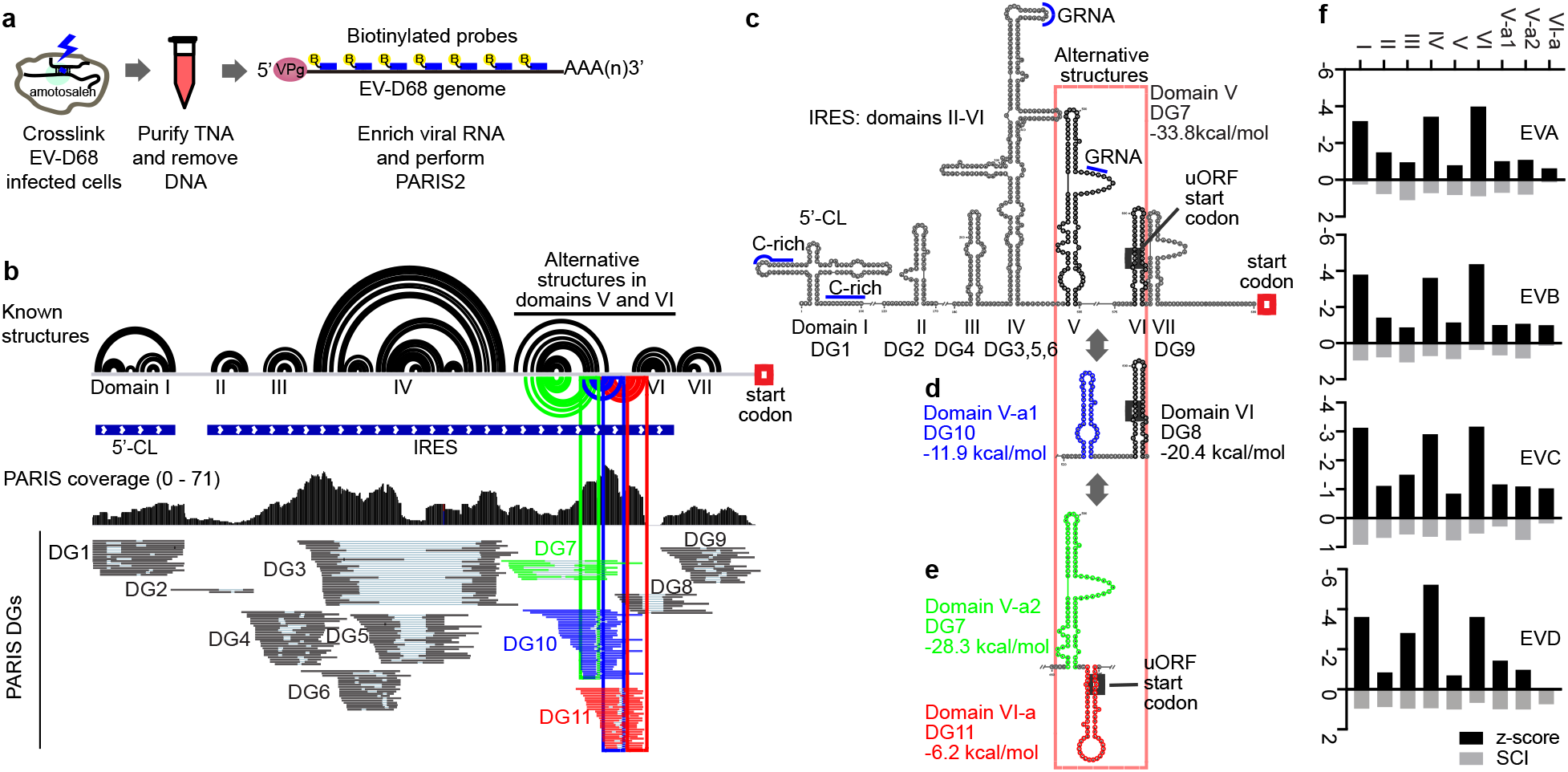
PARIS2 reveals alternative structure conformations in the IRES element of EV-D68 in cells. **a**, Schematic diagram of the experimental strategy. Virus infected HeLa cells were crosslinked. RNA was extracted using the TNA method and the viral genome RNA was enriched using biotinylated oligos. VPg: viral genome-linked protein. **b**, PARIS2 validates predicted structures (black arcs) and reveals new alternative conformations (green, red and blue arcs) in the 699nt 5’UTR of the US47 strain of EV-D68. 5’CL: cloverleaf structure. IRES: internal ribosome entry site, including domains II-VI. Duplex groups (DGs) map to all known structure domains. The 3 DGs that support the alternative conformations are color coded. **c-e**, Stemloop style structure models for the EV-D68 5’UTR. DG and domain correspondence is labeled under the model. The putative uORF and polyprotein start codons are labeled with gray and red boxes. The C-rich and GNRA motifs are labeled with blue lines. V-a1, V-a2 and VI-a: alternative conformations for domains V and VI. Minimal free energies are labeled (in kcal/mol). **f**, Analysis of structure conservation in the 5’UTR using RNAz in 4 EV species.

### PARIS2 enables profiling of ribosome SSU binding across the transcriptome

During translation, mRNAs directly contact the 18S ribosomal RNA (rRNA) in the small subunit (SSU), and some of these contacts can be crosslinked by UV alone^21^ (**Fig. 5a-b**). We reasoned that capturing the mRNA-rRNA interactions may allow direct analysis of translation. To analyze interactions with multi-copy genes, such as those encoding rRNAs, spliceosomal snRNs etc., we designed reference genomes masking multicopy genes and adding back single copies (**Supplementary Figs. 17-19**). After mapping HEK293 mRNA PARIS2 data to the engineered references, we extracted chimeric reads connecting cytoplasmic rRNAs and mRNAs. 18S rRNA regions crosslinked to mRNAs are limited to helix 18 and 26 (h18 and h26), with a minor peak on h44 (**Fig. 5c**). Both h18 and 26 are in the mRNA channel (**Fig. 5a,b** and **Supplementary Fig. 20a,b**). Such specific rRNA binding was not observed for other abundant RNAs, like the mitochondrial rRNAs and snRNAs, suggesting that interaction was captured during translation. Then we analyzed 18S h18/h26 binding sites on mRNAs (**Fig. 5d**). The strongest binding is with the coding sequence (CDS), followed by the 5’UTR, while the 3’UTR has little binding. The highest peak in right next to the start codon. Interestingly, h26 binding peak precedes that of h18, consistent with their locations in the mRNA channel, with h18 near the entry and h26 near the exit. The distance between the two peaks is around 40-50nt, slightly longer than the ribosome footprint, likely due to the random fragmentation used in PARIS2. The mRNA-rRNA crosslinking could be a result of dynamic flipping of the h18 and h26 bases that transiently pair with mRNAs, which is also necessary for direct UVC crosslinking reported in earlier studies^21^. The binding in the 5’UTR but not 3’UTR may represent the scanning phase of translation initiation which has been previously captured in translation complex profiling^22^. This is different from the standard ribosome profiling where the ribosome binding to the 5’UTR is limited to uORFs^23^.

Similar patterns of rRNA-mRNA interactions were observed in individual mRNAs and in mouse brain oligo(dT) enriched RNAs, confirming the specificity of these interactions (**Supplementary Fig. 20c-g**). We also analyzed our previous total RNA PARIS data in HEK293T cells and mouse ES cells (**Supplementary Fig. 20f-g**). Similar specific interactions were observed, despite elevated background. Together, we demonstrate PARIS2 as a powerful alternative method that enables direct analysis of mRNA translation.

### PARIS2 enables global profiling of snRNP targets

The spliceosomal snRNPs have multiple functions beyond splicing^24–28^. Accurate analysis of their binding sites across the transcriptome is necessary for mechanistic studies of these RNP complexes. Using the engineered genome references containing only single copies of snRNAs, we determined the interactions between snRNAs and other RNAs in both total and oligo(dT) enriched RNAs. Analysis of total RNA PARIS data revealed extensive interactions of snRNAs especially U1 and U2, with other RNAs (**Fig. 5e-f, Supplementary Fig. 21**). Meta analysis of snRNA binding sites on intron-containing RNAs revealed major peaks focused at the expected locations, including the 5’ splice site for U1 and the branch site for U2 (**Fig. 5g**). Interestingly, such interactions were also captured in the polyA-enriched samples, suggesting that at least some of the interactions persist after polyadenylation (**Supplementary Fig. 21b, c**). Many U1 binding sites reside in the exons, far away from the splice sites, consistent with our earlier studies^9, 24^ (**Supplementary Fig. 21d-i**). Surprisingly, we identified strong U1-XIST and U6-MALAT1 interactions, suggesting yet unknown functions of these snRNAs (**Supplementary Fig. 21d, j-k**). Together, these studies demonstrate the power in global analysis of snRNP binding sites.

### PARIS2 determines the genome structure of the enterovirus EV-D68

The genomes of RNA viruses carry the genetic information, and at the same time fold into complex structures to regulate multiple steps of their infection life cycles. However, the direct analysis of viral genome structures and interactions in cells remains challenging. We applied PARIS2 to EV-D68, an RNA virus, whose recent global outbreaks have been associated with severe respiratory symptoms and acute flaccid paralysis, which resembles poliomyelitis^29^ (**Fig. 6a)**. After *in vivo* amotosalen crosslinking of HeLa cells infected with the EV-D68 strain US/MO/14-18947 (US47), we enriched the ~7300nt viral RNA genome using biotinylated antisense probes for PARIS2 library construction (**Fig. 6a**). With ~100-fold enrichment of viral RNA, we were able to obtain full coverage of the genome using ~400,000 reads, where a quarter of them map to the virus (**Supplementary Fig. 22a-e**). The gapped reads revealed a complex global architecture, with extensive long-range structures (**Supplementary Fig. 22f**).

Using the PARIS2-derived duplex groups, we confirmed the previously predicted 5’UTR secondary structure, including the 5’CL (cloverleaf) and IRES, which play critical roles in replication and translation initiation, respectively^30, 31^ (**Fig. 6b-c, Supplementary Fig. 22g-k**). The IRES structure model consists of five domains designated II-VI. All domains are supported by multiple sequence alignments of ~500 complete EV-D genomes (**Supplementary Fig. 23**). Several motifs within the IRES, two GNRA tetraloops in domains IV and V and a pyrimidine-rich track (Yn) between domains V and VI, are essential for the recruitment of translation initiation factors^32^. These motifs are clearly identified on the PARIS2-derived structure model (**Fig. 6c**).

Interestingly, besides the proposed IRES structure, we also identified alternative structures where domain V and/or VI adopt significantly different conformations (**Fig. 6b, c-e**). These alternative conformations are supported by similar numbers of gapped reads compared to the known structure domains, suggesting that they are abundant in cells. All alternative structures are also supported by multiple sequence alignments among sequenced viral genomes in EV-D species (**Supplementary Fig. 24**), indicating that these dynamic structures may play important roles in EV-D68, probably in translation initiation. Using the PARIS2-derived structure models as guides, we analyzed structure conservation in 3 additional major enterovirus species, AC, where large numbers of whole genome sequences are available (**Fig. 6f, Supplementary Fig. 25-28**). Alternative conformations V-a1, V-a2 and VI-a showed comparable conservation to some of the previously proposed domains (II, III and V). Together, the combined PARIS2 analysis and phylogenetic analysis revealed a dynamic model of the IRES structure, setting the stage for further functional studies.

## DISCUSSION

The systematically reinvented PARIS2 method is highly efficient and sensitive, overcoming many of the fundamental bottlenecks in current crosslink-ligation based methods for RNA structure/interaction analysis. In particular, we report the identification of the high solubility psoralen derivative amotosalen as a superior crosslinker, compared to the commonly used AMT. We discover abnormally higher hydrophobicity in crosslinked RNA that renders the classical AGPC RNA extraction method inefficient; and develop a new method capable of complete RNA recovery that is generally applicable to crosslinking studies. The full recovery of intact crosslinked RNA enables targeted analysis of structures and interactions, as demonstrated in two applications on cellular and viral RNAs. The newly developed DD2D gel method is robust and isolates crosslinked RNA fragments with high purity, outperforming alternative approaches. To our knowledge, the TNA and 2D gels are the only methods to completely and specifically recover total and crosslinked RNA, respectively. Our in-depth analysis of the photochemical damages in RNA during both the crosslinking and reversal steps have led to new understanding of these processes. Furthermore, we introduced new chemical and enzymatic approaches that significantly improved the prevention and bypass of these damages, solving long-standing problems in the photochemistry field.

In addition to dramatically improving the PARIS method, the newly developed photochemical and enzymatic approaches are generally applicable in molecular biology. For example, the TNA extraction and DD2D gel system are generally applicable to all types of crosslinkers. The prevention and bypass of photochemical damage will be useful in many RNA experiments, since UV irradiation is a commonly used technique. For example, UV crosslinking of RNA-protein interactions also result in damages that reduce cDNA yield and confound the subsequent analysis of crosslink sites^33^.

Despite these major improvements, there are still several steps that will benefit from further optimizations. For example, faster reacting crosslinkers will enable the analysis of more dynamic structures in vivo. The proximity ligation only produces ~10% gapped reads; more efficient ligation will greatly increase the percentage of useful reads and the sensitivity. Further improvement of the prevention, repair and/or bypass of photochemical damages may provide additional benefits, including even higher yield and higher percentages of gapped reads.

The surprising discovery of increased RNA hydrophobicity after crosslinking is reminiscent of crosslinked RNA-protein complexes that partition to the interphase^34–37^, however, it represents a different mechanism. Here the RNA structure itself, coupled with the low pH, seems to be the determinant of hydrophobicity, in contrast to RNA-protein crosslinks where the nonpolar amino acid residues control the hydrophobic behavior. The abnormal in vitro phase partition may be relevant to the role of RNA in phase separation in vivo, even though they occur at different physicochemical environments^38, 39^.

Using PARIS2 and an improved analysis pipeline, we found that the gapped reads report ribosome small subunit and spliceosomal snRNPs across the transcriptome. The bias towards uridines in psoralen crosslinking may confound analysis of binding sites, however, it is unlikely to be critical given the near uniform distribution of the uridines in most mRNAs. Alternatively, a uridine-abundance-based correction can be applied to obtain unbiased measurement of SSU/snRNP binding sites. The simultaneous measurement of mRNA secondary structure and translation status in one experiment will make it possible to directly analyze the impact of RNA structures on translation. The identification of snRNP binding sites and mRNA structures on nascent RNAs will also enable the analysis of structural basis of splicing regulation.

Based on the improved PARIS2, we determined the in vivo structure of the EV-D68 RNA genome, revealing a complex global architecture and dynamic conformations, especially in the IRES that is critical for translation initiation. IRES elements in picornaviruses are particularly fascinating given their rapid evolution and great diversity among the different species. Despite over three decades of research on IRES, our understanding of their in vivo dynamics remains limited. Others have found that certain host factors can induce minor conformations changes in IRES structures^40^. The dynamic conformations are limited to domains V and VI, which bind the major translation initiation factors, including eIF3, 4B, 4G and PTBP^32^. The critical location and evolutionary conservation suggest yet unknown functions of these conformations. We propose that the alternative conformations may represent different stages in the life cycle, such as translation, replication and packaging, or different stages in translation initiation. Furthermore, the transitions among the conformations may act structural switches among the stages. The high efficiency and low cost of the re-invented PARIS2 method will enable highly multiplexed analysis of viral RNA structures and interactions in various viral strains, physiological and pharmacological conditions, and stages of life cycles. Together, PARIS2 will enable RNA structurome and interactome analysis in increasingly more challenging biological systems and enable functional and mechanistic investigations of RNA-centric regulations.

## Acknowledgements

This work was supported by NIH. R00HG009662, and startup fund from USC to Z.L. Computation for the work described in this paper was supported by the University of Southern California’s Center for High-Performance Computing (https://hpcc.usc.edu). We also acknowledge the USC Research Center for Liver Disease (P30DK48522) and Norris Comprehensive Cancer Center (P30CA014089) for their support of our research.

## Author contributions

Z.L conceived this project and designed the overall PARIS2 strategy. W.A.V. synthesized amotosalen. M.Z., C.Y. and Z.L. developed the method. M.L. R.V.D., W.H.L. M.L.C. and J.-F.C. participated in method optimizations and writing. K.L. and J.B. performed the EV-D68 studies. M.Z., K.L., J.B. and Z.L. performed the analysis. RVD, Z.L. wrote the manuscript with input from all authors.

## References

1. Cech, T.R. & Steitz, J.A. The noncoding RNA revolution-trashing old rules to forge new ones. Cell 157, 77–94 (2014).

2. Lu, Z. & Chang, H.Y. The RNA Base-Pairing Problem and Base-Pairing Solutions. Cold Spring Harb Perspect Biol 10 (2018).

3. Lu, Z., Carter, A.C. & Chang, H.Y. Mechanistic insights in X-chromosome inactivation. Philos Trans R Soc Lond B Biol Sci 372 (2017).

4. Ziv, O. et al. COMRADES determines in vivo RNA structures and interactions. Nat Methods 15, 785–788 (2018).

5. Sharma, E., Sterne-Weiler, T., O‘Hanlon, D. & Blencowe, B.J. Global Mapping of Human RNA-RNA Interactions. Mol Cell 62, 618–626 (2016).

6. Aw, J.G. et al. In Vivo Mapping of Eukaryotic RNA Interactomes Reveals Principles of Higher-Order Organization and Regulation. Mol Cell 62, 603–617 (2016).

7. Kladwang, W., Hum, J. & Das, R. Ultraviolet shadowing of RNA can cause significant chemical damage in seconds. Sci Rep 2, 517 (2012).

8. Chomczynski, P. & Sacchi, N. Single-step method of RNA isolation by acid guanidinium thiocyanate-phenol-chloroform extraction. Anal Biochem 162, 156–159 (1987).

9. Lu, Z. et al. RNA Duplex Map in Living Cells Reveals Higher-Order Transcriptome Structure. Cell 165, 1267–1279 (2016).

10. Calvet, J.P. & Pederson, T. Heterogeneous nuclear RNA double-stranded regions probed in living HeLa cells by crosslinking with the psoralen derivative aminomethyltrioxsalen. Proceedings of the National Academy of Sciences of the United States of America 76, 755–759 (1979).

11. Lin, L. et al. Photochemical inactivation of viruses and bacteria in platelet concentrates by use of a novel psoralen and long-wavelength ultraviolet light. Transfusion 37, 423–435 (1997).

12. Thompson, J.F. & Hearst, J.E. Structure of E. coli 16S RNA elucidated by psoralen crosslinking. Cell 32, 1355–1365 (1983).

13. Banyasz, A. et al. Electronic excited states responsible for dimer formation upon UV absorption directly by thymine strands: joint experimental and theoretical study. J Am Chem Soc 134, 14834–14845 (2012).

14. Merriam, V. & Gordon, M.P. Pyrimidine dimer formation in ultraviolet irradiated TMV-RNA. Photochem Photobiol 6, 309–319 (1967).

15. Beukers, R. The effect of proflavine on U.V.-induced dimerization of thymine in DNA. Photochem Photobiol 4, 935–937 (1965).

16. Pathak, M.A. & Fitzpatrick, T.B. The evolution of photochemotherapy with psoralens and UVA (PUVA): 2000 BC to 1992 AD. J Photochem Photobiol B 14, 3–22 (1992).

17. Besaratinia, A., Kim, S.I., Bates, S.E. & Pfeifer, G.P. Riboflavin activated by ultraviolet A1 irradiation induces oxidative DNA damage-mediated mutations inhibited by vitamin C. Proc Natl Acad Sci U S A 104, 5953–5958 (2007).

18. Bleeke, T., Zhang, H., Madamanchi, N., Patterson, C. & Faber, J.E. Catecholamine-induced vascular wall growth is dependent on generation of reactive oxygen species. Circ Res 94, 37–45 (2004).

19. Alenko, A., Fleming, A.M. & Burrows, C.J. Reverse Transcription Past Products of Guanine Oxidation in RNA Leads to Insertion of A and C opposite 8-Oxo-7,8-dihydroguanine and A and G opposite 5-Guanidinohydantoin and Spiroiminodihydantoin Diastereomers. Biochemistry 56, 5053–5064 (2017).

20. Furge, L.L. & Guengerich, F.P. Analysis of nucleotide insertion and extension at 8-oxo-7,8-dihydroguanine by replicative T7 polymerase exo-and human immunodeficiency virus-1 reverse transcriptase using steady-state and pre-steady-state kinetics. Biochemistry 36, 6475–6487 (1997).

21. Pisarev, A.V., Kolupaeva, V.G., Yusupov, M.M., Hellen, C.U. & Pestova, T.V. Ribosomal position and contacts of mRNA in eukaryotic translation initiation complexes. EMBO J 27, 1609–1621 (2008).

22. Archer, S.K., Shirokikh, N.E., Beilharz, T.H. & Preiss, T. Dynamics of ribosome scanning and recycling revealed by translation complex profiling. Nature 535, 570–574 (2016).

23. Ingolia, N.T., Ghaemmaghami, S., Newman, J.R. & Weissman, J.S. Genome-wide analysis in vivo of translation with nucleotide resolution using ribosome profiling. Science 324, 218–223 (2009).

24. Lu, Z., Guan, X., Schmidt, C.A. & Matera, A.G. RIP-seq analysis of eukaryotic Sm proteins identifies three major categories of Sm-containing ribonucleoproteins. Genome Biol 15, R7 (2014).

25. Ntini, E. et al. Polyadenylation site-induced decay of upstream transcripts enforces promoter directionality. Nat Struct Mol Biol 20, 923–928 (2013).

26. Friend, K., Lovejoy, A.F. & Steitz, J.A. U2 snRNP binds intronless histone pre-mRNAs to facilitate U7-snRNP-dependent 3' end formation. Mol Cell 28, 240–252 (2007).

27. Kaida, D. et al. U1 snRNP protects pre-mRNAs from premature cleavage and polyadenylation. Nature 468, 664–668 (2010).

28. Almada, A.E., Wu, X., Kriz, A.J., Burge, C.B. & Sharp, P.A. Promoter directionality is controlled by U1 snRNP and polyadenylation signals. Nature 499, 360–363 (2013).

29. Cassidy, H., Poelman, R., Knoester, M., Van Leer-Buter, C.C. & Niesters, H.G.M. Enterovirus D68-The New Polio? Front Microbiol 9, 2677 (2018).

30. Lee, K.M., Chen, C.J. & Shih, S.R. Regulation Mechanisms of Viral IRES-Driven Translation. Trends Microbiol 25, 546–561 (2017).

31. Steil, B.P. & Barton, D.J. Cis-active RNA elements (CREs) and picornavirus RNA replication. Virus Res 139, 240–252 (2009).

32. Martínez-Salas, E. The impact of RNA structure on picornavirus IRES activity. Trends Microbiol 16, 230–237 (2008).

33. Lee, F.C.Y. & Ule, J. Advances in CLIP Technologies for Studies of Protein-RNA Interactions. Mol Cell 69, 354–369 (2018).

34. Urdaneta, E.C. et al. Purification of cross-linked RNA-protein complexes by phenol-toluol extraction. Nat Commun 10, 990 (2019).

35. Queiroz, R.M.L. et al. Comprehensive identification of RNA-protein interactions in any organism using orthogonal organic phase separation (OOPS). Nat Biotechnol 37, 169–178 (2019).

36. Trendel, J. et al. The Human RNA-Binding Proteome and Its Dynamics during Translational Arrest. Cell 176, 391–403.e319 (2019).

37. Shchepachev, V. et al. Defining the RNA interactome by total RNA-associated protein purification. Mol Syst Biol 15, e8689 (2019).

38. Jain, A. & Vale, R.D. RNA phase transitions in repeat expansion disorders. Nature 546, 243–247 (2017).

39. Zhang, H. et al. RNA Controls PolyQ Protein Phase Transitions. Mol Cell 60, 220–230 (2015).

40. Tolbert, M. et al. HnRNP A1 Alters the Structure of a Conserved Enterovirus IRES Domain to Stimulate Viral Translation. J Mol Biol 429, 2841–2858 (2017).

