## Supplementary Information for "Optimized photochemistry and enzymology enable efficient analysis of RNA structures and interactions in cells and virus infections"

In these supplementary notes, we provide a historical account of the technical challenges in the field of RNA duplex analysis using photochemical crosslinkers, physical and chemical mechanisms of these problems, and detailed descriptions of the optimizations. Much of the valuable information regarding RNA photochemistry was difficult to find in the literature, and most of the new results are not included in the main text due to limited space. In addition to the optimal conditions that we discovered, we also present negative data on alternative approaches that we have attempted, in the hope that these data will be useful for other researchers who are interested in further optimizations. Some of the studies, although not useful for improving PARIS, revealed fundamental principles of RNA physics and chemistry.

**List of supplementary notes.**

| | |
|---|---|
| Supplementary Note 1. | Phase partition of crosslinked RNA and development of the TNA method |
| Supplementary Note 2. | Optimization of RNA fragmentation |
| Supplementary Note 3. | Developing the DD2D gel separation method |
| Supplementary Note 4. | Protection of RNA against UVC damage |
| Supplementary Note 5. | Bypass of PUVA induced oxidative damage on RNA |

**Supplemental Note 1. Phase partition of crosslinked RNA and development of the TNA method**

<u>1. Background about the AGPC method.</u>

The classical acid guanidinium thiocyanate-phenol-chloroform (AGPC) method has been considered a gold standard in RNA extraction [1] (cited more than 70,000 times by April 2020). This method uses guanidinium thiocyanate (GuSCN), one of the strongest chaotropic agent, and acidic phenol, a denaturant, to disrupt biological materials and stabilize RNA. After adding chloroform, cellular components partition to the two liquid phases according to polarity/hydrophobicity: RNA in the upper aqueous phase, DNA in the interphase, and proteins and lipids in the interphase and lower organic phase (**Fig. 3a-b**). In contrary, the neutral phenol-chloroform isoamyl alcohol (PCI) method uses near neutral pH (~8) to partition DNA to the upper aqueous phase.

The original AGPC method used a mixture of solution D (4M GuSCN, 25mM sodium citrate, pH 7; 0.5% sarcosyl, 0.1M 2-mercaptoethanol (RNase inhibitor)), 0.2M sodium acetate pH 4, and water saturated phenol at 1:0.1:1 ratio [1]. Later, the protocol was modified so that all components are combined in a monophasic solution: 0.8M GuSCN (0.5-2M range), 0.4M ammonium SCN (0.1-0.6M range), 0.1M sodium acetate, pH 5 (4-6 range), 38% phenol (30-50 range), 5% glycerol (3-10% range) [2]. The 5% glycerol was used to blend the components of different polarity into one phase. This method has been commercialized in several kits, such as TRIzol, QIAzol and TRI reagent, and more recently, RNAzol, that were widely used in RNA research. The TRIzol LS (liquid sample) reagent has a proprietary composition, but it likely contains higher concentrations of these components, especially GuSCN, phenol and sodium acetate, so that lower volumes of TRIzol are used for liquid samples. The AGPC method allows quantitative recovery of pure RNA without any degradation of this labile molecule. Almost all cellular RNA molecules >20nt can be completely recovered, with one exception. The Kim lab reported that short structured miRNAs with low GC content are selectively lost during TRIzol extraction, leading to artifacts in miRNA quantification [3].

The theoretical basis of the phase partition of RNA and DNA in the aqueous-organic systems at different pH remains poorly understood. No quantitative analysis of the hydrophobicity/polarity has been published, to the best of our knowledge. Some researchers suggested that the lower pH (4-5) neutralizes the negative charge on DNA, which caused the DNA to be more hydrophobic (see brief overview of the method and its history by Paul Zumbo, "*Phenol-chloroform Extraction*"), but we believe this is not true. The phosphate backbone has a lowest pKa around 1-2, much lower than the pH range 4-5 used for RNA extraction, and thus should remain negatively charged (**Fig. 3a**).

<u>2. Psoralen crosslinked RNA partitions to the interphase in TRIzol extraction</u>

When we used TRIzol to extract RNA from AMT crosslinked cells, we noticed that the yield reduced as crosslinking strength increased (**Supplementary Fig. 3a**). Crosslinking cells with 0.1mg/ml AMT for 30min reduced yield to ~60%, and produced insoluble material in the TRIzol lysate from cells. The insoluble material promoted emulsion formation after addition of chloroform (see Fig. 2 from [4]), and then partitioned to the interphase after phase separation. Higher concentrations of psoralen resulted in higher crosslinking efficiency and even lower recovery of RNA from the aqueous phase (**Fig. 2b-c**). Initially we suspected that RNA was crosslinked to proteins and selectively trapped in the interphase. To test this hypothesis, we performed proteinase K (PK) treatment prior to TRIzol extraction. PK treatment consistently but modestly increased yield; there was still significant loss of RNA (**Supplementary Fig. 3a** and Supplemental data from [5]). This was consistent with previous reports that psoralen can crosslink nucleic acids to proteins, although at much lower efficiency than between nucleic acid strands [6, 7].

3. S1/PK digestion as a temporary solution to recover crosslinked RNA.

We used higher concentrations of AMT and amotosalen for PARIS experiments, which lead to even lower RNA recovery, down to ~20-30% (**Fig. 3d** in this paper and **Fig. S1 from** [5]). To improve yield, we used S1 nuclease, which is active on DNA and RNA in cell lysates even under extremely highly denaturing conditions, such as 9M urea and 0.1% SDS [8]. Lysate digestion with S1 nuclease in 4M urea and 0.1% SDS led to higher yield of crosslinked RNA, especially in comparison with non-crosslinked samples (see **Fig. S1** from [5]). Together, the S1 nuclease and PK treatment led to sufficient RNA yield and was successfully used in our initial PARIS method, but there were still two problems. First, we were not sure if all crosslinked RNA has been recovered. Second, RNA fragmentation prior to extraction made it difficult to perform targeted RNA enrichment, for example using oligo(dT) for mRNAs, or RNA-specific antisense oligos.

4. Psoralen crosslinking increases RNA hydrophobicity.

The earlier observation that PK treatment plus TRIzol purification did not completely recover all crosslinked RNA suggest that crosslinked RNA itself became more hydrophobic. To confirm that AMT crosslinking alters RNA hydrophobicity, we purified normal total RNA from cells for crosslinking in vitro, and then directly precipitated RNA from the solution or used the standard TRIzol method to purify RNA (**Fig. 3c**). Non-crosslinked RNA was extracted efficiently in both methods (slightly lower yield from TRIzol, compared to direct ethanol precipitation, because of incomplete recovery of the aqueous phase), while crosslinked RNA was lost from standard TRIzol extraction, compared to direct ethanol precipitation. Non-crosslinked RNA appears as sharp peaks in the small RNA (50-300nt), 18S and 28S peaks (first 3 panels), while the crosslinked RNA shows as a broad smear spanning the entire profile (4$^{th}$ panel). The electrophoresis profile showed clear separation of noncrosslinked and crosslinked RNA between the aqueous and inter+organic phases (last two panels). While stronger crosslinking lead to gradual loss of RNA from the aqueous phase, more RNA accumulates in the interphase (**Fig. 3d**). These results suggest that crosslinked RNA is more hydrophobic, and partitions to the interphase, and is therefore lost during TRIzol extraction. Given that proteins were removed prior to RNA purification, the increased hydrophobicity was likely due to crosslinked RNA itself.

To further confirm that crosslinking increased RNA hydrophobicity, we tested addition of formamide in the standard TRIzol purification. Adding formamide greatly increased the partition of RNA to the aqueous phase (**Supplementary Fig. 3b**). Together, these results proved that crosslinking increases RNA hydrophobicity, making it similar to DNA, which normally partitions to the interphase during standard TRIzol extraction (see **Fig. 3b** for a summary).

5. Most crosslinked RNA is in the interphase of TRIzol-chloroform mixture

In order to confirm that most crosslinked RNA is in the interphase, we digested RNA from the aqueous and inter+organic phase using RNase III and ran a DD2D gel to separate crosslinked and noncrosslinked RNA (**Supplementary Fig. 3c-d**). From the same amount of RNase III digested RNA, we recovered similar amounts of crosslinked RNA from the aqueous phase and inter+organic phase (**Supplementary Fig. 3e**, both the columns "stuck in 1D" and "2D upper diagonal" are crosslinked RNA). Given that there is significantly more RNA in the interphase than the aqueous phase (**Fig. 3d**), these results suggest that in standard TRIzol purification, most crosslinked RNA are stuck in the interphase. For example, assuming that 20% RNA was recovered in the aqueous phase in standard TRIzol extraction, then more than 80% crosslinked RNA is lost. More importantly, larger RNAs are preferentially lost, leading to bias in the results.

6. Smaller crosslinked RNA do not partition to the interphase.

In summary, our studies showed that, first, psoralen crosslinks RNA to proteins, in addition to among nucleic acids, and second, crosslinking increases RNA hydrophobicity and causes it to re-partition to the interphase. To test whether we can use the difference in the hydrophobicity between crosslinked and noncrosslinked RNA to enrich for crosslinked RNA fragments during TRIzol extraction, we crosslinked RNA in vitro and then digested RNA using S1 nuclease and RNase III to make most RNA < 150nt. After the two digestions, we added TRIzol to the solution and then added chloroform, however, no clear interphase was observed, suggesting that the small crosslinked RNA fragments did not partition to the interphase. Furthermore, the precipitate from the inter+organic phase was not soluble in water and the Nanodrop profile shows major peaks at 230nm and 270nm, clearly different from RNA (260nm). Together these results showed that large crosslinked RNA, but not small crosslinked RNA fragments partition to the interphase, consistent with previous results (**Fig. 3c**). Therefore, we cannot selectively purify small crosslinked RNA fragments from the interphase using TRIzol.

7. Developing the new TNA method.

In the studies described above, we discovered that PK was needed to improve recovery of crosslinked RNA, and crosslinking made RNA more hydrophobic. Based on these observations, we decided to develop a new method to efficiently purify crosslinked RNA. First, we tested direct precipitation of total nucleic acids (TNA) from PK digested lysates from crosslinked cells. To obtain intact RNA, we searched for lysis conditions that would effectively inhibit all nucleases but at the same time allow efficient PK digestion. The TRIzol solution inhibits proteinase K, so it is not appropriate for cell lysis. Guanidine thiocyanate (GuSCN) is one of the strongest chaotropic agent, and at above 4M can denature most proteins, including nucleases [1, 9]. Therefore, we first lysed cells in 4M GuSCN (1 volume cell pellet + 2 volumes 6M GuSCN, pH ~5.3), and this usually leads to a clear solution, even for the crosslinked samples. Then we diluted the lysate with phosphate buffered saline (PBS) to 1M GuSCN and added EDTA to chelate divalent cations and performed PK treatment on cell lysates at 37C for 1 hour. The PBS dilution of GuSCN solution resulted in some insoluble material, which was then cleared by PK, suggesting most proteins were mostly digested. However, addition of 1 volume isopropanol (relative to the sum of cell pellet, PBS, GuSCN and EDTA.) lead to precipitates that could not be dissolved in water (**Supplementary Fig.**

**3f,** blue bars). Surprisingly, adding TRIzol before isopropanol precipitation lead to precipitates that could be dissolved, suggesting that the TRIzol components, probably phenol, help keep the residual proteins in solution during isopropanol precipitation (**Supplementary Fig. 3f**).

To confirm that it was phenol that kept proteins in solution, we compared TRIzol and phenol during isopropanol precipitation (**Supplementary Fig. 3g**). Indeed, adding phenol and isopropanol was sufficient to produce RNA pellets that could be dissolved in water. Both ethanol and isopropanol can be used for precipitation. We preferred isopropanol in precipitation because less volume is needed. The disadvantage is that isopropanol is less volatile, and salts are less soluble in isopropanol, which may result in excess salt precipitation (Paul Zumbo, "*Phenol-chloroform Extraction*"). The salt precipitation problem can be resolved by extended 70% ethanol washes or reprecipitation.

8. Apparent lower yield of crosslinked RNA due to hypochromicity of RNA structures.

While testing direct ethanol precipitation of RNA from in vitro crosslinking, and the TNA method for nucleic acid extraction from crosslinked cells, we noticed that consistently less nucleic acids were recovered from the crosslinked samples (usually ~60-80%) and the A260/A280 ratio was lower (**Supplementary Fig. 3g**). We are certain that this is not caused by nucleic acid loss, because in vitro crosslinked pure RNA samples can be completely recovered. This was probably because crosslinking forced the formation of duplexes that absorb less UV light (1 OD260 Unit = 50ug/ml for dsDNA, or 40ug/ml for ssRNA, or 33ug/ml for ssDNA, or 20ug/ml for ssOligo) [10]. In fact, this higher percentage of double stranded regions in RNA, which is more hydrophobic, is consistent with our observation that crosslinked RNA is more hydrophobic, partitions to the interphase, like DNA, during standard TRIzol extraction. In addition, the selective crosslinking of uridines would also lead to different A260/A280 ratios as adenine and uracil are the major components in nucleic acids that absorbs UV light (see reference: Thermo Scientific T042-Technical Bulletin).

9. Removing DNA from total nucleic acids to recover all RNA.

Given that the TNA extraction protocol recovered total nucleic acids, next we performed DNase treatment to purify crosslinked RNA. Using Turbo DNase, we were able to digest away most DNA (**Fig. 3f-g**), and recover RNA that makes up ~40-60% of TNA. In practical applications, another round of DNase treatment can be performed if necessary, for example, after antisense enrichment of certain RNA populations. This second step of DNase treatment will be more efficient after antisense enrichment, given that the antisense enrichment will also reduce DNA contamination. Therefore, DNA contamination is not a concern in PARIS and similar experiments. In particular, real-time quantitative PCR with or without reverse transcription, a commonly used method, can be employed to assess the amount of DNA contamination. In our experience, one round of DNase treatment reduced DNA contamination to undetectable levels based on qPCR (Ct values always >40). Together, the above results demonstrated that we have developed a new method that is superior to the classical AGPC method in extracting crosslinked RNA.

10. Chlorambucil and carmustine crosslinking increases RNA hydrophobicity.

Is the crosslink-induced hydrophobicity a general property for RNA and different types of crosslinkers? To answer this question, we performed RNA crosslinking using two chemotherapy drugs chlorambucil (CHL) and carmustine (BCNU), which react with nucleic acids via different mechanisms (**Supplementary Figs. 4 and 5**).

Chlorambucil (CHL) is a nitrogen mustard that acts as a bifunctional alkylating agent and is used as a pharmaceutical agent, especially in chemotherapy (IARC 1987) (**Supplementary Fig. 4a**). During crosslinking, the aziridinium rings formed by intramolecular displacement of the chloride by amine nitrogen, alkylates DNA once it is attacked by the N-7 nucleophilic center on the guanine base. Then a second consecutive attack after the displacement of the second chlorine results in the formation of interstrand cross-links (ISC) (**Supplementary Fig. 4b**). The alkylation rates are limited by the rate of aziridinium ions' formation, and DNA ISC induced by CHL is formed at a specific site, 5'-GGC sequence (an 1,3 cross-link, G1-G3) by an DNA strand cleavage assay [11]. Using a synthetic DNA oligo duplex, we tested various concentrations and incubation time, and observed significant crosslinking after 3 hours (**Supplementary Fig. 4c-d**). Then we performed in vitro crosslinking of purified total RNA using CHL and observed strong smear that spans beyond the 28S rRNA peak, suggesting successful in vitro crosslinking (**Supplementary Fig. 4e**). We then precipitated the crosslinked RNA, digested RNA with RNase III and separated RNA using the DD2D gel system (**Supplementary Fig. 4f**). Crosslinked RNA was observed above the diagonal in the 2D gel. These results establish CHL as a strong RNA crosslinker.

To test whether the CHL crosslinking made RNA more hydrophobic, we crosslinked RNA, purified RNA using either direct ethanol precipitation, or the standard TRIzol method, where we extracted RNA from both the aqueous and inter+organic phases (**Supplementary Fig. 4g**). While non-crosslinked RNA was extracted from the aqueous phase efficiently (first three panels), crosslinked large RNA (e.g. 18S and 28S) partitioned to the interphase (bottom 3 panels), similar to psoralen crosslinked RNA (**Fig. 3c**). These results demonstrated that CHL crosslinking increased RNA hydrophobicity, similar to AMT and amotosalen crosslinking.

Carmustine (BCNU) is another category of commonly used chemotherapy drug, which can crosslink DNA in cells and is used for multiple cancers (**Supplementary Fig. 5a**). It belongs to chloroethylnitrosoureas (CENU), a new kind of alkylating agent which was developed later than nitrogen mustards. CENU exert cytotoxicity by inducing DNA interstrand cross-links (ICLs) between guanine and the complimentary cytosine, namely dG–dC crosslink (**Supplementary Fig. 5b)**. The formation of a covalent connection between two DNA strands requires 2 successive reactions: (a) an alkylation or other modification of one strand; (b) a reaction of the modified strand with the complementary DNA strand (Kurt W. Kohn, 1977). We crosslinked purified total RNA, performed RNase III digestion and DD2D gel analysis. Crosslinked RNA fragments above the diagonal indicated that the crosslinking worked well (**Supplementary Fig. 5c-d)**. The crosslinking process induces appreciable degradation (**Supplementary Fig. 5e**), however, this did not affect the experiments. Similar to the experiments on CHL and psoralens, we tested the partition of crosslinked RNA in the two phases during TRIzol extraction (**Supplementary Fig. 5f**). Despite the degradation, it was clear that larger crosslinked fragments are partitioned to the inter and organic phase (bottom panels).

Taken together, the analysis of chlorambucil and carmustine, two different categories of nucleic acid crosslinkers, confirmed that crosslinked RNA is more hydrophobic, the TNA method is generally applicable to crosslinked RNA, and the crosslinked fragments can be isolated using the DD2D gel system. These results showed that the crosslinking-induced hydrophobicity is not unique to psoralens, and is likely to be a general property of RNA. Although both chemotherapy drugs can crosslink RNA, they are not easily applicable to the analysis of RNA structures and interactions, because the crosslinks are not reversible.

11. Comparison among AGPC (TRIzol), TNA and silica-gel (RNeasy) methods

As an alternative to the phase partition approach in the AGPC method, silica gels (e.g. RNeasy kit from Qiagen and many other column-based nucleic aic purification kits) have been used in the isolation of RNA and DNA from cell lysates. Nucleic acids absorb onto the silica in the presence of high concentrations of salt and chaotropic agents, and dissociate from silica at lower concentrations [12]. This method was recently used in another study to purify psoralen crosslinked RNA [13]. To compare the TNA method with the silica gel method, we first crosslinked pure total RNA with psoralen, then purified RNA with either direct ethanol precipitation or the RNeasy kit (**Supplementary Fig. 6a**). Compared to the TNA method, silica gel based method results in partial loss of crosslinked RNA (**Supplementary Fig. 6b**). Shorter RNAs, such as in the range of 50-300nt, are lost in the flow-through (**Supplementary Fig. 6c**).

To test recovery of RNA from psoralen crosslinked cells, we purified RNA using standard TRIzol, TNA and the RNeasy kit (**Supplementary Fig. 6d**). The TNA method retrieved all RNA, while TRIzol and RNeasy kit retrieved much lower amount RNA, and the retrieved RNA are significantly biased towards the lower end of size distribution (**Supplementary Fig. 6e-f**). The difference in performance was most dramatic at the highest crosslinker concentrations. For example, from cells crosslinked with 5mg/ml amotosalen, standard TRIzol and RNeasy methods lost more than 80% of the RNA. Together, these two sets of experiments showed that TNA is the best method for purifying crosslinked RNA. To the best of our knowledge, it is the only method capable of complete recovery of crosslinked RNA

12. Summary and discussion

While this study was in progress, several groups reported a new method for the isolation of crosslinked RNA-protein complexes based on their hydrophobicity [14-17]. The partition of crosslinked RNA-protein complexes to the interphase of an aqueous-organic mixture depends on the nonpolar amino acid residues in the protein part. This mechanism is different from the crosslinking/structure-induced hydrophobicity of RNA by itself.

We note that, in all previously published studies that employ psoralen crosslinking, it is very likely that most of the highly crosslinked large RNA were lost during purification [18]. Most previous in vitro and in vivo psoralen crosslinking protocols use way less AMT and much shorter time, which is why this abnormal behavior of crosslinked RNA has never been noticed. Earlier studies often focused on highly abundant RNAs, like rRNAs, snRNAs and snoRNAs, so the reduced sensitivity was not a problem. None of the recent studies that employ high throughput sequencing noticed or investigated this problem either [13, 19-21]. Some of the previous studies used PK to digest the samples after crosslinking, but only partially resolved the problem of low RNA recovery [22].

In summary, we made a surprising discovery that crosslinked RNA behaves differently from non-crosslinked RNA. Crosslinking increases RNA hydrophobicity, leading to its repartition to the interphase during standard TRIzol extraction. Based on this discovery, we have developed a new method to extract crosslinked RNA from cells. Our method represents a major breakthrough in solving this bottleneck problem and will greatly facilitate future studies using psoralen and other crosslinkers.

**Supplementary Note 2. Optimization of RNA fragmentation**

1. Introduction to the RNA fragmentation problem

Fragmentation of crosslinked RNA to small pieces is necessary for establishing secondary structures or RNA-RNA interactions at near base pair resolution. Various types of metal ion buffers and nucleases have been described, including RNase III (commercial name ShortCut from NEB, which produces dsRNA fragments above 18bp) [23], S1 nuclease, RNase A/T1, RNase I and magnesium ($Mg^{2+}$) [18]. These approaches differ in their cost, sensitivity to experimental conditions, product size distribution, product terminal chemistry (phosphate or hydroxyl on the 5' or 3' ends) etc. Initially, we tested the commonly used RNase T1 but found that it is difficult to control the digestion to obtain a narrow distribution of the fragment size (**Supplementary Fig. 7a**). RNase A/T1, RNase I, and divalent cations all produce 5' OH and 3'phosphate that require additional repair to make 5'phosphate and 3'hydroxyl for the next step of proximity ligation and adapter ligation.

2. Earlier optimizations of RNA fragmentation

We have chosen S1 nuclease and RNase III for several reasons [5]. First, S1 is active under highly denaturing conditions, such as 9M urea and 0.1% SDS. Second, S1 nuclease and RNase III both produces 5' phosphate and 3' hydroxyl (OH) that can be directly used for ligation and library preparation without further repair. Third, *E. coli* RNase III, when used with $Mn^{2+}$, cleaves dsRNA while protecting the products such that minimal RNA fragments are around 18-25bp [23], a suitable size window for our structure analysis (small enough for accurate modeling of base pairing, and big enough for mapping to the genome). To test whether S1 nuclease alone is sufficient for the fragmentation, we tested digestion for various time, but could not bring most of the RNA fragments to below 100nt even after prolonged incubation (**Supplementary Fig. 7b**).

3. Optimization of RNase III fragmentation of RNA.

While using the Native-Denatured 2D (ND2D) gel system to select crosslinked RNA fragments, we noticed that cross-linked and RNase III digested RNA tend to be bigger in apparent size than noncrosslinked RNA in the first dimension gel (see Fig. S1 from Lu et al., 2016). We reasoned that fragmentation conditions that amplify this difference could be used to isolate crosslinked RNA in a 1D gel alone, which would greatly reduce the time and effort of PARIS experiments and reduce loss of RNA during gel extraction. In fact, earlier studies have shown that *E. coli* RNase III does not only cleave dsRNA [24]. At lower ionic strength, it cleaves both ssRNA and dsRNA efficiently to small sizes (see Figure 3 in [24]. We found that the low ionic strength buffers did produce shorter fragments from the noncrosslinked RNA, mostly less than 40nt, while the digestion of crosslinked RNA gave rise to larger fragments (**Supplementary Fig. 7c,** significant tail above 40nt, indicated by the arrows). This result is consistent with previous studies of the wildtype E. coli RNase III [24]. However, when we run the 2D gels to select crosslinked RNA, the yield was consistently lower than before (~0.1%, compared to previous yield of 0.25-0.5%)[5], suggesting that the denatured 1D gel alone was insufficient for retrieving all crosslinked RNA fragments.

Given that RNase III can cleave both single and double stranded RNA, we sought to determine whether RNase III alone was sufficient for RNA fragmentation for the 2D gel separation. Varying the enzyme amount and incubation time, we found that the kinetics of the reaction determines the size distribution (**Supplementary Fig. 7d**). A short reaction time with high amounts of RNase III produced fragments that mostly lie in the range of 30-100nt (indicated by arrows), perfectly suited for 2D gel separation and library preparation. This kinetic effect is due to the tight binding of RNase III to the products, which inhibits further digestion of the short fragments [23]. High ratios of enzyme vs. RNA lead to efficient digestion leading to a more uniform size protected by the enzyme, while low ratios lead to both shorter fragments and longer fragments.

4. Summary and discussion

Here we presented a simplified strategy for RNA fragmentation that resulted in a narrow size distribution perfectly suited for RNA structure analysis and many other studies that require fragmented RNA. This method is better than other approaches for several reasons. First the protocol is simple, requiring only one enzyme. Second, the size distribution is narrow. Third, the products have 5' phosphate and 3' OH, suited for direct subsequent ligation reactions in library preparation and other types of enzymatic reactions.

**Supplementary Note 3. Developing the DD2D gel separation method**

1. Overview of the problem of enriching crosslinked RNA fragments

Several strategies have been used for enriching crosslinked RNA in recent high throughput analysis of RNA duplexes, including ND2D gel [5], biotinylated psoralen pull down [13, 21], and RNase R trimming of single stranded RNA [20] (**Supplementary Fig. 8a-c**). It has been shown that monoadducts are the major products in psoralen crosslinking, while only 20-40% of adducts are crosslinks in DNA [25]. Higher concentrations of psoralens are likely to cause higher ratios of monoadducts because less efficient sites are also forced to react with psoralens. We also found that RNA crosslinking is much less efficient, producing much more monoadducts than crosslinks (see details in **Supplementary Note 4** and **Supplementary Fig. 11**). Purification of biotinylated psoralen crosslinked nucleic acids recovers more monoadducts than crosslinks, dramatically reducing the sensitivity of the method (only a small percentage of RNA fragments can be used to produce proximally ligated RNA). This is in contrast to single chemical tagging reactions with target biomolecules, where the biotin handle is good enough for purifying the reacted molecules [26]. RNase R digestion of single stranded RNA is also blocked by the bulky monoadducts. As a result, the ND2D gel is the only method that ensures isolation of pure crosslinked RNA fragments. We calculated the percentage of gapped and chimeric reads in published methods, such as hiCLIP, SPLASH, LIGR-seq and COMRADES [13, 20, 21, 27], and found that PARIS [5], which used the ND2D gel method, consistently outperforms other methods, consistent with the idea that the 2D gel isolation of pure crosslinked RNA is essential for obtaining high percentages of gapped/chimeric reads (**Supplementary Fig. 8d)**.

2. The ND2D method and its problems

While performing the RNase III digestion and denatured gel purification of crosslinked RNA fragments, we noticed that the yield (~0.1%) is much lower than what we have achieved previously (0.23-0.55%) [5] (see Figure S1 in [5]). This low yield is likely due to the strong RNase III digestion that reduced size of certain crosslinked RNA to below 30-40nt. To solve this problem, we tested other nuclease fragmentation methods, including S1 alone, lighter RNase III alone, RNase A and RNase A/T1. While testing these conditions, we noticed that the 12% native and 20% denatured 2D gel cannot separate crosslinked fragments from non-crosslinked below 40-50nt. This observation prompted us to reexamine the theoretical basis of the 2D gel method.

Vigne and Jordan described the first native-denatured 2D (ND2D) gel system to analyze RNA structures [28]. In this pioneering study the authors showed that the second denatured dimension gel will separate an RNA duplex from the first native dimension and thus allow the identification of the two fragments held together by hydrogen bonding (not covalent bound). Zwieb and Brimacombe adapted ND2D gel for the analysis of crosslinked RNA fragments [29] (see diagram in **Supplementary Fig. 8a**). Here the crosslinked fragments run as a tight duplex in the first dimension and then opened up to an "octopus" shape in the second denaturing dimension and therefore ran much slower, separating from the non-crosslinked fragments. Thompson and Hearst applied the ND2D gel to analyze AMT-crosslinked RNA fragments [30]. However, the claim that "the crosslinked hairpin loops actually run slightly faster in the second dimension because their radius decreases somewhat" is likely to be an incomplete statement, because hairpins are likely to run slower upon denaturation. It has been shown that short circular RNAs (e.g. 20nt), which are constrained and base paired, run faster than their linear forms which would be more single-stranded in TBE-urea gel. We also noticed that in native gels, the singled stranded RNA runs much slower than an RNA duplex of the same molecular weight, suggesting that upon denaturation in the second dimension, stem-loops would run slower to match that of an RNA of the same molecular weight that was already single-stranded in the first dimension. This slow-down would mask some crosslinked dsRNA fragments in the second dimension.

3. The DD2D method for separating crosslinking RNA from noncrosslinked.

To solve this problem, we explored the possibility of running denatured-denatured 2D (DD2D) gel, where the gel percentage is lower in the first dimension than the second (**Supplementary Fig. 8b**). The rationale for this design is as follows: structured RNA molecules encounter higher friction in the higher percentage gels, therefore, they run much slower relative to their linear counterparts. In previous experiments, I noticed that the 113nt tricRNAs runs near its real size in a 6% gel, but close to 200nt in a 10% gel [31]. Earlier studies also showed that a 5% denatured – 10% denatured 2D system can be used to separate lariat RNAs from linear RNAs [32]. We tested several combinations of gel percentages in the first and second dimensions and found that the 8%+16% combination produced clear separation of crosslinked RNA from the non-crosslinked (**Supplementary Fig. 8e-g**). The crosslinked RNA forms a smear above the upper diagonal, or gets stuck in the 1D-2D interface. Compared to the ND2D gel, the recovery of crosslinked RNA increased by 50%, therefore increasing the sensitivity of the PARIS method (**Supplementary Tables 1 and 2**).

We found that RNA crosslinked with BCNU and chlorambucil can be easily separated from noncrosslinked fragments (**Supplementary Figs. 4 and 5**). As long as the density of the second dimension is different from the first dimension, the separation should work well. Therefore, combinations different from the 8%+16% gel system shown in this study can be used in other applications where either longer or shorter crosslinked fragments can be studied.

A caveat with this design is that shorter crosslinked RNA duplexes may not run slower in the higher percentage gel. This is based on the observation that while longer circular RNAs actually run slower than their counterpart, shorter circular RNAs run faster than their linear counterparts. Indeed, in our 8% denatured – 16% denatured 2D gels, we cannot see any crosslinked dsRNA below 50nt above the diagonal. Because of this, we need to make sure that the nuclease digestion keeps most RNA fragments above this range. Given that this new system does not separate RNA based on their base pairing status, it is generally applicable to all types of nucleic acid crosslinking studies.

Here is a simplified quantitative model for the DD2D separation of linear and structured nucleic acids.
$f(p) = \exp\{-\mathrm{k}_1 p\}$ for $p \in [p_1, p_2]$, speed dependence on gel concentration p. $k_1$ is a constant within certain range of p.
$g(l) = \mathrm{k}_2/\ln(l)$ for $l \in [l_1, l_2]$, speed dependence on nucleic acid length l. $k_2$ is constant within certain range of l.
$v_{\mathrm{L}}(p,l) = f(p) \cdot g(l)$, speed for linear nucleic acids
$v_{\mathrm{S}}(p,l,s) = f(p) \cdot g(l) \cdot h(p,l,s)$, speed for structured nucleic acids
$f(p)$ is a monotonically decreasing function of gel concentration, or increasing function of gel pore size (Stellwagen and Stellwagen 2009, Effect of the matrix on DNA electrophoretic mobility)
$g(l)$ is a monotonically decreasing function, inversely proportional to the logarithm of nucleic acid length within a certain range, therefore, $v_{\mathrm{L1}}/v_{\mathrm{L2}}$ still has some dependence on nucleic acid length (find original reference) (https://www.thermofisher.com/us/en/home/life-science/cloning/cloning-learning-center/invitrogen-school-of-molecular-biology/na-electrophoresis-education/na-separation-overview.html).
$v_{\mathrm{L}}$ is the migration speed of a linear nucleic acid.
$v_{\mathrm{S}}$ is for the structured, e.g. circular, branched, lariat, crosslinked duplexes, etc.
$v_{\mathrm{L,2D}}/v_{\mathrm{L,1D}} = f(p_{\mathrm{2D}})/f(p_{\mathrm{1D}}) = \exp\{-\mathrm{k}_1(p_{\mathrm{2D}} - p_{\mathrm{1D}})\}$ , relative speed of linear nucleic acid for the same length

$$(v_{\mathrm{S,2D}}/v_{\mathrm{S,1D}})/(v_{\mathrm{L,2D}}/v_{\mathrm{L,1D}}) = h(p_{\mathrm{2D}},l,s)/h(p_{\mathrm{1D}},l,s) \begin{cases} > 1, \text{if } l < l_0 \\ = 1, \text{if } l = l_0 \\ < 1, \text{if } l > l_0 \end{cases}$$ , relative speed of structured vs. linear nucleic acids.

$l_0$ is close to 30 in the case of circular RNAs. For example, we know $h(p,l,s) > 1$ for $l \leq 24$, and < 1 for $l > 37$, see discussion of the circular ssDNA C44, C60, C66 and C70 [33]. The dependence of migration speed on RNA structure s is more difficult to quantify. $h(p_{\mathrm{2D}},l,s)/h(p_{\mathrm{1D}},l,s)$ is a monotonically decreasing function of nucleic acid length and gel concentration, which is why on the higher percentage gel, the impact of gel concentration on structured nucleic acids is larger. The size of the crosslinked RNA fragments is difficult to estimate based on the denatured gel, because they run much slower than a linear RNA of exactly the same size. Earlier studies showed that in a 14% denatured polyacrylamide gel, a 72nt circular ssDNA runs close to 500nt linear ssRNA in size [33].

4. Summary and discussion

Here we showed that a denatured 2D gel with different density between the two dimensions effectively separates crosslinked from noncrosslinked nucleic acids. The DD2D method does not depend on specific structures of the crosslinkers or the RNA fragments, therefore it is generally applicable to all types of nucleic acid crosslinking experiments.

**Supplementary Note 4. Protection of RNA against UVC damage**

1. Introduction to the problem of photochemical damage of RNA

When preparing PARIS sequencing libraries, we noticed that the vast majority of the DNA in the final step of gel selection have very short inserts, many in the range of a few base pairs, even though the original crosslinked RNA fragments that we selected were at least 30-40nts [5]. These results suggest that additional factors besides proximity ligation lead to the small insert size, and we suspect that this is due to RNA damage during to AMT-mediated long-wavelength (UVA, 365nm) photo-crosslinking and subsequent short wavelength (UVC, 254nm) reversal. Psoralen + UVA (PUVA) primarily lead to oxidative damage, while UVC primarily induces the formation of cyclobutane pyrimidine dimers (CPDs), and several other forms of photoproducts, such as (6-4) pyrimidine dimers, the Dewar valence isomers, hydrates, oxidized bases (mostly 8oxoG) and single strand breaks [34-39]. Absorption of UVC photons produces RNA singlet and triplet excited states. The primary precursor of CPDs and other damages is the singlet state, while the triplet state only plays a limited role (less than 10%) [40, 41]. These damages could block reverse transcription and lead to both the lower overall cDNA yield and lower percentages of gapped reads (**Fig. 4a**).

Damages to nucleic acids in the form of pyrimidine dimers occur much faster than strand breaks. These pyrimidine dimers and hydrates lead to incomplete cDNA products similar to the consequences of failed proximity ligation (even if the proximity ligation worked). Recent studies showed that a short period of UV 254nm irradiation, even within a few seconds, can already strongly block reverse transcription [34, 42]. Given the broad application of UV in molecular biology, a general strategy for reducing photodamage is particularly important.

2. Earlier attempts to reduce UVC damage of RNA.

To reduce the impact of RNA damage on the percentage of gapped reads, we first evaluated the size selection. The shorter cDNA products due to incomplete reverse transcription can be selectively removed. This approach can increase the proportion of gapped reads from successful proximity ligation and processive reverse transcription. However, the size selection protocol can be difficult to establish given that the RNA and cDNA fragments are a broad smear. In addition, the total cDNA yield will be much lower. Other recent studies have attempted to minimize UV damage while maximizing the reversal efficiency by limiting the 254nm UV irradiation time, but the benefit is limited [21]. Given the fast photodamage (which may occur within a few seconds) and that at least 5-10min is needed for reversal of the crosslinks, reducing the reversal time is far from enough for optimal reversal and damage reduction.

3. PhrB (E. coli photolyase) failed to repair UV damaged RNA

In many organisms (except humans), a special enzyme called photolyase can repair CPDs in DNA (**Supplementary Fig. 9a**). In addition, photolyase possess residual activity towards RNA [43]. We tested two commercially available photolyases from E. coli on UVC damaged cDNA or RNA and then performed quantitative PCR to measure the repair efficiency. Modest repair was observed on 254nm UV damaged DNA, but not on RNA (**Supplementary Fig. 9b**, only one PhrB data was showed here). Prolonged incubation with PhrB even reduced the amount of intact RNA. Therefore, we conclude that the residual activity of PhrB is currently insufficient to repair pyrimidine dimers. Nevertheless, engineered PhrB with enhanced activity towards RNA may be useful for repairing RNA.

4. Stronger stacking did not speed up photoreversal of psoralen crosslinking or reduce UVC damage.

Earlier studies showed that T-Pso-T diadducts were much harder to reverse than crosslinked DNA oligo duplexes, suggesting a role of the structural context in efficient reversal [44-47]. Therefore, we tested whether stronger stacking could speed up UVC reversal and reduce the time needed and therefore reduce the damage. Addition of 1M NaCl to the RNA solution, which should help RNA form stable secondary structures, did not speed up the reversal or reduce the damage (**Supplementary Fig. 9c**). It is likely that the stacking of RNA base pairs was not significantly enhanced by the 1M NaCl. In other experiments, we found that the reversal is quite efficient in the DNA and RNA oligo duplexes, suggesting that speeding up the reversal is not a good strategy (**Supplementary Fig. 10i-l**).

5. Denaturing agents failed to prevent UVC damage of RNA.

The excited states of nucleobases that are the energetic precursors to CPDs are extremely short-lived and the formation of CPDs depend on the proper pre-alignment of the neighboring bases in stacked conformations before absorption of the photons. It has been shown that some solvents with hydrophobic groups to compete for base stacking interactions reduce crosslinking efficiency [48-50]. Therefore, we tested whether denaturing conditions can be used to disrupt stacking and reduce UVC induced CPD damage to RNA. Addition of DMSO or formamide to very high percentage did not prevent the damages (**Supplementary Fig. 9d**). This is probably because the properly stacked conformations still exist for a sufficiently long time to allow the dimers to form, despite the ability of these solvents to reduce stacking.

6. Variable efficiency of reverse transcriptases on UVC damaged RNA.

Several types of reverse transcriptases (RTs), including the TGIRT and MarathonRT, have been shown to be highly processive on structured RNA, even ones with chemical adducts [51-53]. UVC causes more dramatic changes in the RNA structure than adducts from structure probing, therefore, we need to identify enzymes that may be more processive on UV damaged RNA. First, we introduced UVC damage by irradiating pure total RNA with 254nm UV for 30min. Then we performed reverse transcription using six RTs, including AffinityScript, MarathonRT, TGIRT-III enzyme (TGIRT), Superscript (SS) II, III and IV (**Supplementary Fig. 9e**). Analysis of 4 RNAs showed that SSIV consistently outperformed other RTs on UVC damaged RNA. However, the 7-fold improvement is far from enough to bypass all damages since 30min UVC irradiation reduces qRT-PCR efficiency ~100-1000 fold for RNA amplicons in the range of 70-180nt (from ACTB and GAPDH mRNAs, **Fig. 4c, Supplementary Fig. 10c-d, g**).

7. Quenchers of nucleobase singlet excited states prevent UVC damages on RNA.

The quantum chemical mechanism of pyrimidine dimer formation in DNA has been studied extensively due to its critical role in UV-induced skin cancers. CPD formation in DNA primarily occurs through the singlet excited states and therefore quenching the singlets can be used to inhibit CPD formation [54-58] (see **Fig. 4b** for a diagram). Several types of singlet quenchers have been shown to inhibit CPDs, including DNA binding dyes proflavine, acridine orange (AO), ethidium bromide (EtBr), methyl green, dimeric Zinc(II)-Cyclen complexes, and the organic solvent acetone [59] (**Supplementary Fig. 10a**). These dyes can bind to double strand nucleic acids by intercalating between adjacent base pairs or by exterior ionic bonding. UVC irradiation in the presence of singlet quenchers can even lead to reversal of DNA pyrimidine dimers [55, 60].

However, later studies did not find proflavine or AO effective in protecting RNA against UVC irradiation, suggesting that differences in DNA and RNA structure may affect the protection efficiency [61]. The lack of protection could be due to the lower concentrations used in these studies (5uM for proflavine and 50uM for AO) and the lower affinity of these dyes toward RNA compared to DNA. Nevertheless, Merriam and Gordon found that the singlet quencher methanol partially protects RNA from UVC damage [56-58]. Methanol used at 90% concentration did not lead to complete protection, suggesting that its protective effect is limited, and more efficient quenchers are needed. Furthermore, many of these dyes absorb light in the UVC range (e.g. **Supplementary Fig. 10b**), raising concerns of the usefulness of such quenchers in the reversal of psoralen crosslinks.

Kleopfer and Morrison showed that acetone has no effect on the dimerization of dimethyl thymidines or DMT, in contrast to the strong inhibition of CPD formation in E. coli DNA reported by Sutherland and Sutherland [49, 57]. This can be potentially explained by the different energetic precursor involved in CPD formation for base monomers vs. polymeric DNA [62]. Greenstock and colleagues showed that the triplet quencher oxygen inhibits dimerization of monomers, such as uracil and thymine, but not dimers like UpU and TpT. In other words, the nucleobase monomers form CPDs primarily through the triplet states that can be quenched by oxygen, while polymers form CPDs primarily through the singlet states that can also be quenched by certain singlet quenchers, such as the DNA dyes and acetone. These studies suggest that singlet quenchers, but not triplet quenchers, are the most likely candidates for protecting RNA against UVC damages.

Based on these previous studies, we set out to systematically test singlet quenchers for protecting RNA from UVC-induced damages (**Supplementary Fig. 10c-h**). All the singlet quenchers protected RNA to various extent. EB and AO showed the highest efficiency at 2.5mM concentrations, protecting ACTB mRNA (180nt PCR amplicon) by ~1000 fold, and GAPDH mRNA (70nt RNA amplicon) by ~50 fold (**Supplementary Fig. 10c-d**). Even RNAs that were treated with PUVA before UVC irradiation was also protected (**Supplementary Fig. 10e-f**). For the PUVA treated samples (ACTB AMT and GAPDH AMT), the absolute quantitative PCR Ct changes (log2(fold) change) after UVC damage with or without prevention were much lower than the non-PUVA treated samples (**Supplementary Fig. 10,** compare panels **c to d** and **d to f**). This is because PUVA lead to additional damages that blunt the UVC damages. The UVC reversal of psoralen crosslinks in the presence of EB and AO lead to net increase in amplifiable RNA compared to crosslinked but not-UVC treated samples (**Supplementary Fig. 10e-f,** EB and AO bars). The level of protection strongly depends on the quencher concentration (**Supplementary Fig. 10g-h)**. The much higher concentrations needed to protect RNA (2.5mM), vs. for DNA (e.g. 50uM AO) can be explained by the lower affinity of these dyes towards RNA [54-58].

We found that AO had a higher prevention efficiency against UVC damages at a lower concentration (0.25 mM) compared to EB (2.5 mM) (**Supplementary Fig. 10g-h**). In order to better remove these quenchers after crosslink reversal, we chose lower concentration of AO (0.25 mM) to prevent UVC damages (**Fig. 4c**). Using the 30min UVC irradiation to reverse crosslink, the protection of AO for ACTB mRNA was 62.6-fold (from 0.005 to 0.313) without PUVA, and 6.46 with PUVA. The protection of GAPDH mRNA was 29.18-fold (from 0.016 to 0.467) without PUVA (**Fig. 4c**). For PUVA treated samples, there is also PUVA-induced oxidative damages (see **Supplementary Note 5** for details) and simultaneous reversal of crosslinking, so it is much more difficult to calculate the contribution of the singlet quencher effect.

Higher concentrations of acridine orange are needed to protect RNA given its lower binding affinity compared to DNA. To test whether higher affinity nucleic acid intercalators are more effective, we used a dimer of EB, EthD1 (**Supplementary Fig. 10a**), which has a binding affinity of $2x10^8 M^{-1}$, compared to $1.5x10^5 M^{-1}$ for EB [63, 64]. However, we found that the high binding affinity intercalators were difficult to remove from the RNA samples after reversal, and the protection was not higher than EB. The SYBR dyes, SYBR Green I and SYBR Gold did not protect RNA well compared to EB and AO (**Supplementary Fig. 11**). The SYBR dyes and acetone partially protected DNA against UVC damages, while AO almost completely protected DNA. Even for DNA that was first crosslinked with psoralen, there was little reduction in PCR efficiency after the 254nm reversal, suggesting little damage after both PUVA and UVC treatment (**Supplementary Fig. 11**). Together these results showed both similarities and differences in the susceptibility and prevention of UVC damages on DNA and RNA.

8. Calculating damages and prevention using the Poisson distribution.

For these calculations, we focus on the normal RNA (without psoralen crosslinking), because the crosslinking induces other types of damages. We assume that the dimers follow a Poisson distribution [61], $P(k) = \lambda^k e^{-\lambda}/k!$, where the interval is defined as the RNA sequence to be reverse transcribed, e.g. 70nt for the GAPDH amplicon; λ is the average number of dimers in the RNA sequence interval; k is the number of dimers in the RNA sequence interval, and takes the values of 0, 1, 2 …. k has an upper limit determined by the sequence length and base composition.

For example, there are 18 potential pyrimidine dimer sites in the GAPDH mRNA amplicon, 11 of which are in the actual reverse transcribed region, with 7 at the primer binding site. The effect of the dimers on reverse transcription depends on their locations, for example, different places within the primer binding sites or in the actual reverse transcribed region to be amplified. Each dimer in the actual reverse transcribed region only partially blocks reverse transcription, therefore, the actual effect on RT need to be determined by a separate factor. We do not know the exact bypass percentage but it is very low based on previous studies [65]. For simplicity, we define a complete damage site as one that completely blocks the RT. The following calculations are based on **Supplementary Fig. 10c-d** for ACTB and GAPDH mRNAs.

| mRNA | UVC | AO | delta_Ct | % intact RNA | Complete damage sites | Protection % vs. w/o AO |
|---|---|---|---|---|---|---|
| ACTB | 10min | – | -5.64 | 2.0 | 3.91 | |
| ACTB | 10min | + | -0.92 | 52.9 | 0.64 | 84 |
| ACTB | 30min | – | -11.07 | 0.05 | 7.67 | |
| ACTB | 30min | + | -1.27 | 41.5 | 0.88 | 89 |
| GAPDH | 10min | – | -3.39 | 9.5 | 2.35 | |
| GAPDH | 10min | + | -0.81 | 57.0 | 0.56 | 76 |
| GAPDH | 30min | – | -6.62 | 1.0 | 4.59 | |
| GAPDH | 30min | + | -0.99 | 50.3 | 0.69 | 85 |

In the presence of AO, ~50% RNA remained intact after 10-30min irradiation. The addition of AO protected both mRNAs by 76%-89%. UVC-induced CPD dimerization is a reversible reaction [54], therefore, the protection will not be 100%, unless all pyrimidines are completely sequestered by quenchers or other types of blockers (see quantitative analysis as follows). In addition, other less frequent types UVC-induced damages, such as strand breaks and oxidative damages, are not reversible, further limiting the extent of protection.

To understand the UVC damage and prevention, using AO as an example, here we treat the process as an equilibrium.

$$M_{quenched} \overset{AO}{\leftrightharpoons} M \underset{k_2}{\overset{k_1}{\rightleftharpoons}} D, \qquad r_1 = k_1[M], \qquad r_2 = k_2[D]$$

Here M stands for monomer, D stands for dimer. Both $k_1$ and $k_2$ are unimolecular reaction constants that depend on UV wavelength. When AO is present, it affects the effective [M] and UV dose at the same time. Increasing AO reduces [M] and effective UV dose, which will require longer time to reach equilibrium. In reality there will always be forward reaction as long as [M] is positive. At equilibrium $\mathrm{r_1 = k_1[M] = r_2 = k_2[D]}$, therefore, $\mathrm{[M]/[D] = k_2/k_1}$, the ratio of [M]/[D] will not change at a fixed wavelength. This analysis shows that it is impossible to completely prevent UVC induced pyrimidine dimers.

9. Singlet state quenchers do not block photoreversal of psoralen crosslinks

To study whether AO will block the reversal of crosslinking, we designed 8-mer DNA and RNA oligos to perform crosslinking and reverse crosslinking test (**Supplementary Fig. 10i-l**). DNA oligos were crosslinked much more efficiently than RNA oligos (compare panel i vs. k). Most RNA oligos only formed monoadducts. Then, these crosslinked products were reversed by 254nm UV, with or without AO. The reversal was completed within 10min, and the presence of AO did not block reversal (**Supplementary Fig. 10k, l**). These results suggest that the absorption of UV light by singlet quenchers does not affect reversal significantly (**Supplementary Fig. 10b**).

10. Summary and discussion

In this systematic optimization, we have discovered that extensive dimers caused the low efficiency of reverse transcription, reducing the yield of cDNA and percentage of gapped reads. We found that SSIV reverse transcriptase was more efficient on UVC damaged RNA. Several intercalating dyes and solvents previously found to protect DNA from UVC damage by quenching the singlet states of excited DNA can also protect RNA from UVC damage, although much higher concentrations are needed. Importantly, these dyes do not block the reversal of psoralen crosslinks.

In summary we developed the first method for efficiently protecting RNA against UVC damage. This important discovery will prove useful for many problems in RNA biology given the broad application of UV irradiation and the growing interests in RNA structure analysis using photochemical crosslinking. For example, this method can be useful for other RNA experiments that involve UVC irradiation, such as analysis of RNA structures by chemical probing [34, 42], and RNA-protein crosslinking studies [66].

**Supplementary Note 5. Bypass of PUVA induced oxidative damage on RNA**

The combination of photosensitizers and UVA irradiation causes extensive oxidative damage to nucleic acids, including base oxidations and subsequent strand breaks [38, 39, 67, 68]. RNA is likely subject to more oxidative damages than DNA due to its location in cells and the structural differences [69, 70]. Such damages will block reverse transcription and reduce cDNA yield in library preparation [71]. In early studies, we noticed that crosslinking induced extensive DNA fragmentation, but few strand breaks on RNA. This is probably due to the DNA repair on psoralen adducts that induce double strand breaks. The photosensitized oxidations also damage other cellular components, including proteins and lipids. The oxidative damages, in addition nucleic acid crosslinking, have been suggested as a major factor in the therapeutic, as well as side effects of PUVA therapies. In PARIS, the protection of RNA against UVC damage was insufficient to keep RNA intact. Therefore, we set out to determine the extent of PUVA induced oxidative damages and develop approaches to prevent, repair or by-pass these damages.

<u>1. PUVA induces RNA damages</u>

First, we used cDNA yield of RT-qPCR to test PUVA damages on RNA molecules. We found that PUVA, but not UVA alone, blocked reverse transcription (**Supplementary Fig. 13a-b**). Lower concentration of AMT (0.05 mg/ml) induced less damage, but also reduced crosslinking efficiency (**Supplementary Figs. 13c-d**). In order to reduce the PUVA damages, we also test the effects of separating the incubation and crosslinking steps to reduce effective psoralen concentrations during crosslinking. HEK293T cells were incubation with 0.5 mg/ml AMT solution for 15 mins, to make sure AMT penetrate into RNA duplex (**Supplementary Figs. 13e-f**). Then AMT solution was changed to PBS solution to perform crosslinking by 365 nm UV for 30 mins. This design slightly reduced RNA damage, but also greatly decreased crosslinking efficiency due to dissociation of AMT from nucleic acids after washing out AMT.

<u>2. Antioxidants reduce PUVA damage but also block psoralen crosslinking.</u>

Next, we explored the possibility of using antioxidants and reactive oxygen species (ROS) quenchers to reduce PUVA-induced damages (**Supplementary Fig. 14a-b**). Psoralen (AMT) can serve as a photosensitizer for UVA, which involves a direct energy transfer reaction between triplet states of excited AMT and ground state oxygen, producing highly reactive ROS that can oxidize RNA base and generate PUVA damages on RNA (**Supplementary Fig. 14b**). In addition, guanine has the lowest one-electron oxidation potential of the nucleobases. The excited singlet state of intercalated AMT can directly react with DNA/RNA molecules, especially guanine, resulting radical guanine and further predominantly from oxidation of guanine. All these PUVA damages will also block the reverse transcription efficiency.

Earlier studies have shown that certain antioxidants and ROS quenchers can reduce PUVA-induced oxidative damages using both in vitro and in vivo models. These chemicals act at different stages of the oxidation process. For example, $O_2$•-scavenger: Tiron and MnTBAP [72]; •OH scavenger: Mannitol [73, 74], DMSO and Glycerol; $^1O_2$ scavenger: $NaN_3$ [67, 75], general radical scavenger: Vitamin C (VC) [76, 77] (**Supplementary Fig. 14a**). The effects of ROS scavengers on PUVA damaged RNA were tested by RT-qPCR and found to be highly variable (**Supplementary Fig. 14c-g**). For example, mannitol and $NaN_3$ had no effect on PUVA damages, Tiron and MnTBAP partially blocked damages, while VC almost completely blocked damages at 100mM concentration. However, the reduction of damages was accompanied by loss of crosslinking (**Supplementary Fig. 14h-j**), suggesting that these compounds quenched a common mechanism in crosslinking and photooxidative damage.

Recent studies showed that psoralen crosslinking involves an electron transfer from DNA to AMT, similar to the electron-transfer induced guanine oxidation [78, 79]. In particular, intercalated psoralen (AMT) is excited to singlet state, and directly induces electron transfer from DNA, charge recombination and crosslinking for pyrimidines, or oxidation for guanines. Together, studies presented here and in earlier publications suggest that it is impossible to block oxidative damage without blocking crosslinking. Furthermore, our studies further suggest that, during PUVA therapy, the protective effects of antioxidants and ROS scavengers are at least partially based on their abilities to block crosslinking, in addition to blocking the oxidation of cellular components.

<u>3. DNA polymerases cannot bypass PUVA-induced RNA damages.</u>

We noticed that PUVA-treated DNA can be amplified in PCR without obvious reduction of Ct value in PCR, suggesting that DNA polymerases can bypass the oxidative damages on DNA samples. Several earlier studies showed that certain DNA polymerases also possess reverse transcriptase activity [80-84]. A few of them, e.g. Bst, Klenow LF, Klenow exo-, were reported to have comparable levels of RT activity to AMV RT for templates shorter than 125nt [85]. Therefore, we tested the possibility of using DNA polymerases in bypassing PUVA-induced oxidative damages on RNA. However, despite extensive tests of conditions, we were unable to obtain comparable levels of cDNA yield using the DNA polymerases (**Supplementary Fig. 15a**).

<u>4. Optimized RT conditions improve bypass of PUVA-induced oxidative damages</u>

Earlier studies showed reverse transcriptases can bypass oxidative damages pn RNA but the efficiency is very low [86]. Multiple types of oxidized guanines can be bypassed to various degrees. During reverse transcription, the enzyme rackets between active and inactive states [87], and the RNA damages trap the RT enzymes in the inactive state for much longer time. Van Nostrand et al. found that reverse transcriptase read-through RNA with peptide adducts was highly dependent on the identity of the reverse transcriptase enzyme as well as on buffer conditions [88]. Therefore, we systematically screened for reverse transcription conditions to increase bypass efficiency.

Higher ratios of enzyme:substrate increase processivity, especially for MMLV derived reverse transcriptases, such as superscript II, III and IV [89]. Because RNA-DNA hybrids can sequester reverse transcriptases and reduce available enzymes for cDNA synthesis, longer incubation time also helps improve the cDNA products yield [89]. We first tested higher amounts of RT enzyme SSIV in reverse transcription. HEK293T cells were crosslinked by 365 nm UV for 30 mins with 0.5 mg/ml AMT, then reverse crosslinking was performed with the protection of acridine orange. Reverse transcription with higher amounts of SSIV did not result in higher cDNA yield (**Supplementary Fig. 15b**).

Next we tested various RT enzymes on PUVA damaged RNA, using primer extension assays and reverse transcription PCR. For the primer extension assay, we crosslinked a 48-mer RNA template oligo to mimic the PUVA damages (**Fig. 4d**). After photo-reversal by 254 nm UV with AO protection, reverse crosslinked RNA oligo was used to primer extension assay. For the PCR assay, we used primers targeting several mRNAs and noncoding RNAs. SSIV outperformed other RT enzymes, including MarathonRT and TGIRT, both of which were previously reported to be highly processive on structured and modified RNA molecules [51-53] (**Supplementary Fig. 15c,** performed in $Mn^{2+}$ buffer, and **Fig. 4f,** performed in $Mg^{2+}$ buffer, which is then quantified in **Supplementary Fig. 15d**).

Several studies suggested that $Mn^{2+}$ induce more bypass of RT blocks in many different conditions, including chemical adducts in SHAPE experiments, and peptide adducts in CLIP experiments [88, 90]. We tested RT bypass of PUVA damages under different concentrations of $Mn^{2+}$. Primer extension assays and RT-qPCR data suggested that SSIV with 1.5 mM $Mn^{2+}$ buffer is the most effective on PUVA damaged RNA (**Fig. 4g, i,** and **Supplementary Fig. 15g-i**).

The racketing behavior of RT enzymes between active and inactive states suggests that longer incubation may increase the bypass of damaged RNA. Therefore, we tested the effect of different RT incubation time using primer extension and RT-qPCR assays (**Fig. 4g, i,** and **Supplementary Fig. 15g-i**). The longer incubation time dramatically improved cDNA yield for several different RNA targets. Together these optimizations identified best combination of conditions that improve RT efficiency on PUVA damaged RNA.

5. Summary and discussion.

In summary, our extensive studies of PUVA-induced oxidative damages clarified important mechanisms in the process. We found that certain antioxidants and ROS scavengers can reduce oxidative damage, but also block crosslinking at the same time. Our in-depth analysis of the reverse transcriptase conditions suggests new variations that greatly improve reverse transcriptase processivity and cDNA yield. We found that the reverse transcriptase activity on damaged RNA can be enhanced by several treatments, including the SSIV variant of enzyme, cofactor $Mn^{2+}$, and much longer incubation. The prolonged incubation is especially effective in promoting the bypass.

**References**

1. Chomczynski, P. & Sacchi, N. Single-step method of RNA isolation by acid guanidinium thiocyanate-phenol-chloroform extraction. *Anal Biochem* **162**, 156-159 (1987).
2. Chomczynski, P. (Google Patents, 1989).
3. Kim, Y.K., Yeo, J., Kim, B., Ha, M. & Kim, V.N. Short structured RNAs with low GC content are selectively lost during extraction from a small number of cells. *Molecular cell* **46**, 893-895 (2012).
4. Lu, Z., Gong, J. & Zhang, Q.C. PARIS: Psoralen Analysis of RNA Interactions and Structures with High Throughput and Resolution. *Methods in molecular biology* **1649**, 59-84 (2018).
5. Lu, Z. et al. RNA Duplex Map in Living Cells Reveals Higher-Order Transcriptome Structure. *Cell* **165**, 1267-1279 (2016).
6. Sastry, S.S., Ross, B.M. & P'Arraga, A. Cross-linking of DNA-binding proteins to DNA with psoralen and psoralen furan-side monoadducts. Comparison of action spectra with DNA-DNA cross-linking. *The Journal of biological chemistry* **272**, 3715-3723 (1997).
7. Sastry, S.S. et al. Laser-induced protein-DNA cross-links via psoralen furanside monoadducts. *Biochemistry* **32**, 5526-5538 (1993).
8. Zechel, K. & Weber, K. Degradation of nucleic acids in cell lysates by S1 nuclease in the presence of 9 M urea and sodium dodecylsulfate. *European journal of biochemistry / FEBS* **77**, 133-139 (1977).
9. Lapanje, S. Denaturation of globular proteins by guanidine thiocyanate. I. Optical rotation in aqueous guanidine thiocyanate solutions. *Biochimica et biophysica acta* **243**, 349-356 (1971).
10. Nwokeoji, A.O., Kilby, P.M., Portwood, D.E. & Dickman, M.J. Accurate Quantification of Nucleic Acids Using Hypochromicity Measurements in Conjunction with UV Spectrophotometry. *Anal Chem* **89**, 13567-13574 (2017).
11. Yoon, J.H. & Lee, C.S. Sequence specificity for DNA interstrand cross-linking induced by anticancer drug chlorambucil. *Arch Pharm Res* **20**, 550-554 (1997).
12. Tan, S.C. & Yiap, B.C. DNA, RNA, and protein extraction: the past and the present. *J Biomed Biotechnol* **2009**, 574398 (2009).
13. Ziv, O. et al. COMRADES determines in vivo RNA structures and interactions. *Nat Methods* **15**, 785-788 (2018).
14. Urdaneta, E.C. et al. Purification of cross-linked RNA-protein complexes by phenol-toluol extraction. *Nat Commun* **10**, 990 (2019).
15. Queiroz, R.M.L. et al. Comprehensive identification of RNA-protein interactions in any organism using orthogonal organic phase separation (OOPS). *Nat Biotechnol* **37**, 169-178 (2019).
16. Trendel, J. et al. The Human RNA-Binding Proteome and Its Dynamics during Translational Arrest. *Cell* **176**, 391-403.e319 (2019).

17. Shchepachev, V. et al. Defining the RNA interactome by total RNA-associated protein purification. *Mol Syst Biol* **15**, e8689 (2019).
18. Lu, Z. & Chang, H.Y. The RNA Base-Pairing Problem and Base-Pairing Solutions. *Cold Spring Harb Perspect Biol* **10** (2018).
19. Lu, Z., Carter, A.C. & Chang, H.Y. Mechanistic insights in X-chromosome inactivation. *Philos Trans R Soc Lond B Biol Sci* **372** (2017).
20. Sharma, E., Sterne-Weiler, T., O'Hanlon, D. & Blencowe, B.J. Global Mapping of Human RNA-RNA Interactions. *Mol Cell* **62**, 618-626 (2016).
21. Aw, J.G. et al. In Vivo Mapping of Eukaryotic RNA Interactomes Reveals Principles of Higher-Order Organization and Regulation. *Mol Cell* **62**, 603-617 (2016).
22. Wassarman, D.A. Psoralen crosslinking of small RNAs in vitro. *Mol Biol Rep* **17**, 143-151 (1993).
23. Xiao, J., Feehery, C.E., Tzertzinis, G. & Maina, C.V. E. coli RNase III(E38A) generates discrete-sized products from long dsRNA. *Rna* **15**, 984-991 (2009).
24. Dunn, J.J. RNase III cleavage of single-stranded RNA. Effect of ionic strength on the fideltiy of cleavage. *The Journal of biological chemistry* **251**, 3807-3814 (1976).
25. Gasparro, F.P., Saffran, W.A., Cantor, C.R. & Edelson, R.L. Wavelength dependence for AMT crosslinking of pBR322 DNA. *Photochem Photobiol* **40**, 215-219 (1984).
26. Spitale, R.C. et al. Structural imprints in vivo decode RNA regulatory mechanisms. *Nature* **519**, 486-490 (2015).
27. Sugimoto, Y. et al. hiCLIP reveals the in vivo atlas of mRNA secondary structures recognized by Staufen 1. *Nature* **519**, 491-494 (2015).
28. Vigne, R. & Jordan, B.R. Conformational analysis of RNA molecules by partial RNAse digestion and two dimensional acrylamide gel electrophoresis. Application to E. coli 5S RNA. *Biochimie* **53**, 981-986 (1971).
29. Zwieb, C. & Brimacombe, R. Localisation of a series of intra-RNA cross-links in 16S RNA, induced by ultraviolet irradiation of Escherichia coli 30S ribosomal subunits. *Nucleic acids research* **8**, 2397-2411 (1980).
30. Thompson, J.F. & Hearst, J.E. Structure of E. coli 16S RNA elucidated by psoralen crosslinking. *Cell* **32**, 1355-1365 (1983).
31. Lu, Z. et al. Metazoan tRNA introns generate stable circular RNAs in vivo. *Rna* **21**, 1554-1565 (2015).
32. Ruskin, B., Krainer, A.R., Maniatis, T. & Green, M.R. Excision of an intact intron as a novel lariat structure during pre-mRNA splicing in vitro. *Cell* **38**, 317-331 (1984).
33. Wang, X., Li, C., Gao, X., Wang, J. & Liang, X. Preparation of Small RNAs Using Rolling Circle Transcription and Site-Specific RNA Disconnection. *Mol Ther Nucleic Acids* **4**, e215 (2015).
34. Kladwang, W., Hum, J. & Das, R. Ultraviolet shadowing of RNA can cause significant chemical damage in seconds. *Scientific reports* **2**, 517 (2012).
35. Cadet, J. & Wagner, J.R. DNA base damage by reactive oxygen species, oxidizing agents, and UV radiation. *Cold Spring Harbor perspectives in biology* **5** (2013).
36. Cadet, J. & Douki, T. Formation of UV-induced DNA damage contributing to skin cancer development. *Photochem Photobiol Sci* **17**, 1816-1841 (2018).
37. Remsen, J.F., Miller, N. & Cerutti, P.A. Photohydration of uridine in the RNA of coliphage R17. II. The relationship between ultraviolet inactivation and uridine photohydration. *Proc Natl Acad Sci U S A* **65**, 460-466 (1970).
38. Pathak, M.A. Mechanisms of psoralen photosensitization reactions. *Natl Cancer Inst Monogr* **66**, 41-46 (1984).
39. Pathak, M.A. & Fitzpatrick, T.B. The evolution of photochemotherapy with psoralens and UVA (PUVA): 2000 BC to 1992 AD. *J Photochem Photobiol B* **14**, 3-22 (1992).
40. Banyasz, A. et al. Electronic excited states responsible for dimer formation upon UV absorption directly by thymine strands: joint experimental and theoretical study. *J Am Chem Soc* **134**, 14834-14845 (2012).
41. Liu, L., Pilles, B.M., Gontcharov, J., Bucher, D.B. & Zinth, W. Quantum Yield of Cyclobutane Pyrimidine Dimer Formation Via the Triplet Channel Determined by Photosensitization. *J Phys Chem B* **120**, 292-298 (2016).
42. Greenfeld, M., Solomatin, S.V. & Herschlag, D. Removal of covalent heterogeneity reveals simple folding behavior for P4-P6 RNA. *The Journal of biological chemistry* **286**, 19872-19879 (2011).
43. Selby, C.P. & Sancar, A. A cryptochrome/photolyase class of enzymes with single-stranded DNA-specific photolyase activity. *Proceedings of the National Academy of Sciences of the United States of America* **103**, 17696-17700 (2006).
44. Cimino, G.D., Shi, Y.B. & Hearst, J.E. Wavelength dependence for the photoreversal of a psoralen-DNA cross-link. *Biochemistry* **25**, 3013-3020 (1986).
45. Shi, Y.B. & Hearst, J.E. Wavelength dependence for the photoreactions of DNA-psoralen monoadducts. 1. Photoreversal of monoadducts. *Biochemistry* **26**, 3786-3792 (1987).
46. Shi, Y.B. & Hearst, J.E. Wavelength dependence for the photoreactions of DNA-psoralen monoadducts. 2. Photo-cross-linking of monoadducts. *Biochemistry* **26**, 3792-3798 (1987).
47. Thompson, J.F., Bachellerie, J.P., Hall, K. & Hearst, J.E. Dependence of 4'-(hydroxymethyl)-4,5',8-trimethylpsoralen photoaddition on the conformation of ribonucleic acid. *Biochemistry* **21**, 1363-1368 (1982).
48. Olmon, E. Solvent effects on photodegradation of the 18-mer of thymidylic acid. *OSU Knowledge Base* (2005).
49. Kleopfer, R. & Morrison, H. Organic photochemistry. XVII. Solution-phase photodimerization of dimethylthymine. *Journal of the American Chemical Society* **94**, 255-264 (1972).
50. Piskur, J. & Rupprecht, A. Aggregated DNA in ethanol solution. *FEBS Lett* **375**, 174-178 (1995).
51. Zubradt, M. et al. DMS-MaPseq for genome-wide or targeted RNA structure probing in vivo. *Nature methods* **14**, 75-82 (2017).
52. Zhao, C., Liu, F. & Pyle, A.M. An ultraprocessive, accurate reverse transcriptase encoded by a metazoan group II intron. *Rna* **24**, 183-195 (2018).
53. Mohr, S. et al. Thermostable group II intron reverse transcriptase fusion proteins and their use in cDNA synthesis and next-generation RNA sequencing. *Rna* **19**, 958-970 (2013).

54. Beukers, R. The effect of proflavine on U.V.-induced dimerization of thymine in DNA. *Photochem Photobiol* **4**, 935-937 (1965).
55. Setlow, R.B. & Carrier, W.L. Formation and destruction of pyrimidine dimers in polynucleotides by ultra-violet irradiation in the presence of proflavine. *Nature* **213**, 906-907 (1967).
56. Sutherland, B.M. & Sutherland, J.C. Inhibition of pyrimidine dimer formation in DNA by cationic molecules: role of energy transfer. *Biophys J* **9**, 1045-1055 (1969).
57. Sutherland, B.M. & Sutherland, J.C. Mechanisms of inhibition of pyrimidine dimer formation in deoxyribonucleic acid by acridine dyes. *Biophys J* **9**, 292-302 (1969).
58. Sutherland, J.C. & Sutherland, B.M. Ethidium bromide-DNA complex: wavelength dependence of pyrimidine dimer inhibition and sensitized fluorescence as probes of excited states. *Biopolymers* **9**, 639-653 (1970).
59. Aoki, S., Sugimura, C. & Kimura, E. Efficient Inhibition of Photo[2 + 2]cycloaddition of Thymidilyl(3‘−5‘)thymidine and Promotion of Photosplitting of the cis-syn-Cyclobutane Thymine Dimer by Dimeric Zinc(II)−Cyclen Complexes Containing m- and p-Xylyl Spacers. *Journal of the American Chemical Society* **120**, 10094-10102 (1998).
60. Setlow, J.K. & Setlow, R.B. Contribution of dimers containing cytosine to ultra-violet inactivation of transforming DNA. *Nature* **213**, 907-909 (1967).
61. Merriam, V. & Gordon, M.P. Pyrimidine dimer formation in ultraviolet irradiated TMV-RNA. *Photochem Photobiol* **6**, 309-319 (1967).
62. Greenstock, C.L., Brown, I.H., Hunt, J.W. & Johns, H.E. Photodimerization of pyrimidine nucleic acid derivatives in aqueous solution and the effect of oxygen. *Biochem Biophys Res Commun* **27**, 431-436 (1967).
63. Glazer, A.N. & Rye, H.S. Stable dye-DNA intercalation complexes as reagents for high-sensitivity fluorescence detection. *Nature* **359**, 859-861 (1992).
64. Gaugain, B., Barbet, J., Capelle, N., Roques, B.P. & Le Pecq, J.B. DNA Bifunctional intercalators. 2. Fluorescence properties and DNA binding interaction of an ethidium homodimer and an acridine ethidium heterodimer. *Biochemistry* **17**, 5078-5088 (1978).
65. Smith, C.A., Baeten, J. & Taylor, J.S. The ability of a variety of polymerases to synthesize past site-specific cis-syn, trans-syn-II, (6-4), and Dewar photoproducts of thymidylyl-(3'-->5')-thymidine. *J Biol Chem* **273**, 21933-21940 (1998).
66. Wheeler, E.C., Van Nostrand, E.L. & Yeo, G.W. Advances and challenges in the detection of transcriptome-wide protein-RNA interactions. *Wiley Interdiscip Rev RNA* **9** (2018).
67. Zhang, X., Rosenstein, B.S., Wang, Y., Lebwohl, M. & Wei, H. Identification of possible reactive oxygen species involved in ultraviolet radiation-induced oxidative DNA damage. *Free Radic Biol Med* **23**, 980-985 (1997).
68. Zhang, X. et al. Induction of 8-oxo-7,8-dihydro-2'-deoxyguanosine by ultraviolet radiation in calf thymus DNA and HeLa cells. *Photochem Photobiol* **65**, 119-124 (1997).
69. Nunomura, A. et al. Oxidative damage to RNA in neurodegenerative diseases. *J Biomed Biotechnol* **2006**, 82323 (2006).
70. Wamer, W.G. & Wei, R.R. In vitro photooxidation of nucleic acids by ultraviolet A radiation. *Photochem Photobiol* **65**, 560-563 (1997).
71. Rhee, Y., Valentine, M.R. & Termini, J. Oxidative base damage in RNA detected by reverse transcriptase. *Nucleic Acids Res* **23**, 3275-3282 (1995).
72. Bleeke, T., Zhang, H., Madamanchi, N., Patterson, C. & Faber, J.E. Catecholamine-induced vascular wall growth is dependent on generation of reactive oxygen species. *Circ Res* **94**, 37-45 (2004).
73. Goldstein, S. & Czapski, G. Mannitol as an OH. scavenger in aqueous solutions and in biological systems. *Int J Radiat Biol Relat Stud Phys Chem Med* **46**, 725-729 (1984).
74. Pelle, E. et al. Ultraviolet-B-induced oxidative DNA base damage in primary normal human epidermal keratinocytes and inhibition by a hydroxyl radical scavenger. *J Invest Dermatol* **121**, 177-183 (2003).
75. Silva-Júnior, A.C., Asad, L.M., Felzenszwalb, I. & Asad, N.R. Mutagenicity induced by UVC in Escherichia coli cells: reactive oxygen species involvement. *Redox Rep* **16**, 187-192 (2011).
76. Besaratinia, A., Kim, S.I., Bates, S.E. & Pfeifer, G.P. Riboflavin activated by ultraviolet A1 irradiation induces oxidative DNA damage-mediated mutations inhibited by vitamin C. *Proc Natl Acad Sci U S A* **104**, 5953-5958 (2007).
77. Darr, D., Combs, S., Dunston, S., Manning, T. & Pinnell, S. Topical vitamin C protects porcine skin from ultraviolet radiation-induced damage. *Br J Dermatol* **127**, 247-253 (1992).
78. Fröbel, S., Reiffers, A., Torres Ziegenbein, C. & Gilch, P. DNA Intercalated Psoralen Undergoes Efficient Photoinduced Electron Transfer. *J Phys Chem Lett* **6**, 1260-1264 (2015).
79. Fröbel, S., Levi, L., Ulamec, S.M. & Gilch, P. Photoinduced Electron Transfer between Psoralens and DNA: Influence of DNA Sequence and Substitution. *Chemphyschem* **17**, 1377-1386 (2016).
80. Jones, M.D. & Foulkes, N.S. Reverse transcription of mRNA by Thermus aquaticus DNA polymerase. *Nucleic Acids Res* **17**, 8387-8388 (1989).
81. Grabko, V.I., Chistyakova, L.G., Lyapustin, V.N., Korobko, V.G. & Miroshnikov, A.I. Reverse transcription, amplification and sequencing of poliovirus RNA by Taq DNA polymerase. *FEBS Lett* **387**, 189-192 (1996).
82. Myers, T.W. & Gelfand, D.H. Reverse transcription and DNA amplification by a Thermus thermophilus DNA polymerase. *Biochemistry* **30**, 7661-7666 (1991).
83. Bustin, S.A. Absolute quantification of mRNA using real-time reverse transcription polymerase chain reaction assays. *J Mol Endocrinol* **25**, 169-193 (2000).
84. Sellner, L.N. & Turbett, G.R. Comparison of three RT-PCR methods. *Biotechniques* **25**, 230-234 (1998).
85. Shi, C., Shen, X., Niu, S. & Ma, C. Innate Reverse Transcriptase Activity of DNA Polymerase for Isothermal RNA Direct Detection. *J Am Chem Soc* **137**, 13804-13806 (2015).

86. Alenko, A., Fleming, A.M. & Burrows, C.J. Reverse Transcription Past Products of Guanine Oxidation in RNA Leads to Insertion of A and C opposite 8-Oxo-7,8-dihydroguanine and A and G opposite 5-Guanidinohydantoin and Spiroiminodihydantoin Diastereomers. *Biochemistry* **56**, 5053-5064 (2017).
87. Furge, L.L. & Guengerich, F.P. Analysis of nucleotide insertion and extension at 8-oxo-7,8-dihydroguanine by replicative T7 polymerase exo- and human immunodeficiency virus-1 reverse transcriptase using steady-state and pre-steady-state kinetics. *Biochemistry* **36**, 6475-6487 (1997).
88. Van Nostrand, E.L., Shishkin, A.A., Pratt, G.A., Nguyen, T.B. & Yeo, G.W. Variation in single-nucleotide sensitivity of eCLIP derived from reverse transcription conditions. *Methods* **126**, 29-37 (2017).
89. Gerard, G.F. et al. The role of template-primer in protection of reverse transcriptase from thermal inactivation. *Nucleic Acids Res* **30**, 3118-3129 (2002).
90. Siegfried, N.A., Busan, S., Rice, G.M., Nelson, J.A. & Weeks, K.M. RNA motif discovery by SHAPE and mutational profiling (SHAPE-MaP). *Nat Methods* **11**, 959-965 (2014).

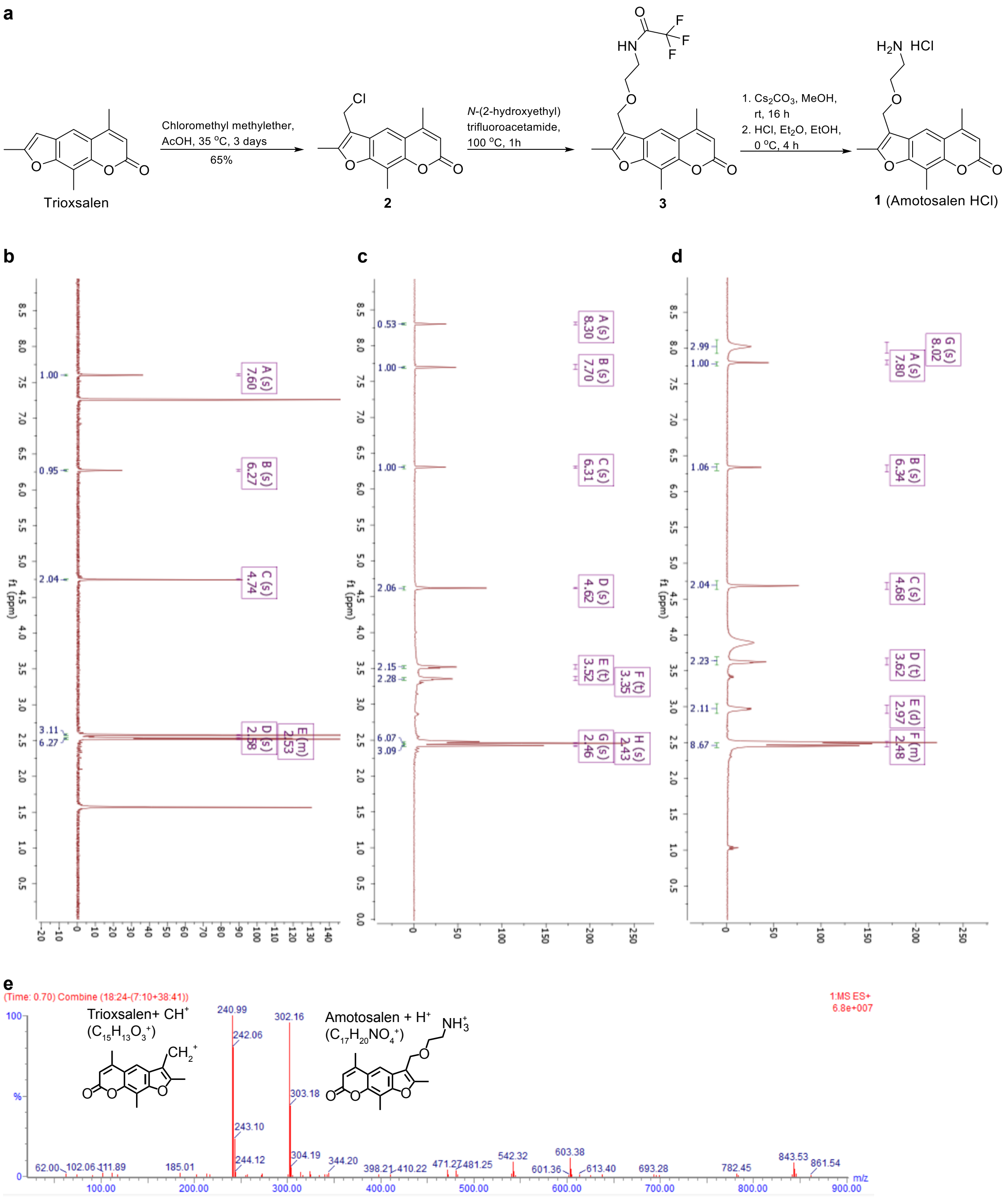

**Supplementary Figure 1. Synthesis and characterization of amotosalen HCl salt.** **a**, Synthesis route and conditions. **b-d,** $^{1}H$ NMR of the three compounds **2**, **3**, and **1**. **e**, Mass spectrometry analysis of compound **1**, Amotosalen HCl. Intact molecular ion and a fragment are identified. See Supplementary Methods for details on synthesis and characterization.

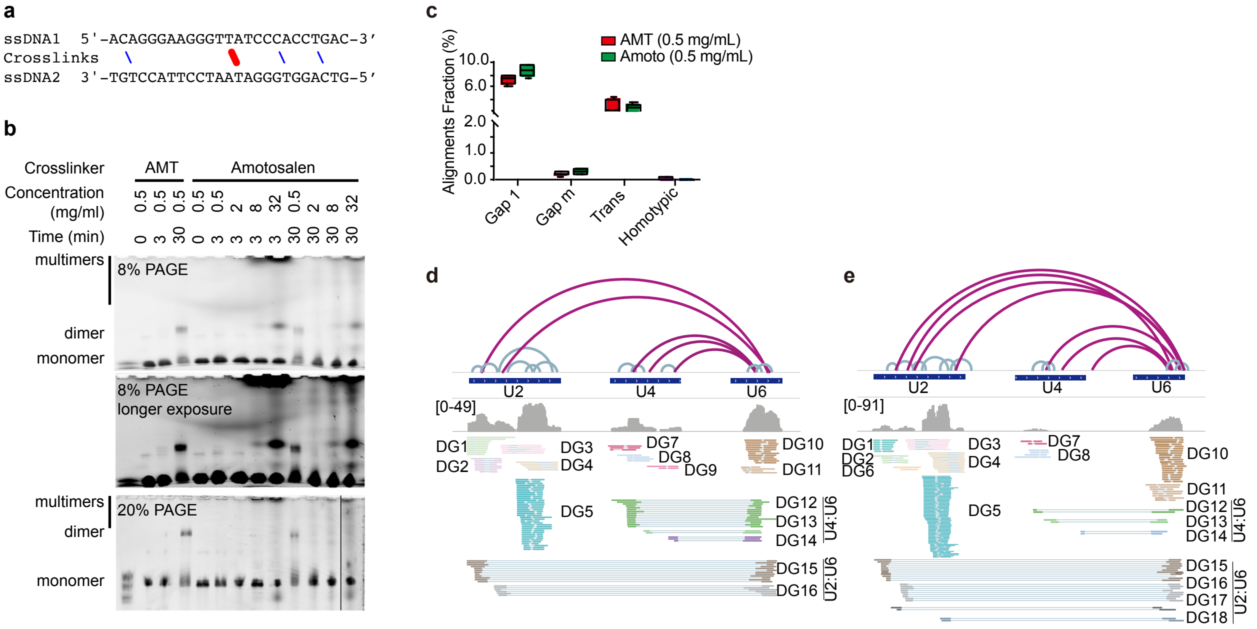

**Supplementary Figure 2.** ***In vitro*** **and** ***in vivo*** **tests of amotosalen crosslinking.** **a**, Secondary structure of the DNA oligo duplex for testing psoralen crosslinking. Blue lines: staggered T-C or C-T pairs as potential psoralen crosslinking sites. Red line: staggered T-T pair. The TpA sequence is the primary crosslinked site. Note: there is one G-A and one G-T mismatch. **b**, DNA duplex crosslinked with AMT and amotosalen at various concentrations and different times. Experimental conditions are as follows. The two oligos were mixed, heated up to 90ºC and slowly cooled down to anneal. In a 20ul system, added 25pmole of each DNA oligo, 2ul 10x PBS, and the cross-linker. Addition of PBS caused some amotosalen to precipitate out of solution in the 32mg/ml group of experiments. After crosslinking, water was added to compensate for evaporation. Samples were run in 8% or 20% urea-TBE PAGE and subjected to different exposure times. siRNA ladder in the 20% PAGE: 17, 21 and 25 nt. Higher amotosalen concentrations resulted in DNA oligos stuck in the well, likely due to their large size and complex structure. **c**, Fraction comparison of non-continuous alignments from 0.5mg/ml AMT and 0.5mg/ml amotosalen crosslinked samples. After sequencing, non-continuous alignments containing RNA structure and interaction information were divided into four different types: gap1 alignment (one gap, 2 segments); gapm alignment (multiple gaps, multiple segments); trans alignment (segments mapped to different chromosomes or strands); homotypic alignment (chimeric alignments where arms overlap). Four replicates of sequencing data shown. **d-e**, Comparison of U2, U4 and U6 snRNA structures and interactions in 0.5mg/ml AMT (d) and 0.5mg/mL amotosalen (e) crosslinked sample. Total RNA sequencing data of HEK293_AMT_rep1 and HEK293_Amoto_rep1 were showed. The number of total input reads was 1.5 and 2.0 million, respectively. DG1-6: U2 alternative structures; DG7-9: U4 alternative structures; DG10-11: U6 alternative structures; DG12-14: U4-U6 interactions; DG15-18: U2-U6 interactions. The structure and interaction model showed high concordance between AMT and amotosalen crosslinking.

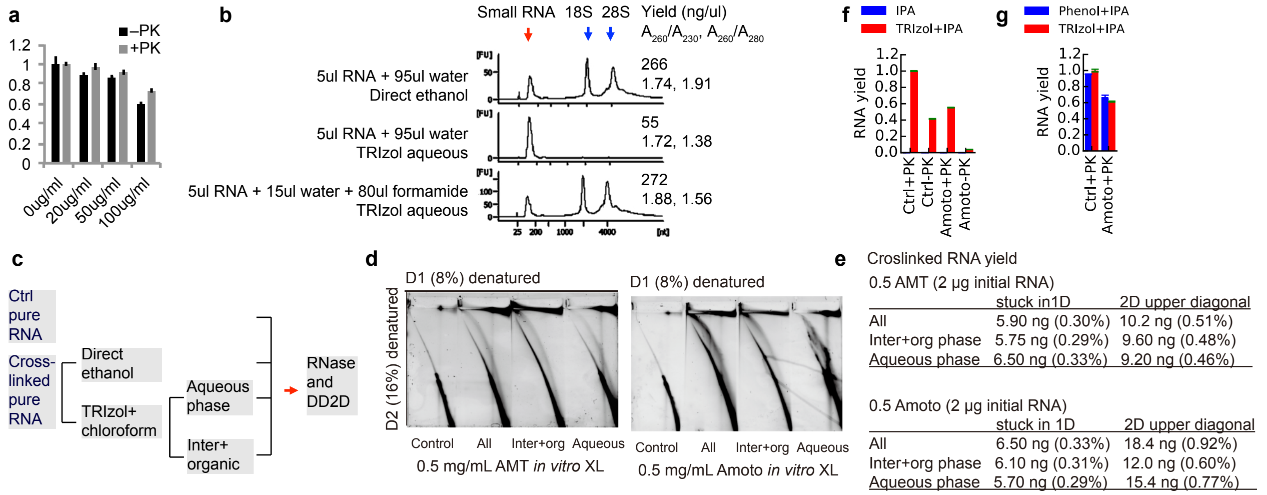

**Supplementary Figure 3. TNA: A new method to recover crosslinked RNA from cells. a**, Psoralen crosslinking reduces yield of TRIzol extracted RNA. AMT crosslinking was performed in triplicate for four conditions: without AMT, with AMT at 20, 50 and 100µg/ml, each with 30min UV 365nm irradiation in 12-well plate, 0.5ml Drosophila S2 cells per well, at 20million/ml. After irradiation, cells were collected in 0.1% SDS (200µl). Seventy five percentage of each lysate was treated with proteinase K (PK) while the other 25% was not. PK treatment condition: 20mM Tris·Cl; 10mM EDTA; 1% SDS and 0.1mg/ml proteinase K, incubating at 40ºC for 40min. After PK treatment, RNA was extracted from the cells using the standard TRIzol method, quantified by Nanodrop, and normalized against the non-crosslinked samples. **b**, Phase partition of *in vitro* crosslinked RNA in TRIzol+chloroform with or without formamide. Five µg total HEK293 RNA was *in vitro* crosslinked with 0.5mg/ml AMT for 30min. After crosslinking, RNA was either directly precipitated with ethanol, or supplemented with water or formamide and then purified with TRIzol. Each 5µl RNA sample in PBS was supplemented with 95µl water, or 15ul water plus 80µl formamide. For TRIzol purifications, 100µl TRIzol and 50µl chloroform were added. After ethanol precipitation, RNA was resuspended in 20µl water and quantified by Nanodrop and Bioanalyzer. **c**, Experimental design to test how much crosslinked RNA was in the aqueous *vs.* inter+organic phase. As a control, total noncrosslinked RNA was used. Alternatively, crosslinked pure RNA was either directly precipitated with ethanol or purified using TRIzol+chloroform. All RNA samples were digested with RNase III and subjected to DD2D gel purification. **d**, 2D gel purification of crosslinked RNA from noncrosslinked, all crosslinked, crosslinked in the inter+organic phase, croslinked in the aqueous phase by standard Trizol purification. 2µg RNA samples were crosslinked with 0.5mg/ml AMT (left panel) or 0.5mg/ml amotosalen (right panel). **e**, Recovery of crosslinked RNA from the 1D-stuck and 2D-upper diagonal were quantified. Total yield was similar for the three crosslinked samples with identical input amount. **f**, Total nucleic acids can be purified after PK treatment and alcohol precipitation in presence of TRIzol. Each tube of HEK293T cell pellet from a 10-cm plate was resuspended in 100µl 6M GuSCN, shaken vigorously, then diluted to 600µl in 1x PBS, 10mM EDTA and passed through a 26G needle 20 times to reduce viscosity. Each 300µl lysate was added PK to 1mg/ml and incubated at 37ºC for 60min. After the digestion, samples were directly precipitated using 1 volume isopropanol (300µl), or supplemented with 150ul Trizol and then precipitated with 1 equivalent volume isopropanol (450µl). The samples treated with TRIzol + isopropanol resulted in precipitates becoming water soluble, albeit with lower yield in the crosslinked samples. **g**, Precipitation of total nucleic acids using isopropanol and phenol, after PK treatment. One tube each of control and amotosalen crosslinked cells was lysed in 100µl 6M GuSCN to make a homogenous solution, then diluted to 600µl with 10x PBS and water; the final solution contained 1M GuSCN, 1x PBS, 10mM EDTA and 1mg/ml PK. The dilution often lead to some insoluble material. Samples were passed through 26G needles 20 times to reduce viscosity, and then incubated at 37ºC in a thermomixer for 1 hour. Then 60µl of each samples was taken out for precipitation as follows: 60µl + 6µl NaOAc 3M pH 5.0 + 60µl phenol + 120µl isopropanol (addition of isopropanol produced obvious stringy precipitate), or 60µl + 6µl NaOAc 3M pH 5.0 + 120µl Trizol LS + 180µl isopropanol (no obvious stringy material). Yields of the both methods were similar, suggesting that phenol was the primary component necessary in keeping residual proteins in solution, while alcohol precipitated nucleic acids.

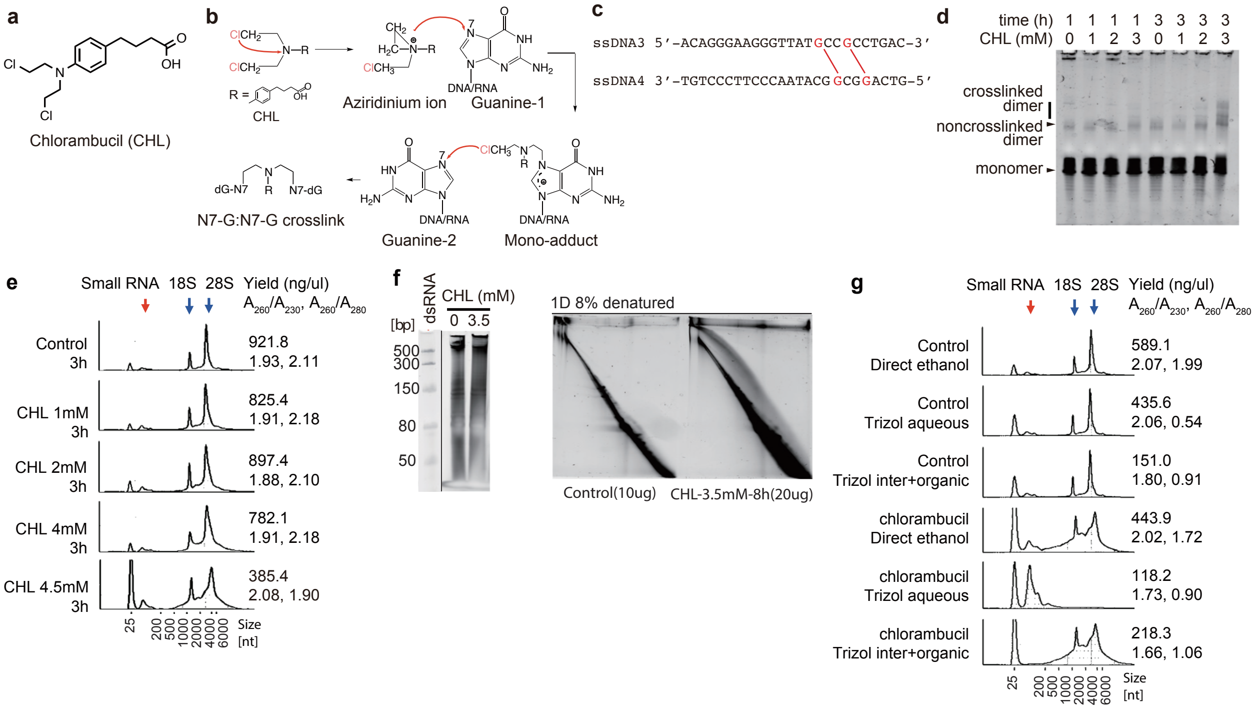

**Supplementary Figure 4. Chlorambucil crosslinks RNA and makes RNA more hydrophobic. a,** Chemical structure of chlorambucil (CHL). **b**, The mechanism of chlorambucil crosslinking of nucleic acids. Chlorambucil, like many other nitrogen mustards, forms aziridinium ions by intramolecular displacement of the chloride by the amine nitrogen. This aziridinium group then alkylates DNA once it is attacked by the N-7 nucleophilic center on the guanine base to become a mono-adduct. A second attack after the displacement of the second chlorine results in the formation of interstrand cross-links. **c**, Synthetic 25-mer DNA duplex used to test CHL crosslinking.The red lines indicate predicted guanine-guanine DNA interstrand cross-link selectively formed by chlorambucil at the 5'-GGC sequence (an 1,3 cross-link, G:G3). **d**,15% gel electrophoresis of 25-mer DNA oligos crosslinked with different concentrations of CHL at various times. Equal amounts of complementary RNA oligoes were mixed together and denatured at 72°C for 3min, cooled down at room temperature for 5min, then were incubated in a final solution of 1x PBS, 1mM EDTA, with CHL at 0mM, 1mM, 2mM, 3mM respectively, for either 1h or 3h. **e**, TapeStation electropherograms of crosslinked total RNA by different concentrations of CHL. 17µg of purified total RNA from HEK293 cells was incubated with 0mM, 1mM, 2mM, 4mM, 4.5mM CHL respectively, containing 100mM Tris, 1mM EDTA, at 37°C in the dark for 3h. Each sample was precipitated with 3 times volume of ethanol. **f**, DD2D gel system showing CHL crosslinked total RNA. 20µg of purified total RNA from HEK293 cells was crosslinked by 3.5mM CHL in 100mM Tris and 1mM EDTA buffer for 8h, 10ug noncrosslinked RNA served as control. After fragmentation by RNase III, crosslinked RNA fragments were separated by DD2D gel. The red outlined area above the diagonal indicated crosslinked RNA. **g**, CHL crosslinked total RNA partitions into the interphase during TRIzol extraction. 12µg purified total RNA from HEK293 cells was incubated with or without 4.8mM CHL for 6hr. After crosslinking, each sample was divided equally into two tubes, one for direct ethanol precipitation, another for TRIzol extraction, which was further divided into aqueous phase and inter+organic phase. RNA in aqueous phase and inter+organic phase were precipitated by ethanol. RNA profile of each condition was visualized using TapeStation.

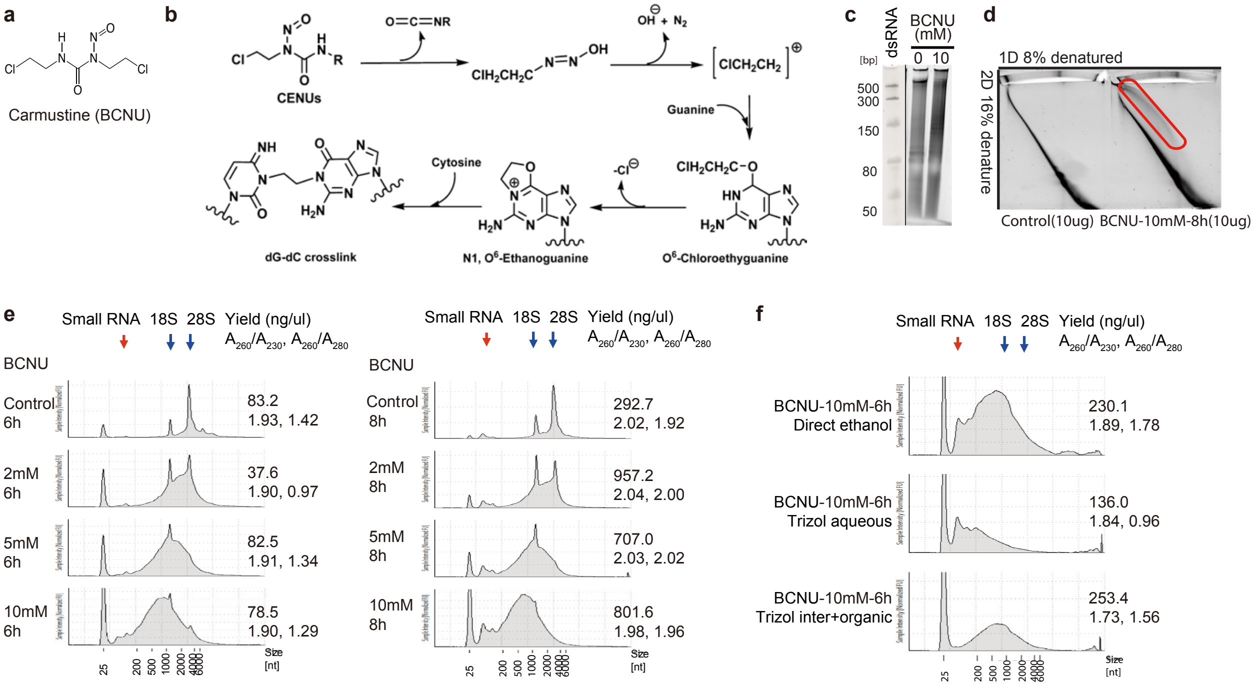

**Supplementary Figure 5. BCNU crosslinks RNA and makes it more hydrophobic**. **a**, Chemical structure of carmustine (BCNU). **b**, Molecular mechanisms of chloroethylnitrosoureas (CENUs) crosslinking G-C base pairs. BCNU is one of the CENUs, where R=$CH_2CH_2Cl$. The chloroethyl diazonium ion produced by the decomposition of CENUsalkylates guanine on the O6 site formed O6-chloroethylguanine (O6-ClEt-Gua), followed by further alkylation of the complementary cytidine on the N3 site via a cationic intermediate, N1,O6-ethanoguanine. **c-d**, DD2D gel showing BCNU crosslinked total RNA. 10ug purified total RNA from HEK293T cells was incubated with or without 10mM BCNU in 100mM Tris, 1mM EDTA, and in the dark for 8 hours. RNA samples were then purified, fragmented with RNase III and run on the DD2D gels. **e**, TapeStation profiles of BCNU crosslinked total RNA under different conditions. Pure RNA from HEK293T cells was crosslinked with 2, 5 or 10 mM BCNU at 37°C in the dark for either 6 or 8 hours. Long-term crosslinking induced partial RNA degradation. **f**, TapeStation profiles of different phases from TRIzol extracted BCNU crosslinked total RNA. Pure RNA was crosslinked with 10mM BCNU for 6 hours. After crosslinking, sample was divided equally into 2 tubes, one for the direct ethanol precipitation, another for TRIzol extraction, which were further divided into the aqueous phase and the inter+organic phase. RNA in two phases were precipitated by ethanol. Larger RNA partitioned to inter+organic phase from aqueous phase.

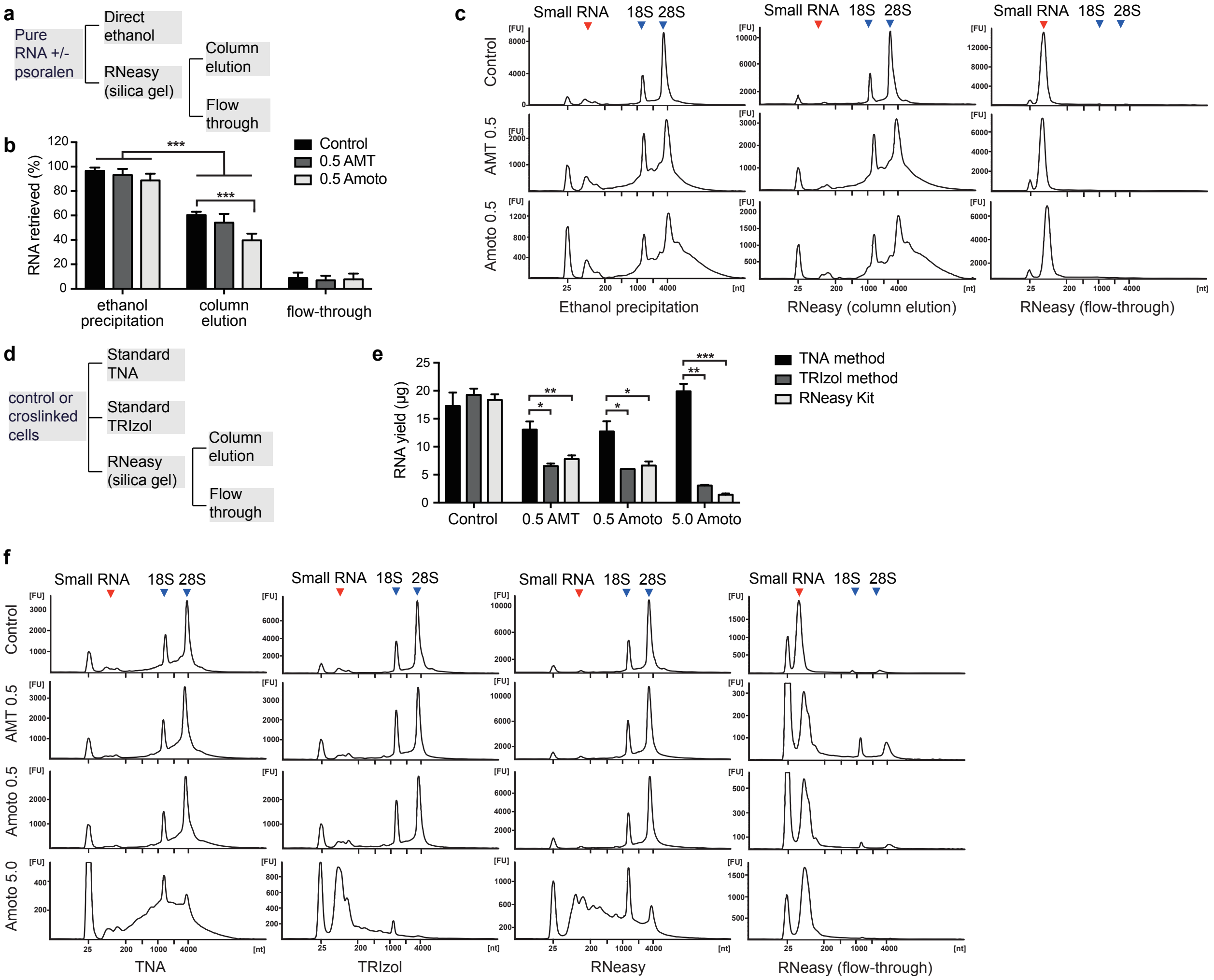

**Supplementary Figure 6. Comparison among TRIzol, TNA and RNeasy in recovering crosslinked RNA from *in vitro* or *in vivo* sources. a**, Diagram for the experimental design. Purified total RNA was used directly or crosslinked with 0.5mg/ml AMT or amotosalen. Then RNA is directly precipitated out of solution or purified using the RNeasy kit. Flow-through fraction was also precipitated using ethanol. **b**, Quantity of RNA recovered ratio from 5µg of control and *in vitro* crosslinked total RNA. *** P-value < 0.001. **c**, Size distribution of RNA after purification using direct ethanol precipitation or the RNeasy kit. Profiles were obtained using Tapestation. Here small RNA includes RNA in the range of 50-300nt, such as tRNAs, snRNAs and snoRNAs. The smear, especially the tail after the 28S peak was indicative of successful crosslinking. **d**, Diagram for the experimental design testing various methods in extracting crosslinked RNA from cells. Cells were used as is or crosslinked using various psoralens and concentrations (0.5 or 5 mg/ml). Then RNA was extracted using one of the three methods. **e**, Quantity of RNA isolated from paired control and AMT/amotosalen crosslinked two million cells. * P value < 0.05, ** P-value < 0.01, *** P-value < 0.001. For the TNA method, DNA was removed before quantification. TNA consistently outperforms other methods in the extraction of crosslinked RNA. **f**, Size distribution of RNA after purification using different methods.

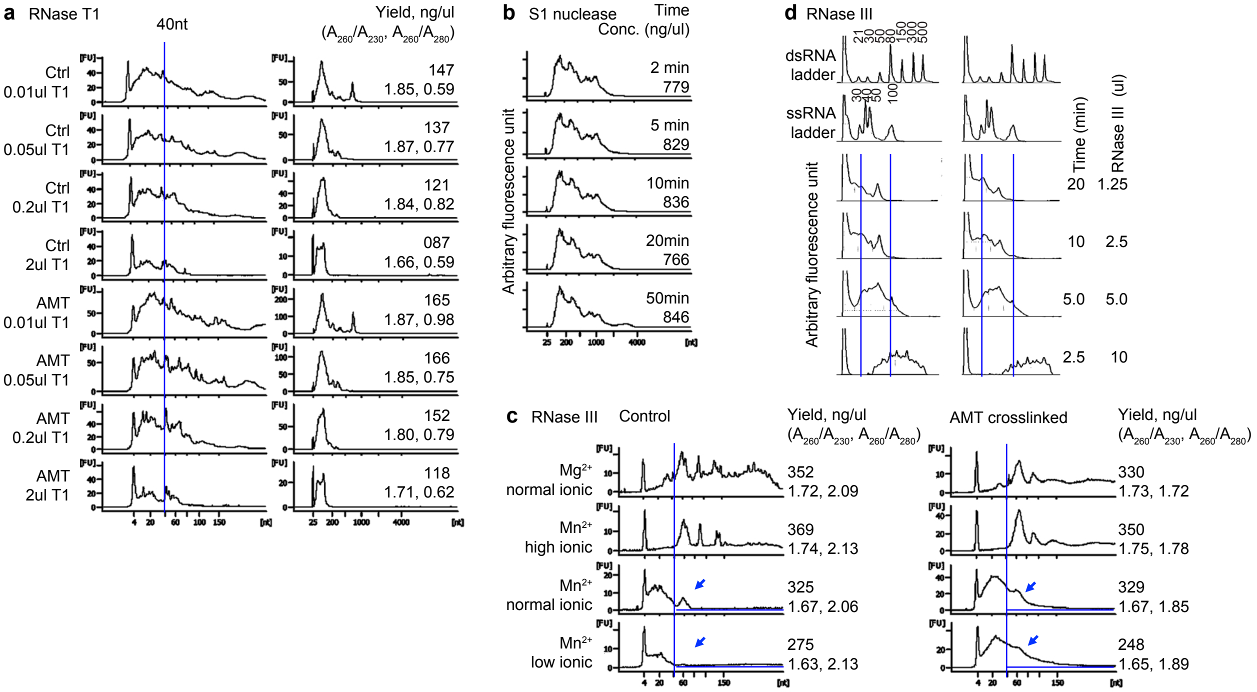

**Supplementary Figure 7. Optimization of nuclease fragmentation of crosslinked RNA.** **a**, RNase T1 treatment reduces total RNA amount and size, but the distribution is still broad. HeLa cells were crosslinked with 0.5mg/ml AMT and 365nm UV in Stratalinker for 30min. RNA was extracted from control and crosslinked cells using the S1/PK method (Lu et al., 2016). RNase T1 buffer (10x) contains 3M NaCl, 100mM Tri-HCl, pH7.0, 20mM EDTA. 4μg S1/PK extracted control and AMT crosslinked RNA was incubated with RNase T1 (Fermentas, EN0541, 1000U/μl) in 40μl total volume, at 22-24°C for 30min. RNA was purified using TRIzol and analyzed by Nanodrop and Bioanalyzer. **b**, Extensive S1/PK digestion does not reduce size or yield significantly. One 10cm plate of cells was crosslinked with 0.5mg/ml AMT and then split into 5 samples for S1/PK digestion. S1 digestion time was varied from 2 to 50 min. Samples were quantified using Nanodrop and Bioanalyzer. **c**, AMT crosslinking affects RNase III digestion. Total HEK293 RNA was crosslinked *in vitro* at 1μg/μl, with 0.5mg/ml AMT and 365nm UV light in Stratalinker for 30min. Non-crosslinked and crosslinked RNA samples were digested with RNase III at 37°C for 20min; each sample contained buffer + 2μl RNase III + 10μg RNA, in a total volume of 20μl. Buffer conditions were as follows: $Mg^{2+}$ normal ionic strength: 50mM Tris-HCl, 1mM DTT, 50mM NaCl, pH 7.5 at 25°C + 5 mM $MgCl_2$. $Mn^{2+}$ low ionic: 5mM Tris-HCl, 1mM DTT, pH 8.0 at 25°C + 5mM $MnCl_2$. $Mn^{2+}$ normal ionic: 50mM Tris-HCl, 1mM DTT, 50mM NaCl, pH 7.5 at 25°C + 5 mM $MnCl_2$. $Mn^{2+}$ high ionic: 50mM Tris-HCl, 1mM DTT, 1M NaCl, pH 7.5 at 25°C + 5mM $MnCl_2$. After precipitation, samples were profiled using Bioanalyzer. The blue arrows point to the differences induced by crosslinking, which is more obvious after RNase III digestion under lower ionic strength. **d**, RNase III reaction kinetics affect RNA fragment size. Reaction conditions: 5μg RNA sample from the TNA method was treated with variable amount of RNase III in low ionic strength buffer for various times. The ssRNA ladder and samples were denatured before loading into TapeStation, whereas the dsRNA ladder was not.

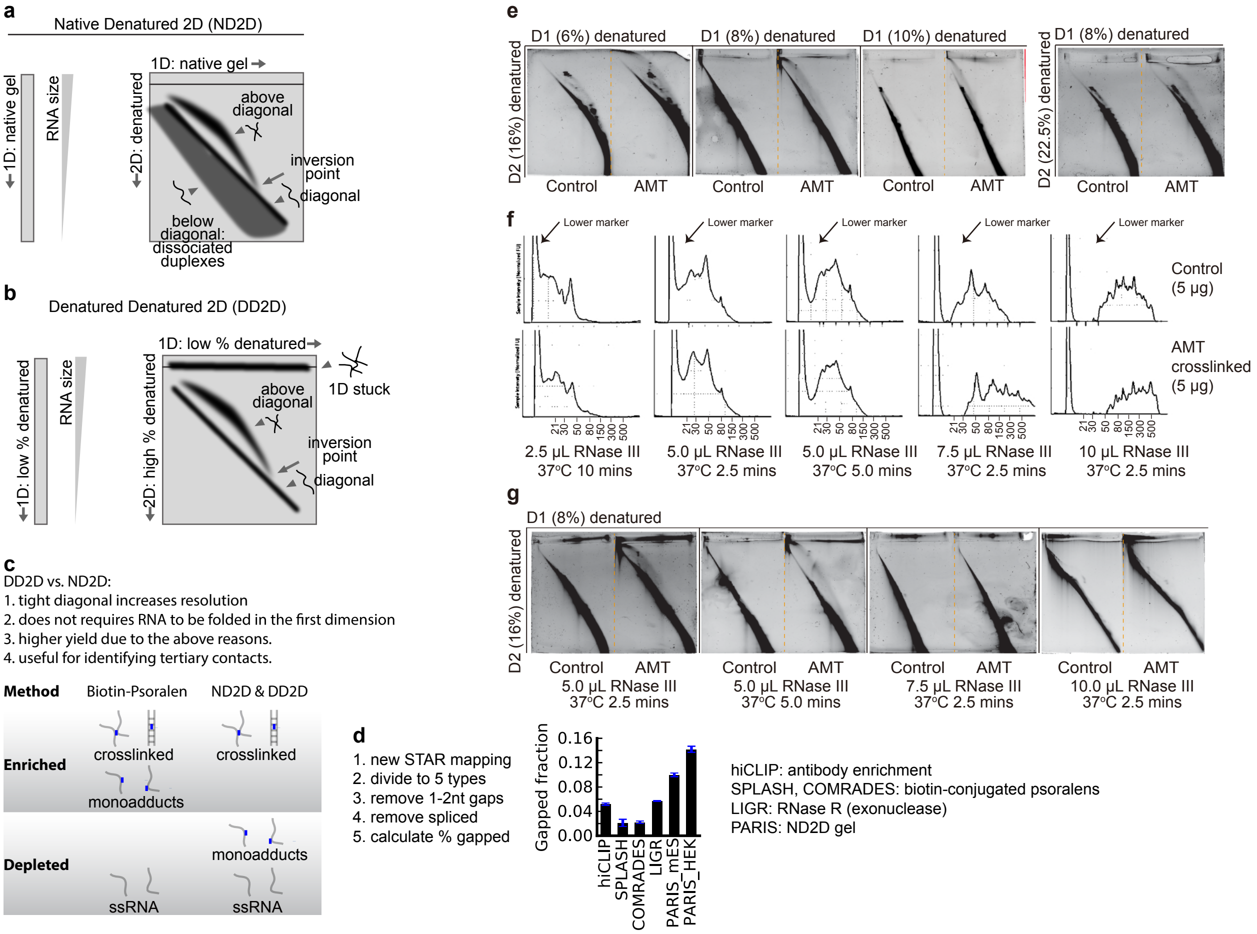

**Supplementary Figure 8. Optimization of the DD2D gel method. a,** Diagram for the ND2D gel system. In the first dimension, RNA separates roughly based on size. In the second dimension, RNA further separates based on shape where crosslinked fragments migrate much slower due to much larger hydrated radius. Inversion point is at an RNA size where the increase in hydration radius is no longer contributing to the migration. Noncrosslinked RNA duplexes that run together in the native first dimension would dissociate in the second dimension, causing downward smear in the second dimension. **b**, Diagram for the DD2D gel system. Like the ND2D system, RNA separates based on size in the first dimension. Here the shape also contributes to the migration due to denaturation, but the lower gel density (bigger pore size) reduces the effect of the shape of crosslinked RNA. In the second dimension, the higher gel density (smaller pore size) increases friction for the crosslinked RNA fragments, substantially more than in noncrosslinked ones. The RNA stuck in the first dimension may be more structured RNA that could not enter the second dimension of higher percentage gels. **c**, Comparison of ND2D and DD2D gels. It is worth noting that the DD2D gel does not require crosslinked RNA to be strongly base paired and is therefore also useful for identifying crosslinkable tertiary contacts. **d,** PARIS outperforms other methods in terms of percentage of gapped alignments. Fraction of gapped alignments for various methods were calculated as (gapped alignments + RNA-RNA interactions) / total alignments. Short indels are discarded since they are often due to sequencing errors, which are especially high in the presence of psoralen adducts. The following data were used. PARIS (HEK293, Lu et al. 2016), PARIS (mES, Lu et al. 2016), LIGR (Sharma et al. 2016), SPLASH (Aw et al. 2016), COMRADES (Ziv et al. 2018). **e**, Testing combinations of various gel concentrations for the first and second dimensions. **f**, Analysis of the RNase III digested total RNA with different shortcut conditions. **g**. DD2D purification of the crosslinked RNA with different RNase III conditions.

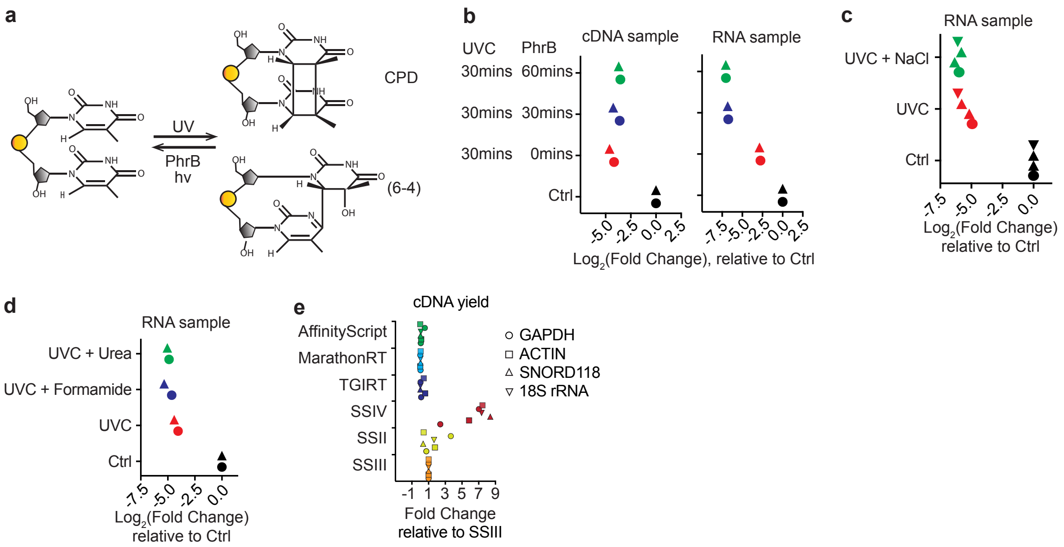

**Supplementary Figure 9. Effects of various condtions on the repair, prevention or bypass of UVC damages on RNA. a**, Funtion of photolyase in DNA damage repair. UV irradiation produces heavy DNA damages, such as cyclobutane pyrimidine dimer (CPD) and (6-4) lesion. These DNA damages can be repaired by the enzyme photolyase (PhrB) in the presence of long-wavelength light (e.g. UVA and blue). **b**, PhrB cannot repair UVC damaged RNA. UVC damage repair efficiency showed by Ct value obtained by qPCR. cDNA sample is generated from total RNA of HEK293T cells by SSIII with random hexamer. cDNA and RNA sample is irradiated by 254 nm UV for 30 mins to introduce damage. Then the enzyme PhrB is used to repair UVC damage under 365 nm UV for 30 and 60 mins respectively. Log2-fold change values determined by qPCR and normalized to control sample. Two replicates are shown for each condition. **c**, High salt reverse crosslinking condition with UVC. 1 M NaCl was added to reversal reaction, to help RNA form stable secondary structures. UVC 254nm 30min damage was tested by cDNA yield of ACTB. Log2-fold change values determined by qPCR and normalized to control sample. **d**, UVC reversal of crosslinking with denaturing buffer, such as 50% formamide or 4 M Urea. UVC 254nm 30min damage was tested by cDNA yield of ACTB mRNA. Log2-fold change values determined by qPCR and normalized to control sample. **e**, Bypass UVC damages by different reverse transcriptases. cDNA yield of GAPDH (circles), beta-ACTIN (square), SNORD118 (up triangle) and 18s rRNA (down triangle) were tested to show the bypass by reverse transcriptases after UVC damage. UVC damage is introduced by 254 nm UV irradiated RNA for 30 mins. cDNA yield was determined as the Ct value obtained by qPCR of the reverse transcriptional cDNA and normalized to a Superscript III condition. Standard buffers and reaction conditions were used unless otherwise indicated.

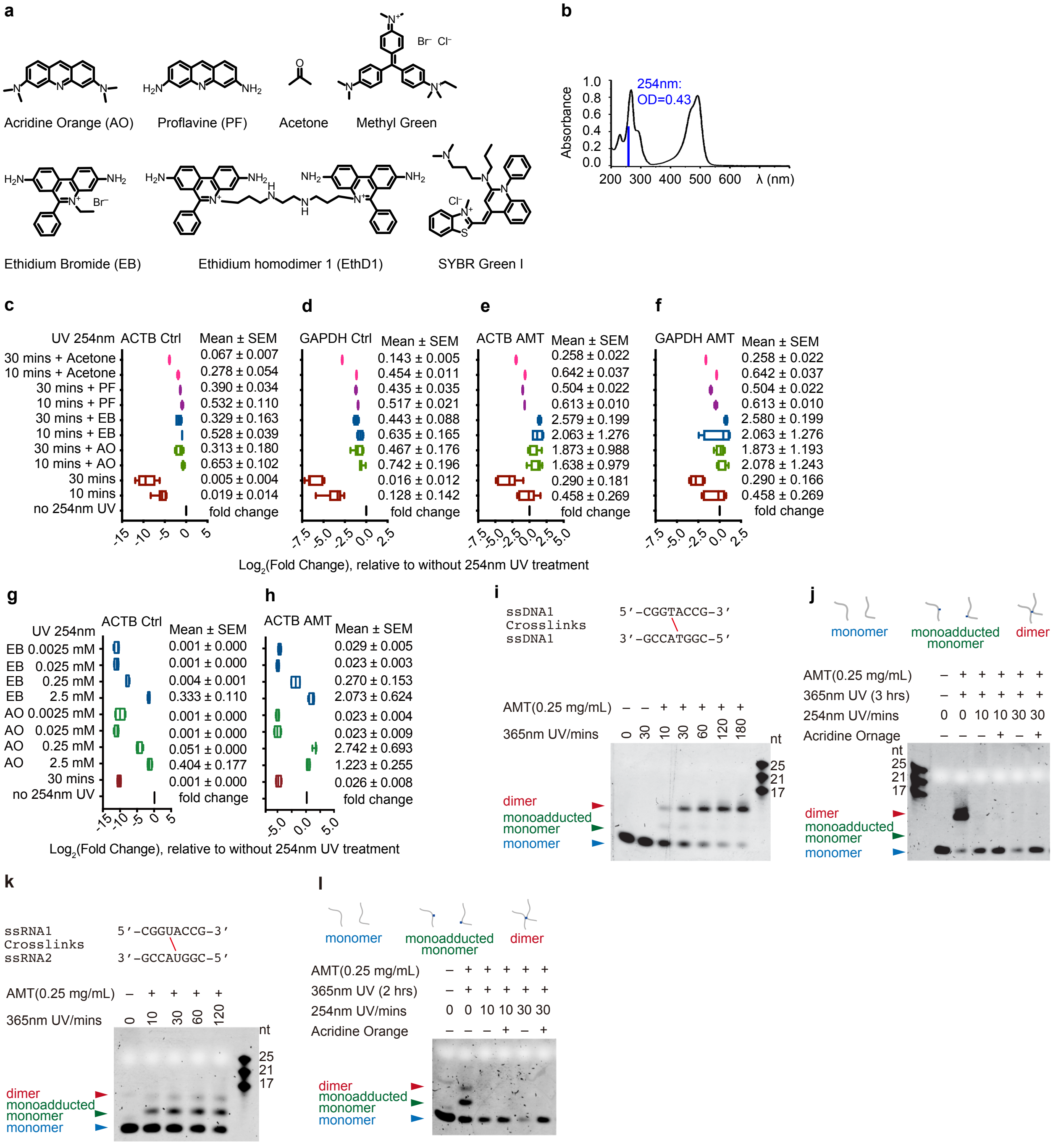

**Supplementary Figure 10. Singlet quenchers prevent UVC induced RNA damage without inhibiting reversal of psoralen crosslinks. a**, Structures of singlet state quenchers used to protect RNA from UVC damages. The structure of SYBR Gold is proprietary so SYBR Green I is shown instead. **b**, UV absorbance of acridine orange; OD=0.43 at 254nm (Selvaggi et al. 2015, Applied Catalysis B: Environmental). **c-f**, cDNA synthesis yield of ACTB and GAPDH mRNAs with or without different dimer inhibitors during 254nm photo-reversal. Log2-fold change values determined by qPCR and normalized to non-photo-reversal samples. 254nm UV 10 mins and 30 mins were tested. (AO) Acridine Orange, 2.5 mM; (EB) Ethidium Bromide, 2.5 mM; (PF) Proflavine 25 mM; Acetone, 50%. **g-h**, cDNA synthesis yield for ACTB mRNA with or without different dose of dimer inhibitors during photo-reversal step. Log2-fold change values determined by qPCR and normalized to non-photo-reversal sample. 254nm UV 30 mins were applied for photo-reversal condition. **i**, Photo crosslinking of 8-mer DNA oligos at 365 nm UV. Crosslinking times were 10, 30, 60, 120 and 180 mins, respectively. Crosslinked products of 180 mins were used to test photo-reversal condition. **j**, AO did not affect the photo-reversal efficiency of DNA oligo dimers. Crosslinked DNA oligo dimers were reversed by 254 nm UV with or without the protection of AO. The reverse crosslinking times were 10 and 30 mins, respectively. **k**, Photo crosslinking of 8-mer RNA oligos at 365 nm UV. The crosslinking time is 10, 30, 60 and 120 mins, respectively. Most RNA oligos formed monoadducts. Crosslinked products of 120 mins were used to test photo-reversal condition. **l**, AO did not affect the photo-reversal efficiency of RNA oligo dimers and monoadducted monomer. Photo-reversal of crosslink was performed at 254 nm UV. AO was added to inhibit pyrimidine photoproducts. The reverse crosslinking times were 10 and 30 mins, respectively.

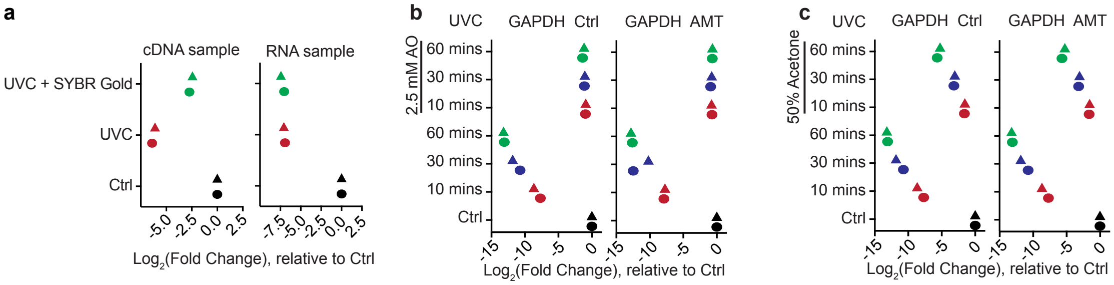

**Supplementary Figure 11. Singlet quenchers prevent UVC induced DNA damage. a**, SYBR Gold partially protects DNA against UVC damage. UVC damage was quantified by qRT-PCR on the UVC treated cDNA or mRNA. cDNA sample is generated from total RNA of HEK293T cells by SSIII. UVC, 254 nm UV for 30 mins. SYBR Gold can partially block UVC damage for cDNA sample (left side), but not RNA sample (right side). **b**, Acridine orange prevents DNA sample from UVA damages. 300 ng of control and 0.5 mg/ml AMT crosslinked DNA were irradiated by 254 nm UV for 10 mins, 30 mins and 60 mins. 2.5 mM acridine orange was used to protect DNA. UVC damages was quantified by qRT-PCR on the GAPDH mRNA. Log2-fold change values determined and normalized to control sample. **c**, Acetone partly block UVC damages on DNA sample. 300 ng of control and 0.5 mg/ml AMT crosslinked DNA are used for testing. 50% of acetone was added to protect DNA sample.

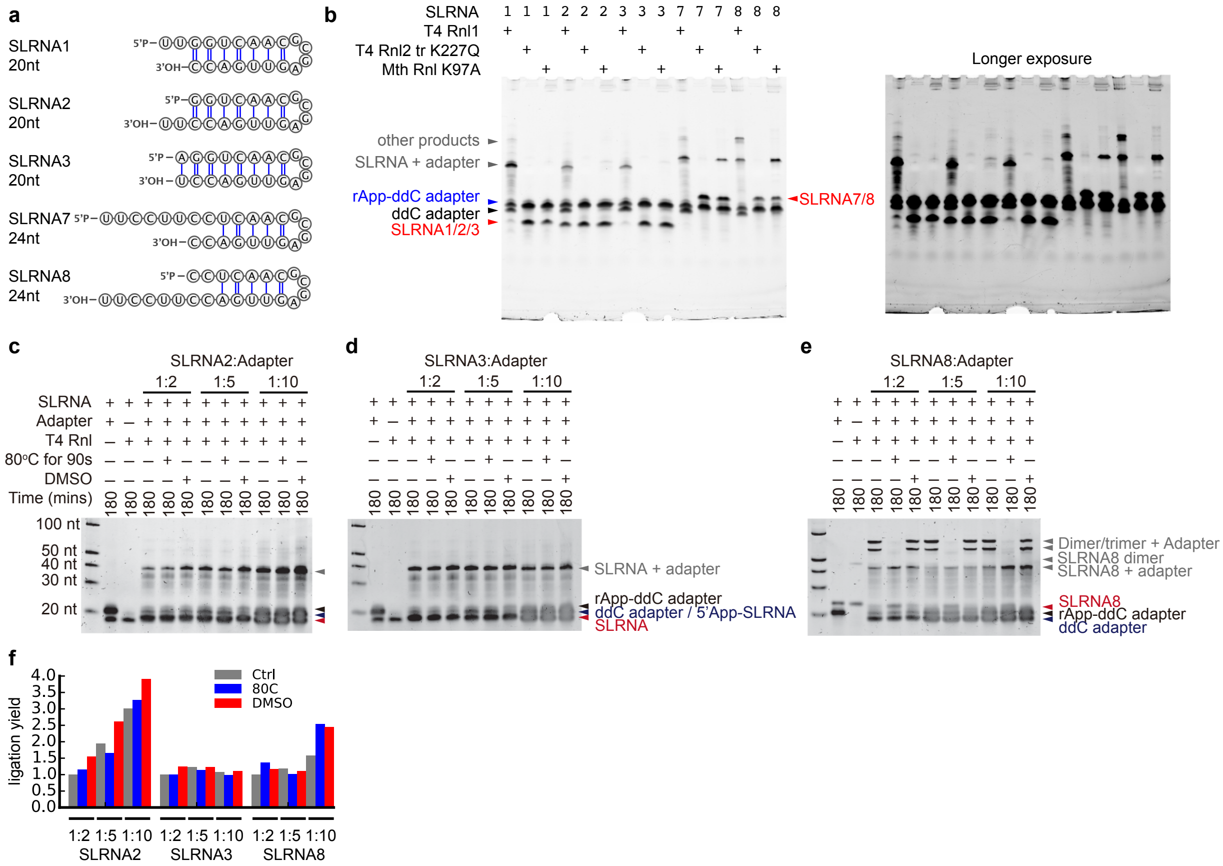

**Supplementary Figure 12. Systematic optimization of adapter ligation. a**, Sequences and secondary structures of synthetic stemloop RNA (SLRNA) oligos used to test ligation conditions. The various stemloop structures were designed to test the efficiency of ligating to structure RNAs. **b**, Electrophoretic gel showing the adapter ligation efficiency of T4 Rnl, T4 Rnl2 tr K227Q and Mth Rnl K97A, respectively. Tested ligation of SLRNA oligos to adenylated ssDNA adapters. Exposure: left panel 2 sec, right panel 10 sec. **c-e**, Denaturing conditions and abundant adapter increase adapter ligation efficiency. 5pmole SLRNA2/3/8 and 10/25/50 pmole rApp-ddC Adapter were used ligagted by T4 Rnl at room temperature for 3hour. Denaturing treatment: SLRNA 2/3/8 was incubated at 80°C for 90 seconds and snap cooled on ice for at least 1min; DMSO treatment, 10% (v/v) DMSO was added to sample. **f**, Quantified ligation efficiency for panels c-e.

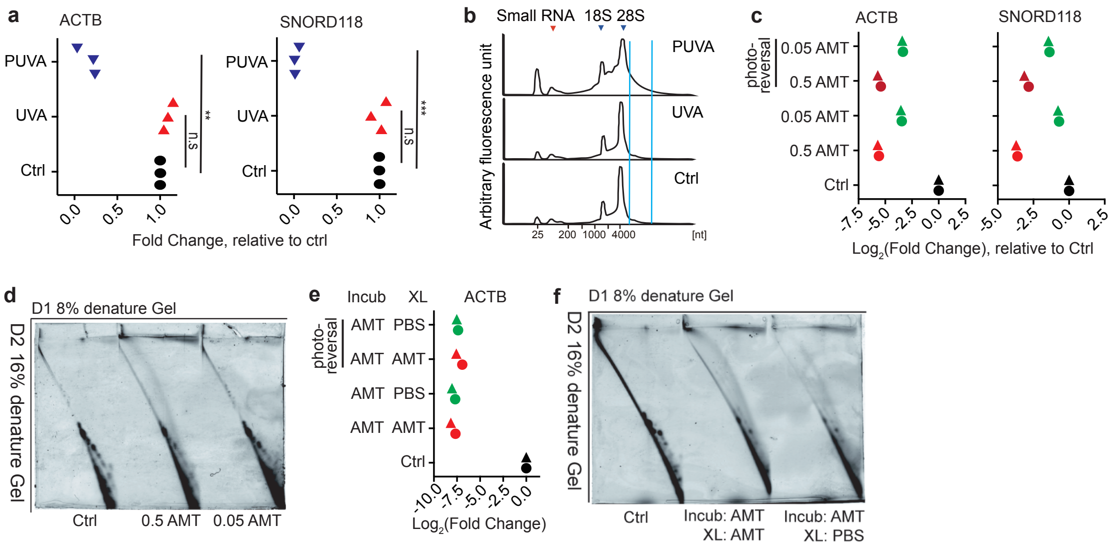

**Supplementary Figure 13. PUVA cause RNA damages.** **a**, PUVA, not UVA, will induce RNA damge. cDNA synthesis yield of ACTB and SNORD118 are tested to study the RNA damage. UVA, 365 nm UV irradiated RNA sample for 30 mins; PUVA, AMT 365 nm UV irradiated RNA sample 30 mins with 0.5 mg/ml; Three replicates are shown for each condition. ** $p < 0.01$, *** $p < 0.001$. **b**, RNA profile after UVA and PUVA treatment. UVA alone did not affect RNA interity. PUVA will induce RNA crosslinking (within blue two lines). **c**, PUVA damage on RNA is related with the concentration of AMT. 0.5 mg/ml and 0.05 mg/ml AMT are used to introduce PUVA damage. The lower concentration of AMT, the less PUVA damages on RNA. PUVA damage still remained after reverse crosslinking by 254 nm UV with the protection of arcidinr orange. cDNA synthesis of ACTB and SNORD118 are used to test PUVA damage. Two replicates are shown for each condition. **d**, 2D gel showing the crosslinking efficiency of different concentration of AMT. Less AMT will reduce the crosslinking efficiency. 0.5 mg/ml AMT is necessary for high efficient crosslinking. **e**, Incubation with AMT solution and crosslinking with PBS solution do not reduce PUVA damage. Incubation step (Incub): HEK293T cells were incubated in 0.5 mg/ml AMT solution for 15 mins, to make sure AMT penetrate into RNA duplex. Crosslinking step (XL): after incubation, HEK293T cells are irradiated by 365 nm UV for 30 mins in either the same AMT solution or PBS solution. **f**, Incubation with AMT solution and crosslinking with PBS solution reduce the crosslinking efficiency.

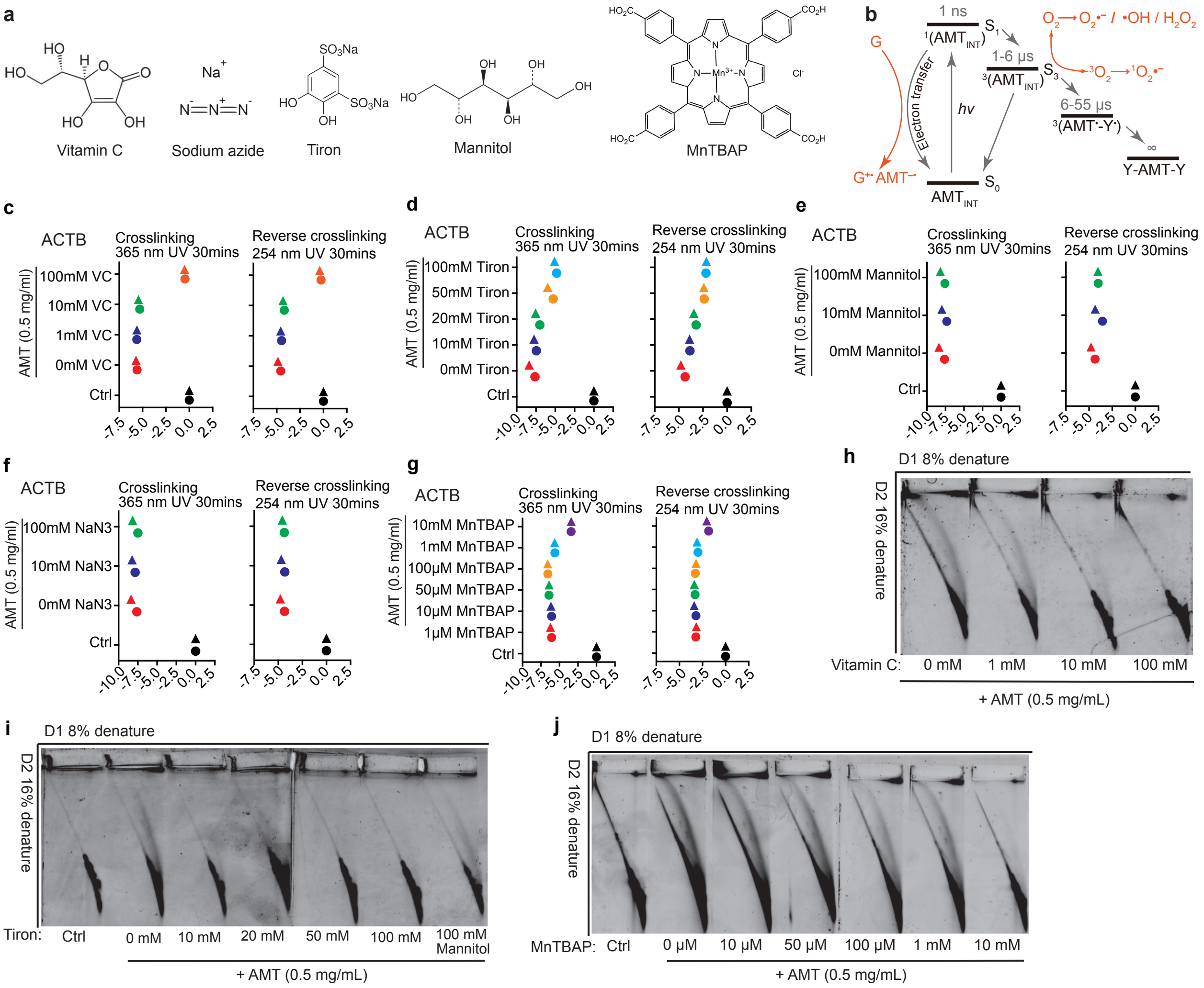

**Supplementary Figure 14. Scavengers prevent PUVA damage but also block crosslinking. a**, Structures of oxidant scavengers tested in this study. Antioxidants: vitamin C (VC, electron donor); $O_2$•- scavenger: Tiron and MnTBAP; •OH scavenger: Mannitol; $^1O_2$ scavenger: $NaN_3$. **b**, An overview of two types of photosensitized RNA damages. Type I, photo-induced electron transfer mechanism. Guanine radical cation forms through the electron transfer reacts with singlet state of AMT, leading to the formation of the oxidized products of guanine. Type II, generation of reactive oxygen species mechanism. Different types of oxygen species are generated from the triplet state AMT. $O_2^{\bullet-}$, superoxide anion radical; •OH, hydroxyl radical; $H_2O_2$, hydrogen peroxide; $^1O_2$, singlet oxygen. Guanin has lowest oxidation pontential among the four bases are the most frequently oxidized. Less frequent damages, such as hydrates and strand breaks are not shown here. The direct involvement of electron transfer in guanine damage is similar to the pyrimidine crosslinking process, both of which can be quenched by antioxidants. **c-g**, PUVA induced damage in RNA are prevented by some scavengers based on analysis of cDNA synthesis yield from ACTB mRNA. HEK293T cells in 6 well plates are crosslinked by 0.5 mg/ml AMT in the presence of different scavengers. After crosslinking by 365 nm UV for 30 mins, total RNA extracted using the TNA method are used to perform RT-qPCR. Log2-fold change values were determined by qPCR and normalized to control samples. Reverse crosslinked samples by 254nm UV 30 mins are also tested. High concentration of VC (c), Tiron (d) and MnTBAP (g) can reduce PUVA damage. Mannitol (e) and $NaN_3$ (f) have no effects on PUVA damage. **h-j**, 2D gel showing the crosslinking efficiency of HEK293T cells by 0.5 mg/ml AMT in the presence of different scavengers. High concentration of VC (h), Tiron (i) and MnTBAP (j) blocked crosslinking, while mannitol did not (i, last panel).

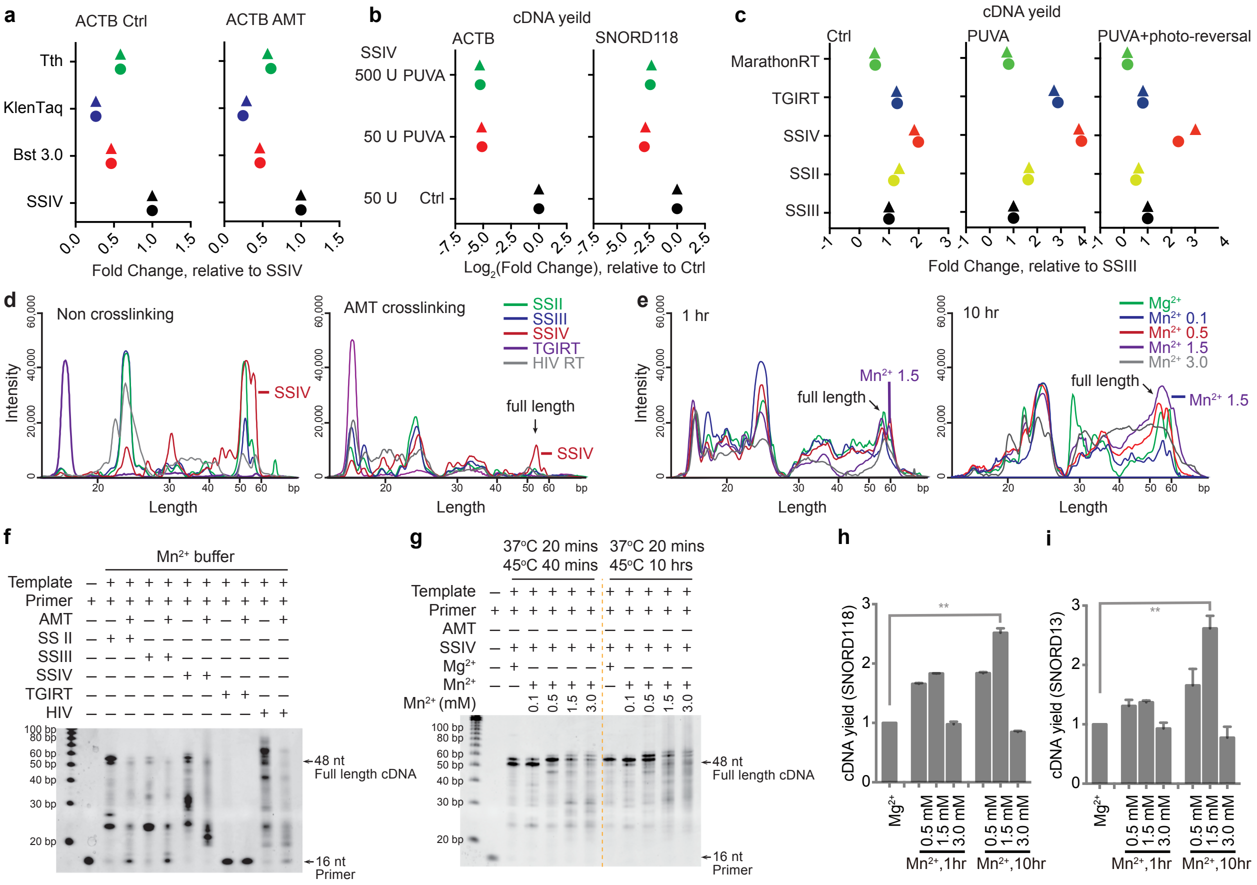

**Supplementary Figure 15. Bypass of PUVA induced oxidative damage of RNA. a**, DNA polymerases with reverse transcriptase activity cannot bypass PUVA damages. Bypass ability is shown based on cDNA synthesis yield of ACTB. Gene specific primer targeting ACTB (AGCACTGTGTTGGCGTACAG) is used for reverse transcription. Tth DNA polymerase, KlenTaq DNA polymerase and Bst 3.0 DNA polymerase were tested and compared to SuperScript IV (SSIV). Reverse transcription step is performed according the standard manual of each enzyme. PUVA damage is induced by 0.5 mg/ml AMT crosslinking under 365nm UV for 30 mins. Two replicates are shown for each condition. **b**, Reverse transcriptase units with the PUVA damage on RNA. 30 ng of control and PUVA damaged RNA are used to test the by pass ability of SuperScript IV (SSIV). Reverse transcription are performed according to standard manual of SSIV. **c**, The effects of different reverse transcriptases on PUVA damage. cDNA yield of 18S rRNA were tested. PUVA, HEK293T cells were crosslinked by 365 nm UV for 30 mins with 0.5 mg/ml AMT. PUVA+photo-reversal, PUVA sample plus 254 nm UV reversal with the protection of acridine orange. cDNA yield was determined as the Ct value obtained by qPCR of the reverse transcriptional cDNA and normalized to a Superscript III condition. Standard buffers and reaction conditions were used here. **d-e**, Quantifying the gel pictures in Figure 4f-g. The intensity profile for each gel lane was extracted using iBright Analysis Software (Invitrogen). Background was substracted by using a rolling-ball algorithm with 100 μm radius to estimate the amount of background at each position. Pixel positions were converted to DNA length by interpolating the 10 bp DNA ladder against pixel position. **f**, Analysis of primer extension products synthesized by different reverse transcriptases in manganese buffer. **g**,The effect of $Mg^{2+}$ and $Mn^{2+}$ buffer on full-length of cDNA synthesis by SSIV. Initial RNA is non-crosslinked. **h-i**, cDNA yield of two snoRNAs SNORD118 and SONRD13, using SSIV in different reaction buffers and different incubation time. Normalized to a standard $Mg^{2+}$ buffer. ** p< 0.01.

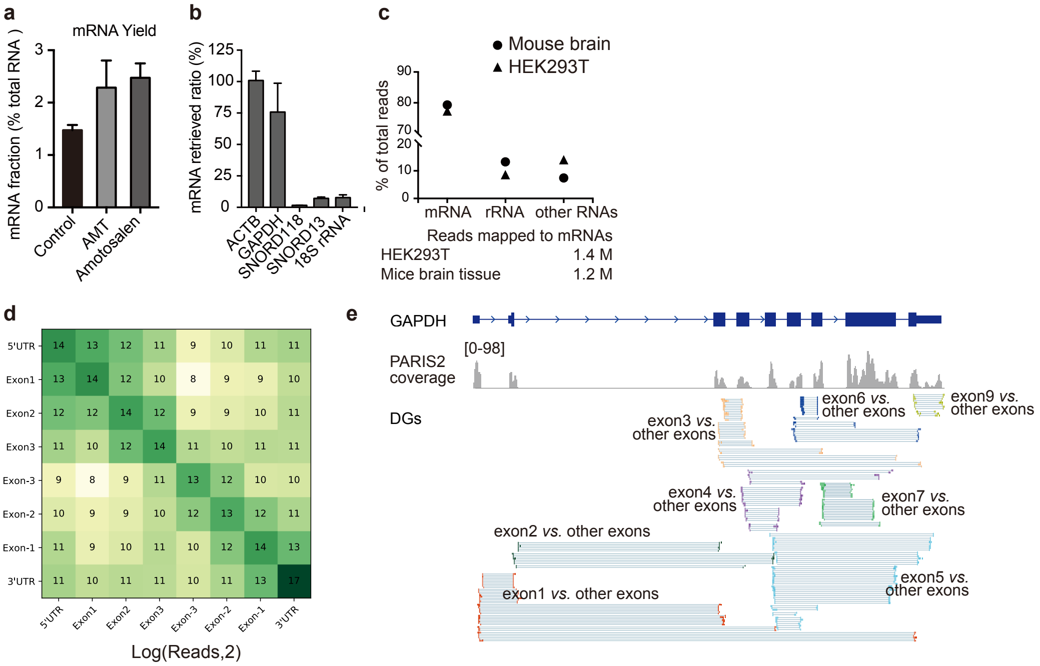

**Supplementary Figure 16. PARIS2 analysis of polyA enriched RNAs. a,** Enrichment of polyA RNA (including mRNAs and other polyA RNAs) using oligo-dT beads, from control, 0.5 mg/mL AMT and 0.5 mg/mL Amotosalen crosslinked HEK293 cells. The obvious higher yield from crosslinked samples indicate other RNAs that are covalently linked to polyA RNAs. **b,** Enrichment of mRNAs relative to noncoding RNAs based on qRT-PCR. **c**, Enrichment of mRNAs based on PARIS2 sequencing data from mouse brain and HEK293 cells. Only filtered gapped or chimeric reads are used in the calculation. **d**, Highly structured mRNA. Metagene distribution of PARIS-determined helices among each exons. **e**, PARIS2 identifies GAPDH mRNA secondary structures in HEK293T polyA enriched RNAs.

**a**

**gencode.v33.annotation.gtf, for hg38, 25 "chr"**

Pseudogenes:
IG_C_pseudogene 9
IG_J_pseudogene 3
IG_pseudogene 1
IG_V_pseudogene 188
TR_J_pseudogene 4
TR_V_pseudogene 33
polymorphic_pseudogene 55
processed_pseudogene 10173
pseudogene 18
rRNA_pseudogene 500
transcribed_processed_pseudogene 497
transcribed_unitary_pseudogene 135
transcribed_unprocessed_pseudogene 924
translated_processed_pseudogene 2
translated_unprocessed_pseudogene 1
unitary_pseudogene 97
unprocessed_pseudogene 2629

Mostly repetitive and masked

Mitochondrial RNA genes:
Mt_rRNA 2, mask and add back
Mt_tRNA 22

Multicopy noncoding genes:
scaRNA 49
snRNA 1901
snoRNA 942
misc_RNA 2212
rRNA 52 (some pseudogenes)

snRNA, mask and add back:
RNU1 (1,11 and 12): 127
RNU2: 71
RNU4 (4 and 4atac): 108
RNU5: 31
RNU6 (6 and 6atac): 1318
RNU7: 149
RNVU1: 24
RN7SK: 1
Others (alphanumeric): 81, mask

repetitive snoRNAs, do not mask:
SNORD113: 9
SNORD114: 30
SNORD115: 48
SNORD116: 30
(special analysis)
SNOR, SCA, RNU (no 113-116): 321
Others (alphanumeric): 504, mask

misc_RNA:
RNY: 56, mask, add back
7SK: 292, mask, add back
7SL: 679, mask, add back
VTRNA: 5
Others (alphanumeric): 1189, mask

Protein coding genes:
protein_coding 19957
IG_C_gene 14
IG_D_gene 37
IG_J_gene 18
IG_V_gene 144
TR_C_gene 6
TR_D_gene 4
TR_J_gene 79
TR_V_gene 106

Other RNA genes:
TEC 1060
lncRNA 16892
miRNA 1881
ribozyme 8
sRNA 5 (alphanumeric)
scRNA 1 (BCYRN1)
vaultRNA 1

Left untreated

ribozyme:
RMRP (1 real and 1 pseudo)
RPPH1 (1 real and 2 pseudo)
Others (alphanumeric): 3

**b**

**hg38_refGene.txt, from IGV, 33478 genes, masking 345**

rRNA: RNA5S, 161, RNA5-8, 16, RNA18S, 14, RNA28S, 4, RNA45S, 4.
snRNA: RNU1, 40, RNU2, 10, RNU4, 3, RNU5, 5, RNU6, 56, RNU7, 1, RNVU1, 22
Others: 7SK, 1, 7SL, 4, RNY, 4

**c**

**Dfam RNAs to mask:**

5S 1280
7SK 1276
7SLRNA 3007
HY1 462
HY3 572
HY4 174
HY5 25
LSU-rRNA_Cel 131
LSU-rRNA_Hsa 238
SSU-rRNA_Hsa 86
U1 202
U13 484
U14 8
U17 10
U2 830
U3 216
U4 169
U5 176
U6 1789
U7 241

**d**

**Human RNA, 734 from 2450 Rfam families:**

miRNAs (mir, let, lin-4): 249. Do not add back
tRNA and tRNA-Sec: 2. Do not add back
SNORA, SNORD, sno, SCAR, ACA, U3, U8: 212
ignore the others
(incomplete compared to genome)

RF00030_RNase_MRP (RMRP, 1 real, 1 pseudo in hg38)
RF00009_RNaseP_nuc (RPPH1, 1 real, 2 pseudo in hg38)
RF00024_Telomerase-vert (1 in hg38)

RF00019_Y_RNA
RF00017_Metazoa_SRP (7SL)
RF00100_7SK
RF00001_5S_rRNA
RF00003_U1
RF00004_U2
RF00015_U4
RF00020_U5
RF00026_U6
RF00548_U11
RF00007_U12
RF00618_U4atac
RF00619_U6atac
RF00066_U7

**e**

**Addback (14 "chr"):**
hssnRNA (9 together)
RNU7
hs12S
hs16S
hs5S
hs45S (18S, 5.8S, 28S)
RN7SK
RN7SL
RNY (RNY1,3,4,5)
U3
U8
U13
U14AB
U17

**f**

Before processing:
hg38.fa, 25 chr
gencode.v33.annotation.gtf, 60662 genes,

Processing:
1. Mask pseudogenes and multicopy genes from gencode, refGene and Dfam
2. Add back 14 new "genes", each as a "chromosome"

After masking and adding back single copy genes:
hg38genrefdfamadd.fa, 39 chr
gencode.v33.annotation.gtf, 60662 genes (no need to remove masked)
Plus addback genes.

**Supplementary Figure 17. Manually curated human genome reference and annotations.** We used the basic hg38 assembly, which contain 25 reference sequences, or "chromosomes", masked the multicopy genes and added back single copies. This reference is best suited for the PARIS analysis. The adjusted genome reference is used for mapping reads and IGV visualization. **a,** Classification and annotation of the Gencode v33 GTF file. Some of the snoRNAs and scaRNAs are repetitive in the genome, but we did not mask them, because there is no easy way to add back a complete set of non-redundant ones. For example, SNORD3 (10 copies), SNORD113-SNORD116 all have multiple copies. Several other snoRNAs have fewer copies. snoRNA and scaRNA paralogs can be gathered after mapping to examine interactions. **b**, List of multi-copy genes in the hg38_refGene.txt file from IGV. In the hg38_refGene.txt file, RNU1 means RNU1* (including RNU11, RNU12 etc.). **c**, Repetitive RNA genes in Dfam that needs to be masked. **d**, Classification and annotation of the human RNA genes in Rfam. **e**, The list of RNAs to add back to the masked human genome. The 9 snRNAs, U1, U2, U4, U5, U6, U11, U12, U4atac and U6atac are concatenated into one reference, separated by 100nt "N"s. The entire 45S unit is added as one reference. Note. ITS and ETS regions in rRNAs are not masked properly, so reads mapped to these regions should be treated properly. The 4 RNY genes are concatenated with 100N spacers. **f,** Pipeline to mask hg38 and add back single copy genes, and summary of the input and output files. Scripts used: maskgencode.py, maskrefgene.py, maskdfam.py.

**a**

**gencode.vM24.annotation.gtf for mm10, 22 “chr”**

Pseudogenes:
IG_C_pseudogene 1
IG_D_pseudogene 3
IG_pseudogene 2
IG_V_pseudogene 158
TR_J_pseudogene 10
TR_V_pseudogene 34
polymorphic_pseudogene 88
processed_pseudogene 10002
pseudogene 60
transcribed_processed_pseudogene 300
transcribed_unitary_pseudogene 25
transcribed_unprocessed_pseudogene 272
translated_unprocessed_pseudogene 1
unitary_pseudogene 58
unprocessed_pseudogene 2716

Mostly repetitive and masked

Mitochondrial RNA genes:
Mt_rRNA 2, mask and add back
Mt_tRNA 22

Multicopy noncoding genes:
scaRNA 51 (no distinct names)
snRNA 1383 (no distinct names) mask and add back
snoRNA 1507 (no distinct names)
misc_RNA 562 (no distinct names)
rRNA 354 (no distinct names) mask and add back

Protein coding genes:
protein_coding 21856
IG_C_gene 13
IG_D_gene 19
IG_J_gene 14
IG_LV_gene 4
IG_V_gene 218
TR_C_gene 8
TR_D_gene 4
TR_J_gene 70
TR_V_gene 144

Other RNA genes:
TEC 3238
lncRNA 9959
miRNA 2202
ribozyme 22
sRNA 2 (alphanumeric)
scRNA 1 (Bc1-ps1)

Left untreated

ribozyme:
Rmrp (1 real and ?)
RPPH1 (1 real, 3 Rprl and ?)
Others (alphanumeric): 17

**b**

**mm10_refGene.txt, from IGV, 36868 genes**

Unlike hg38, many RNA genes are not annotated in mm10_refGene.txt. The 5.8S rRNA locations were extracted from mouse genome+transcripts using BLAST. Among three full length matches, one perfect match is on a 45S sequence two imperfect matches are on assembled chromosomes: chr18:73533406-73533537 (88% identity), chr6:94826786-94826922 (77% identity).

**c**

**Dfam RNAs to mask:**

LSU-rRNA_Cel 61
LSU-rRNA_Hsa 182
SSU-rRNA_Cel 1
SSU-rRNA_Hsa 39
U1 320
U13 41
U14 4
U17 21
U2 819
U3 99
U4 97
U5 79
U6 1472
U7 53
U8 6
HY1 43
HY3 14
HY4 5
HY5 1
4.5SRNA 2230
5S 1159
7SK 719
7SLRNA 711
BC1_Mm 31912

**d**

**Mouse RNA, 539 from 2450 Rfam families:**

miRNAs (mir, let-7, lin-4): 179. Do not add back
tRNA and tRNA-Sec: 2. Do not add back
SNORA, SNORD, sno, SCAR, ACA, U3, U8: 191
ignore the others (IRES, RNA motifs, etc.)

incomplete compared to genome

RF00030_RNase_MRP (RMRP)
RF00009_RNaseP_nuc (RPPH1)
RF00024_Telomerase-vert (TERC)

RF00019_Y_RNA
RF00017_Metazoa_SRP (7SL)
RF00100_7SK
RF00001_5S_rRNA
RF00003_U1
RF00004_U2
RF00015_U4
RF00020_U5
RF00026_U6
RF00548_U11
RF00007_U12
RF00618_U4atac
RF00619_U6atac
RF00066_U7

**e**

**Addback 16 “chr”:**

mmsnRNA (9 together)
RNU7
mm12S
mm16S
mm5S
mm45S (18S, 5.8S, 28S)
RN7SK
RN7SL
RNY (RNY1,3,4,5)
U3
U8
U13
U14
U17
mm4.5S
mmBC1

**f**

Before processing:
mm10.fa, 22 chr
gencode.vM24.annotation.gtf, 55385 genes

Processing:
1. Mask pseudogenes and multicopy genes from gencode, refGene and Dfam
2. Add back 16 new “genes”, each as a “chromosome”

After masking and adding back single copy genes:
mm10genrefdfamadd.fa, 38 chr
gencode.vM24.annotation.masked.gtf, 55385 genes (no need to remove masked)
Plus addback genes

**Supplementary Figure 18. Manually curated mouse genome reference and annotations.** We used the basic mm10 assembly, which contain 22 reference sequences, or “chromosomes”, masked the multicopy genes and added back single copies. This reference is best suited for the PARIS analysis. The adjusted genome reference is used for mapping reads and IGV visualization. **a**, Classification and annotation of the Gencode vM24 GTF file. Some of the snoRNAs and scaRNAs are repetitive in the genome, but we did not mask them, because there is no easy way to add back a complete set of non-redundant ones. For example, SNORD3, SNORD113-SNORD116 all have multiple copies. Several other snoRNAs have fewer copies. snoRNA and scaRNA paralogs can be gathered after mapping to examine interactions. **b**, To maintain consistency with the hg38 genome curation, we also examined the mm10_refGene.txt from IGV, but this annotation missed most of the multicopy RNA genes. **c**, Multicopy RNA genes from Dfam that needs to be masked. **d**, Classification and annotation of the mouse RNA genes in Rfam. **e**, The list of RNAs to add back to the masked mouse genome. The 9 snRNAs, U1, U2, U4, U5, U6, U11, U12, U4atac and U6atac are concatenated into one reference, separated by 100nt “N”s. The entire 45S unit is added as one reference. Note. ITS and ETS regions in rRNAs are not necessarily masked properly, so reads mapped to these regions should be treated properly. **f,** Pipeline to mask hg38pri and add back single copy genes. The 2 RNY genes, they are not all masked in the Gencode list. mm4.5S and mmBC1 are two noncoding RNAs that are not present in the human genome.

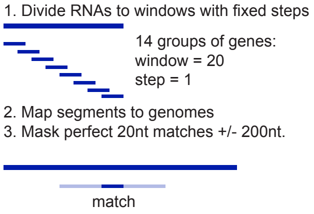

**Supplementary Figure 19. Masking multi-copy genes in the genome.** This is a general strategy for analyzing targets of repetitive ncRNAs across the genome. Fragments of multi-copy genes are extracted in windows (e.g. window size of 20 and step=1), and maped to the genome using STAR. Then perfect matches in the genome are extended to both sides by a fixed length, e.g. 200nt. The match+extension regions are masked, and then single copies of these genes are added back before mapping. This approach may lead to false positive masking: some genes may be unrelated to snRNAs and still masked. However, the false masking should be rare, especially for exons in protein-coding and lncRNAs, and the results should be very reliable.

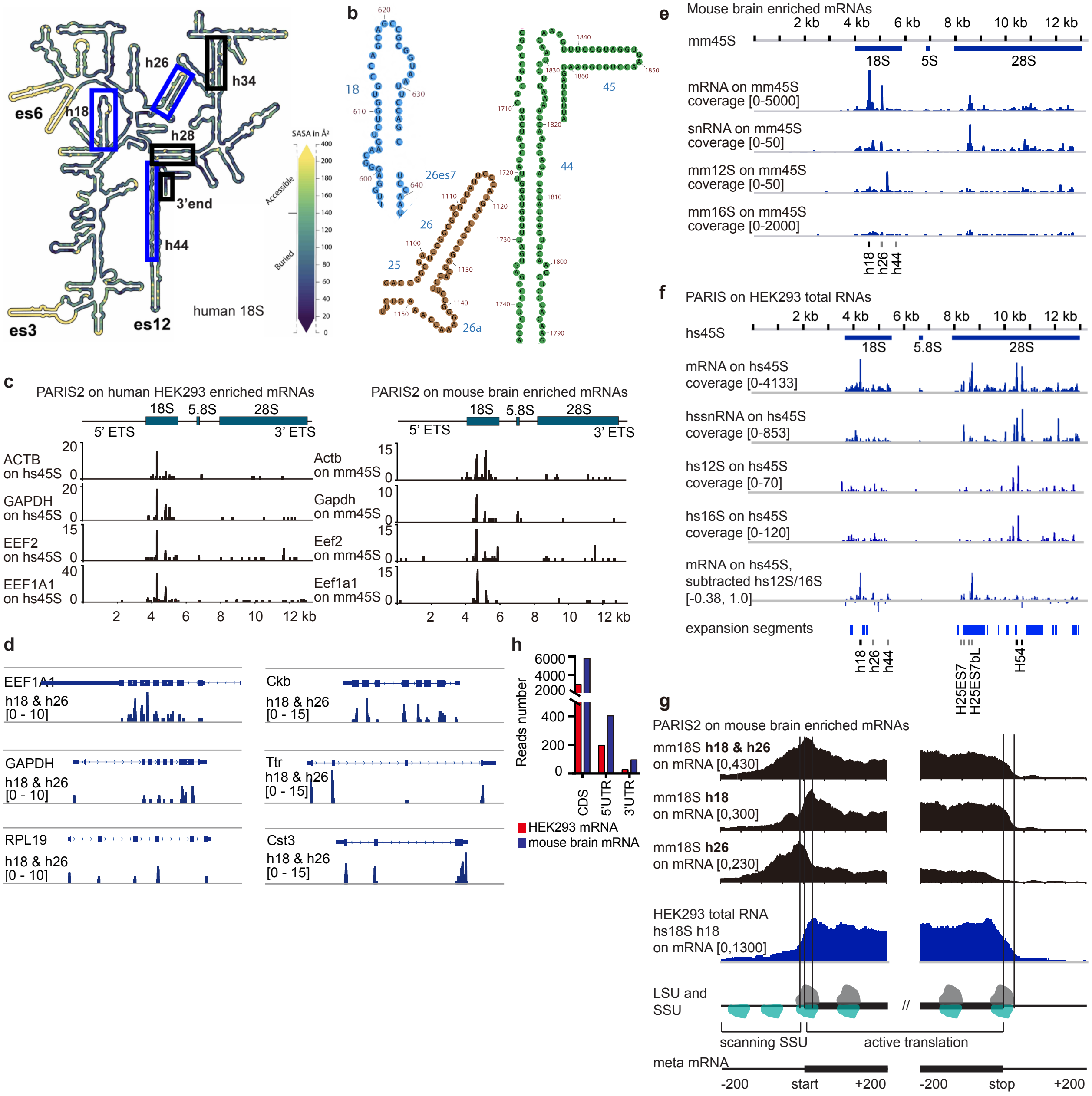

**Supplementary Figure 20. PARIS enables ribosome SSU profiling. a,** Locations of the mRNA-interacting regions in the 18S ribosomal RNA secondary structure. es3, es6 and es12 are three prominent expansion sequences in 18S. Solvent accessible areas are marked based on the ribosome gallery, from GATech. The blue boxes indicate the three mRNA-interacting 18S regions detected in PARIS. The 3 black boxes highlight additional regions in 18S that can be crosslinked to mRNAs based on Pisarev et al. 2008. **b**, The three18S regions highlighted in blue in panel d, including h18, h26, h44-h45 and the 3' end of the transcript. For h18, only the left arm is crosslinked to mRNAs. For the h26-h26es7 helix, both arms are crosslinked to mRNAs. h44 left arm is croslinked to mRNAs. **c**, Examples of mRNA-rRNA interactions in human HEK293 and mouse brain enriched mRNA PARIS2 data. **d**, Examples of mRNAs interact with h18 & h26 domains on 18S rRNA. Left side, HEK293T cells enriched mRNAs; Right side, mouse brain tissues enriched mRNAs. **e**, mRNA binding sites on the mouse 45S rRNA. snRNA, 12S and 16S serve as controls for the specificity. **f**, mRNA binding sites on human 45S rRNA, based on data from PARIS on total RNA in HEK293T cells (Lu et al. 2016 Cell). The H25ES7 and H25ES7bL peaks that remain after subtracting hs12S and hs16S binding sites are likely to be real crosslinking events. **g**, SSU h18 and h26 binding sites on mouse and mouse mRNAs. **h**, Comparison of 18S h18 & h26 associated reads number on mRNAs in CDS, 5'UTR and 3'UTR regions. 50 nt of 5'UTR and 3'UTR next to CDS are excluded to avoid the extended tails from reads mapped to the CDS. Read numbers are as follows in the order of CDS, 5'UTR and 3'UTR. HEK293: 2958, 196 and 26, mouse brain tissue: 5799, 403 and 95.

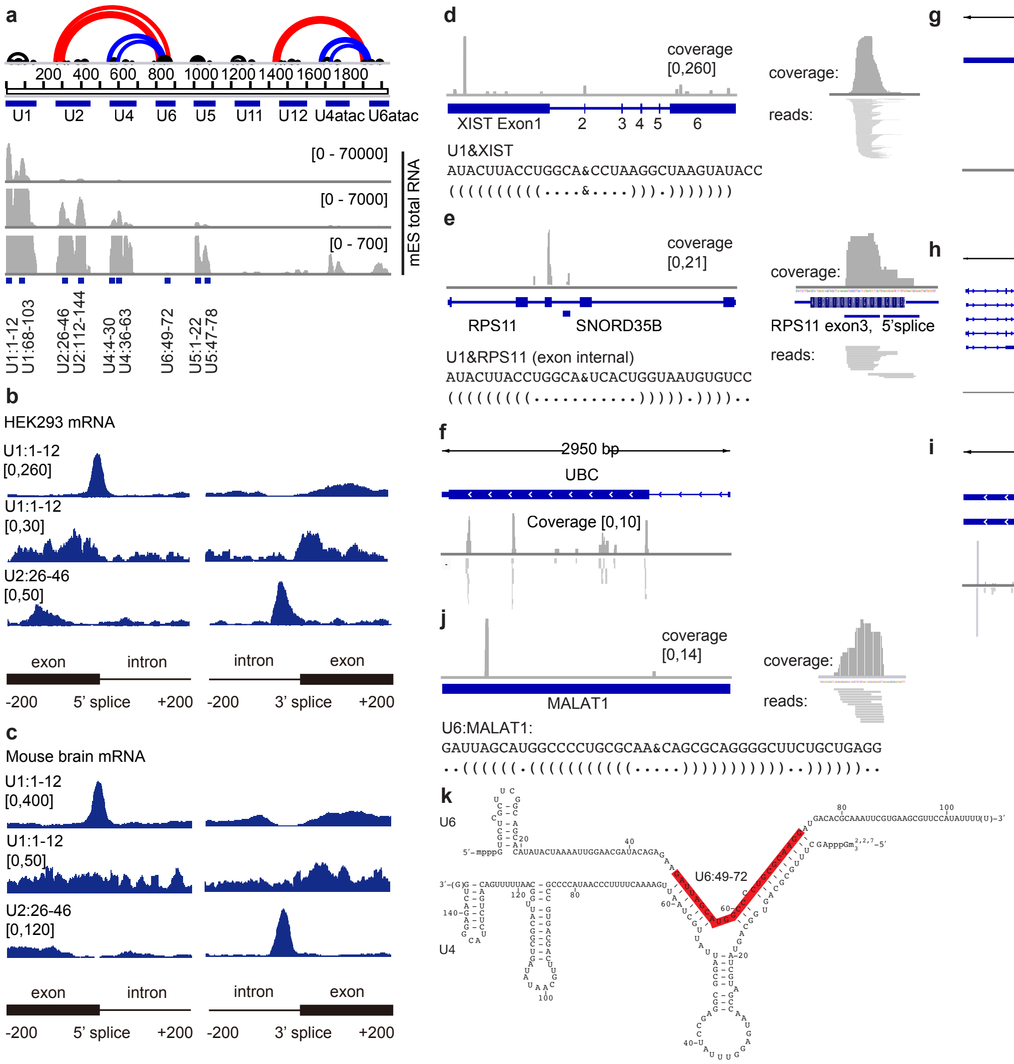

**Supplementary Figure 21. snRNP profiling using PARIS. a**, mouse snRNAs that interact with mRNAs. Chimeric reads in mouse ES cell total RNA PARIS data that connect mRNAs to snRNAs were plotted. Major peaks are highlighted. **b-c**, HEK293T polyA enriched mRNAs and mouse brain enriched mRNAs were plotted, showing the major binding sites of snRNAs on a meta-mRNA. **d-i**, Examples of snRNA U1 binding sites on exonic regions. In panel (e), there is clear binding both at the splice site and far from the splice site in RPS11 mRNA exon3. **j**, U6 binding site on MALAT1 mRNA. **k**, Location of the U6 sequence that bind MALAT1, plotted on the U4:U6 heterodimer.

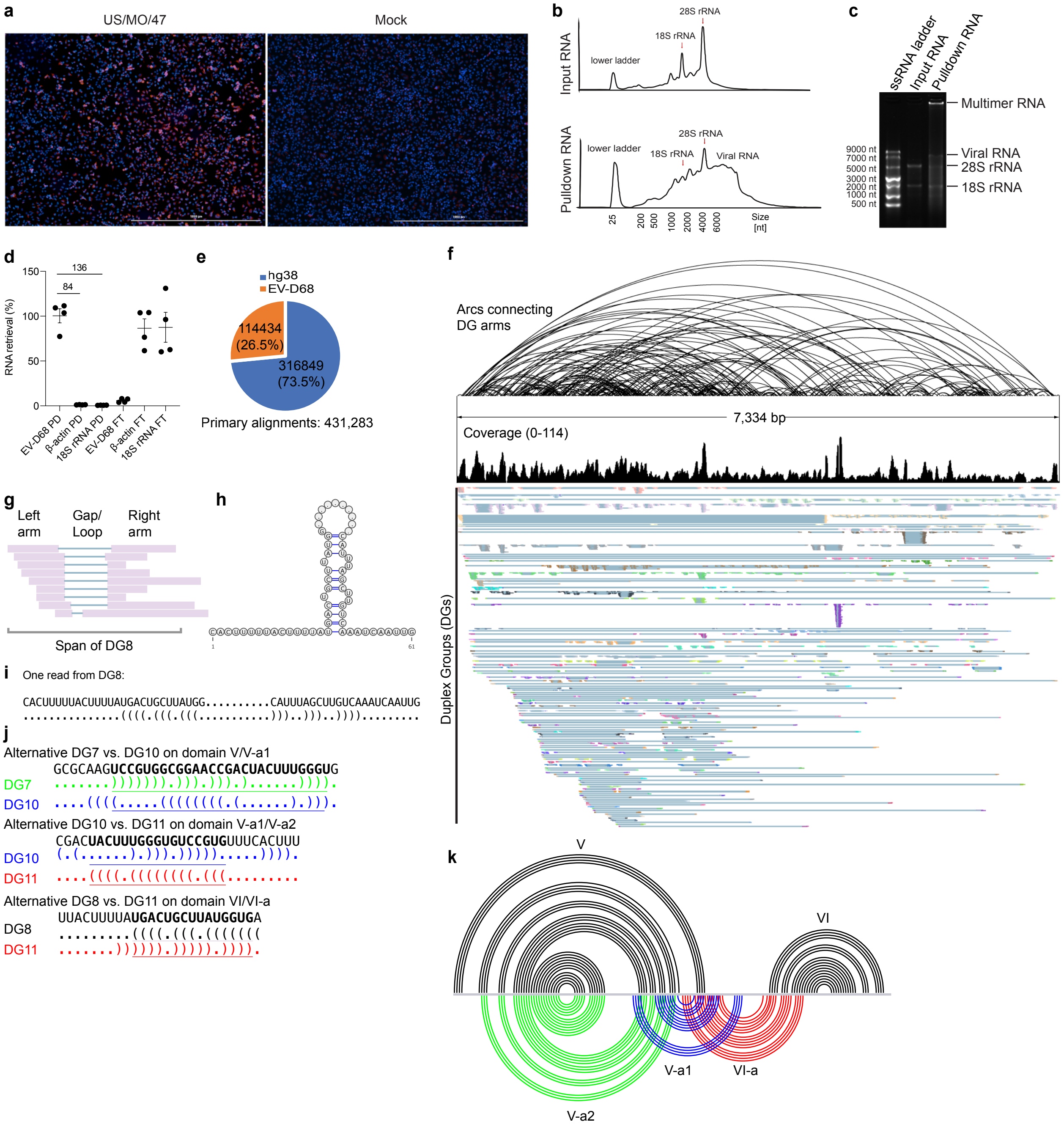

**Supplementary figure 22. Enrichment of EV-D68 genome RNA for PARIS2 analysis and the global architecture. a**. EV-D68 US47 infection of HeLa cells detected by immunostaining assay. Cells were fixed at 18 hours post infection of US/MO/47 strain at an MOI of 1.0 and stained with polyclonal antibody against VP1 of EV-D68 (red) and counterstained with DAPI (blue) for detection of nuclei. **b**.Input and EV-D68 RNA pulldown profiles based on TapeStation analysis (retraced and aligned based on the agarose gel electrophoresis). **c**. Agarose gel RNA profiles of EV-D68 pulldown or input RNA from cells infected with EV-D68. **d**. Enrichment of viral genome RNA and depletion of host RNAs based on qRT-PCR. PD: pulldown, FT: flow-through. Experiments were repeated independently 4 times. Viral RNA primers and two sets of control primers for beta-actin and 18S rRNA, respectively, were used to amplify EV-D68 PD and FT samples. The enrichment folds of viral RNA versus both controls are labeled above the black lines. **e**, Pie chart showing highly efficient retrieval of viral RNA based on mappable reads to the human genome vs. the EV-D68 genome. **f**. The full coverage of gapped reads cover the whole genome RNA of EV-D68, the duplex groups and the structure model. The uneven coverage is due to variations in the strength of local structures. **g-i**, An example structure from Domain VI based on PARIS-derived DG8, showing the gapped reads (g), secondary structure model (h) and one example read (i). **j**. Three alternative structures are displayed in dot-bracket format and color matched to Fig. 6. **k**, Zoom in view of domain V, VI and their alternative conformations. V-a2 is part of V. The alternative structures of domain V (V-a1 and V-a2) and domain VI (VI-a) are highlighted in blue, green, and red, respectively.

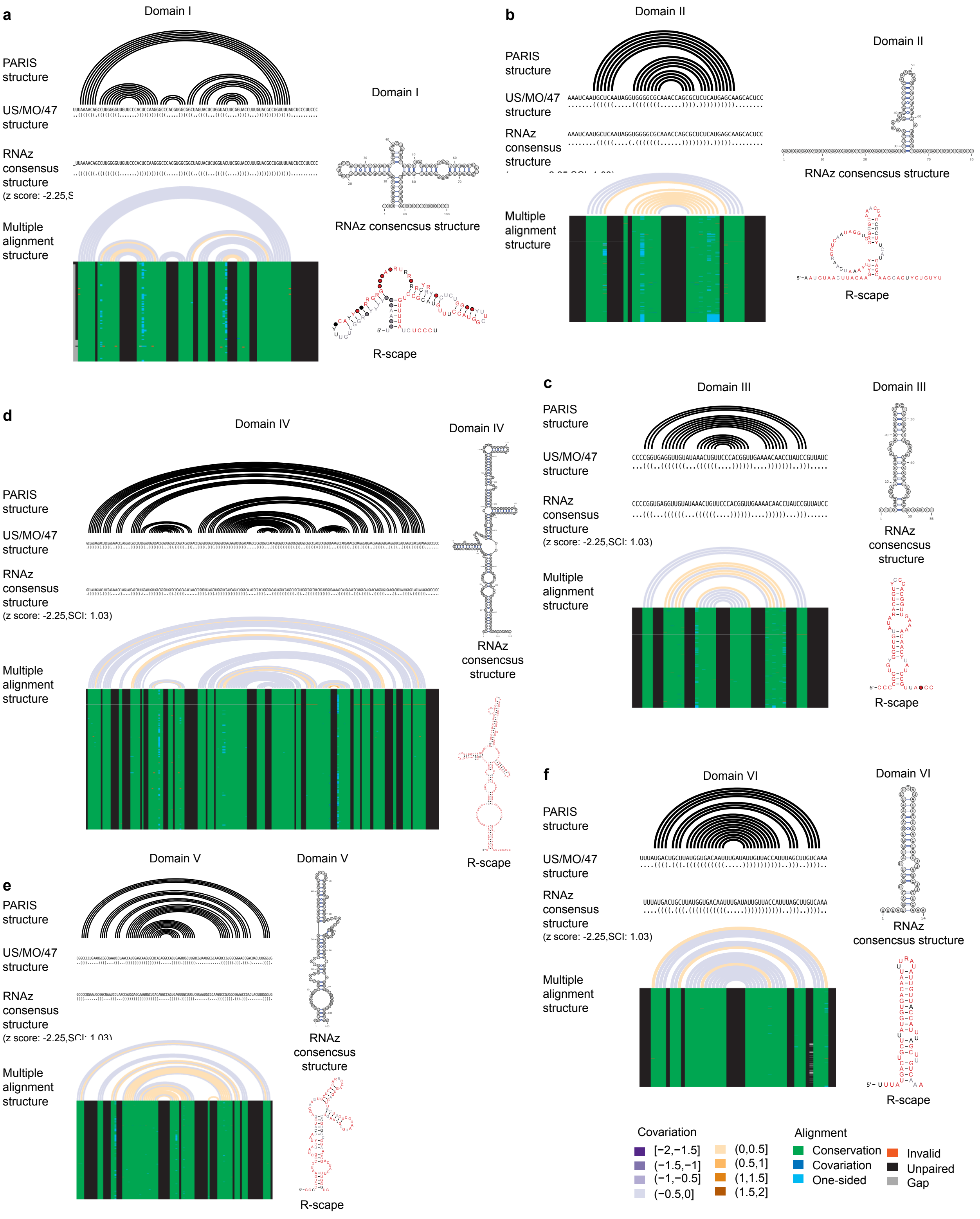

**Supplementary Figure 23. Conservation analysis of EV-D68 alternative conformations. a-f**, RNAz and R-scape analysis of the conservation of Domains 1-VI in all EVD strains with complete genomes.

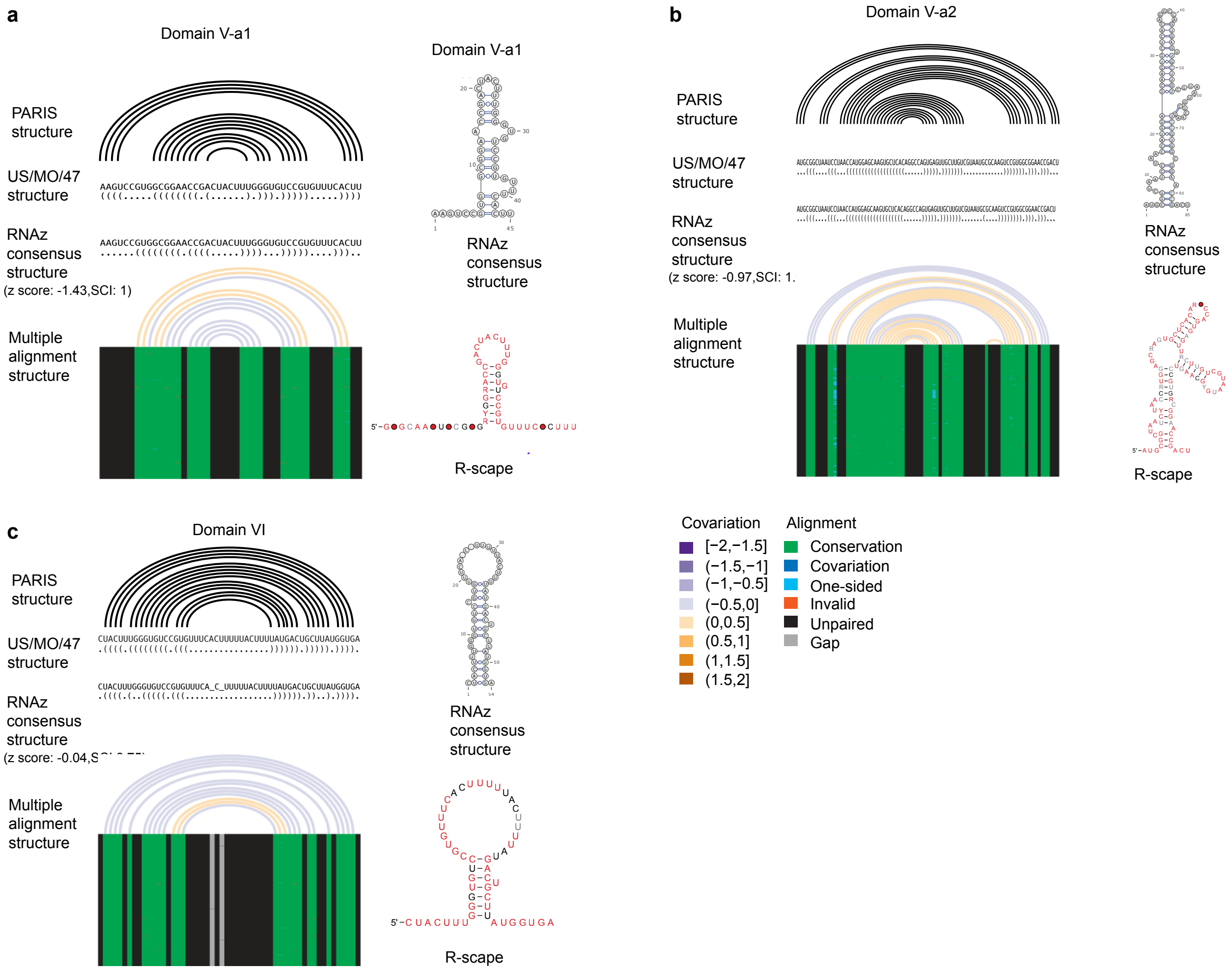

**Supplementary Figure 24. Conservation analysis of EV-D68 alternative conformations. a-c**, The new domains in the identified IRES alternative structures supported by conservation analysis using MUSCLE multiple sequence alignment (492 unique EV-D68 complete genomes) and RNAz. The consensus structures above arc diagrams are predicted from RNAalifold and arc diagrams are displayed by R-chie. On the right is consensus structures supported by RNAz and R-scape. By default only positions in the alignment with more than 50% occupancy are depicted.

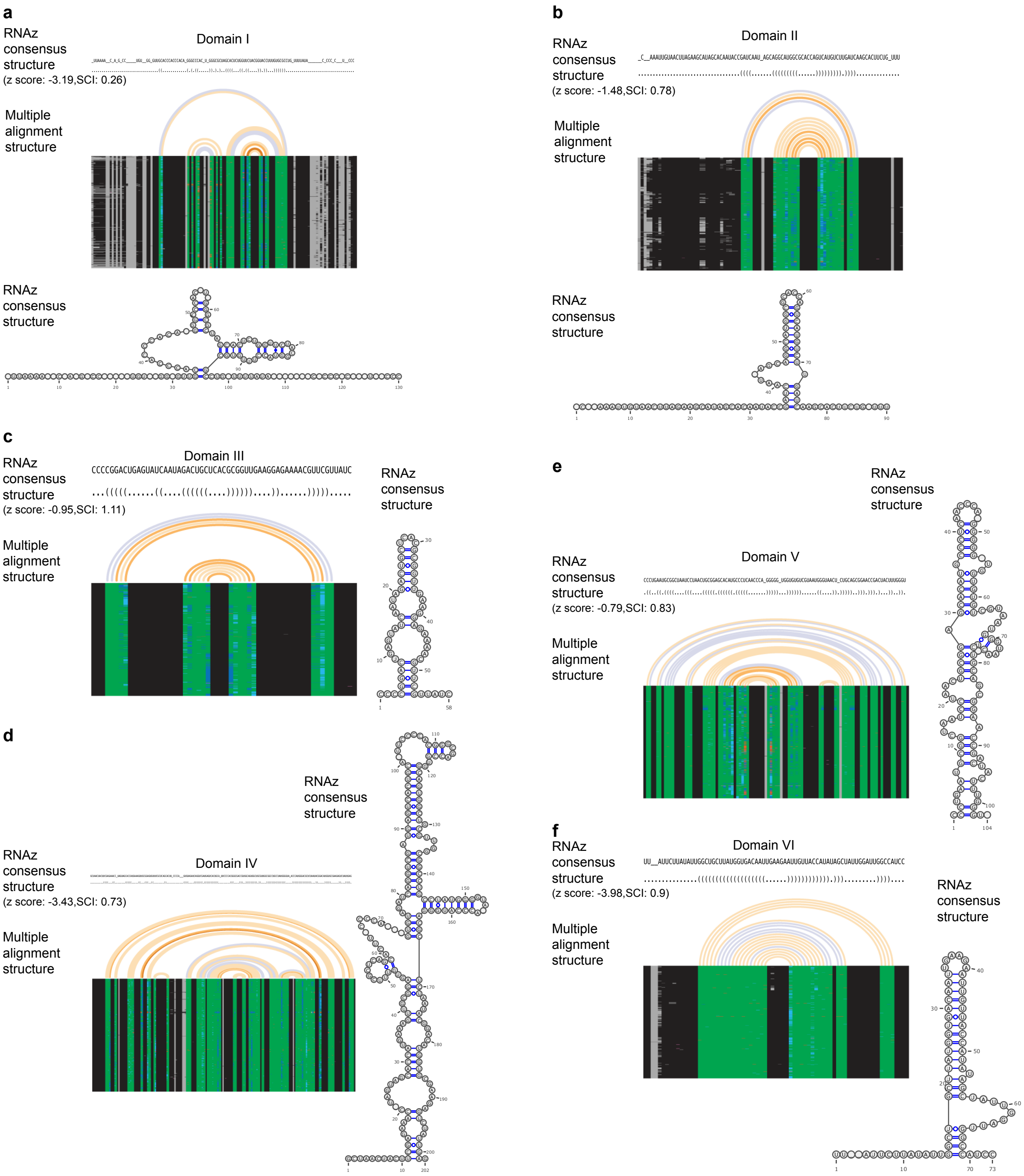

**Supplementary Figure 25. Conservation analysis of enterovirus A using RNAz guided by US/MO/47 structures identified by PARIS2. a-f**. Predicted structures of domain I-VI of typical type I IRES from 499 sequences of enterovirus A species.

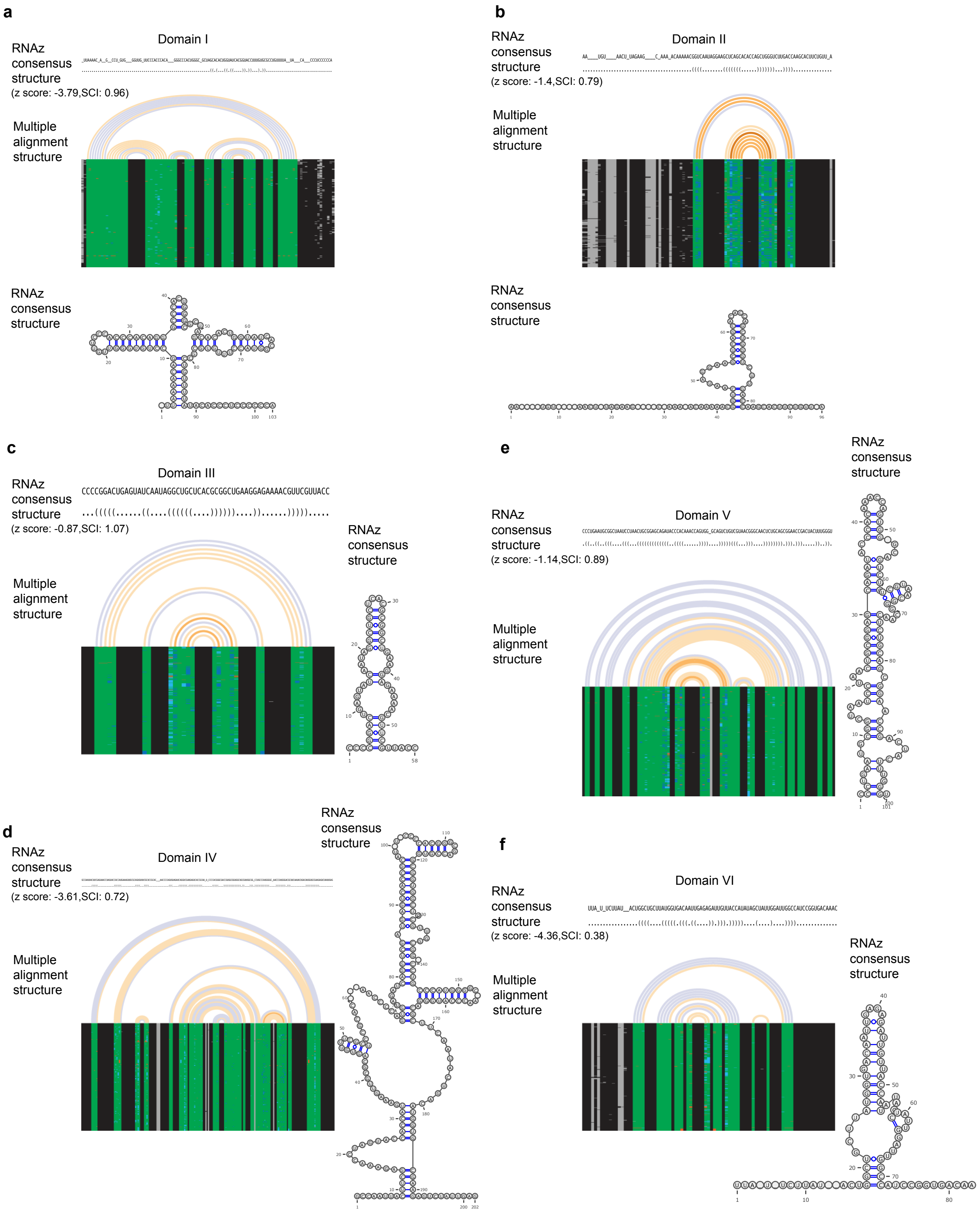

**Supplementary Figure 26. Conservation analysis of enterovirus B using RNAz guided by US/MO/47 structures identified by PARIS2. a-f**. Predicted structures of domain I-VI of typical type I IRES from 379 sequences of enterovirus B species.

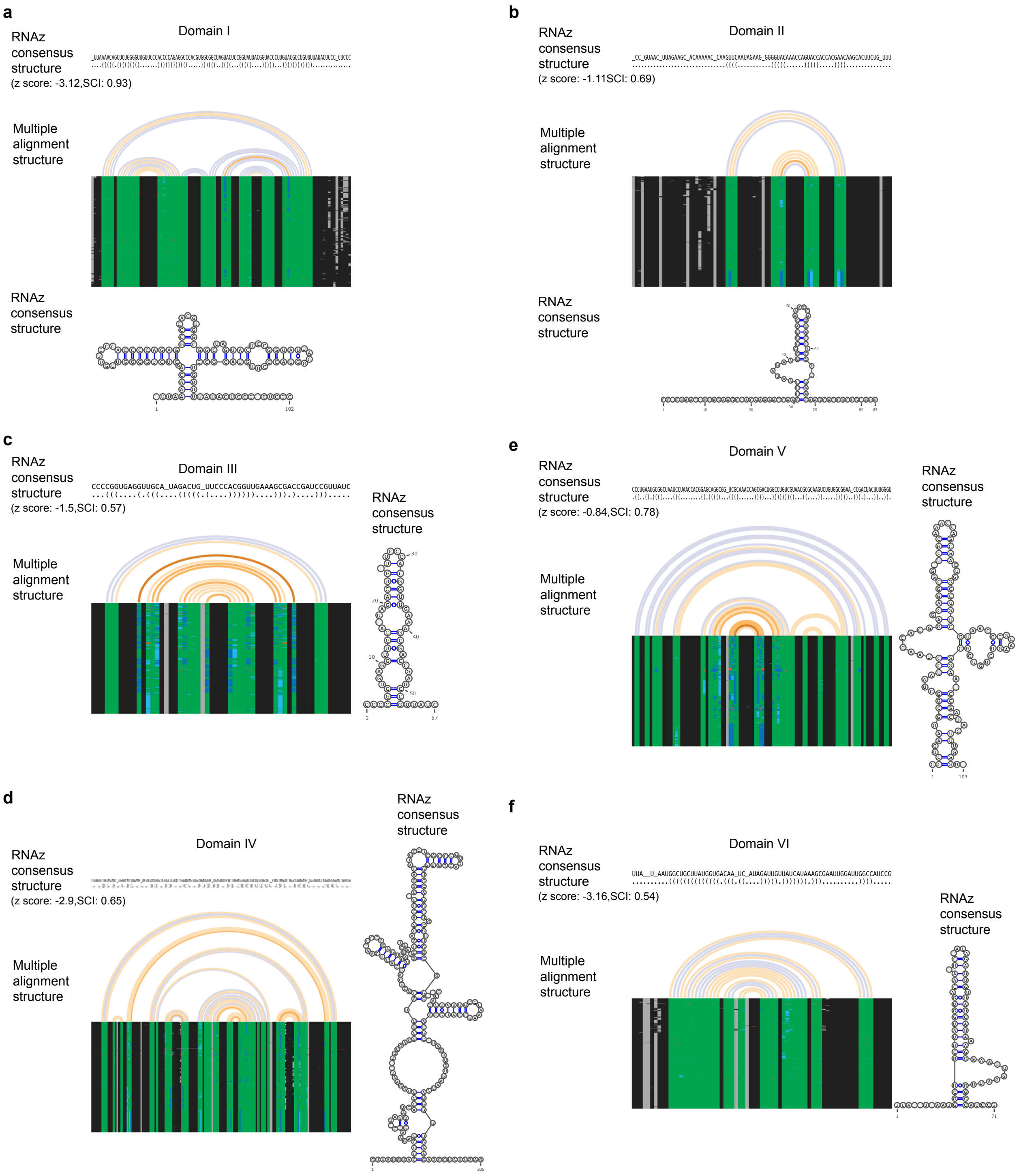

**Supplementary Figure 27. Conservation analysis of enterovirus C using RNAz guided by US/MO/47 structures identified by PARIS2. a-f**. Predicted structures of domain I-VI of typical type I IRES from 499 sequences of enterovirus C species.

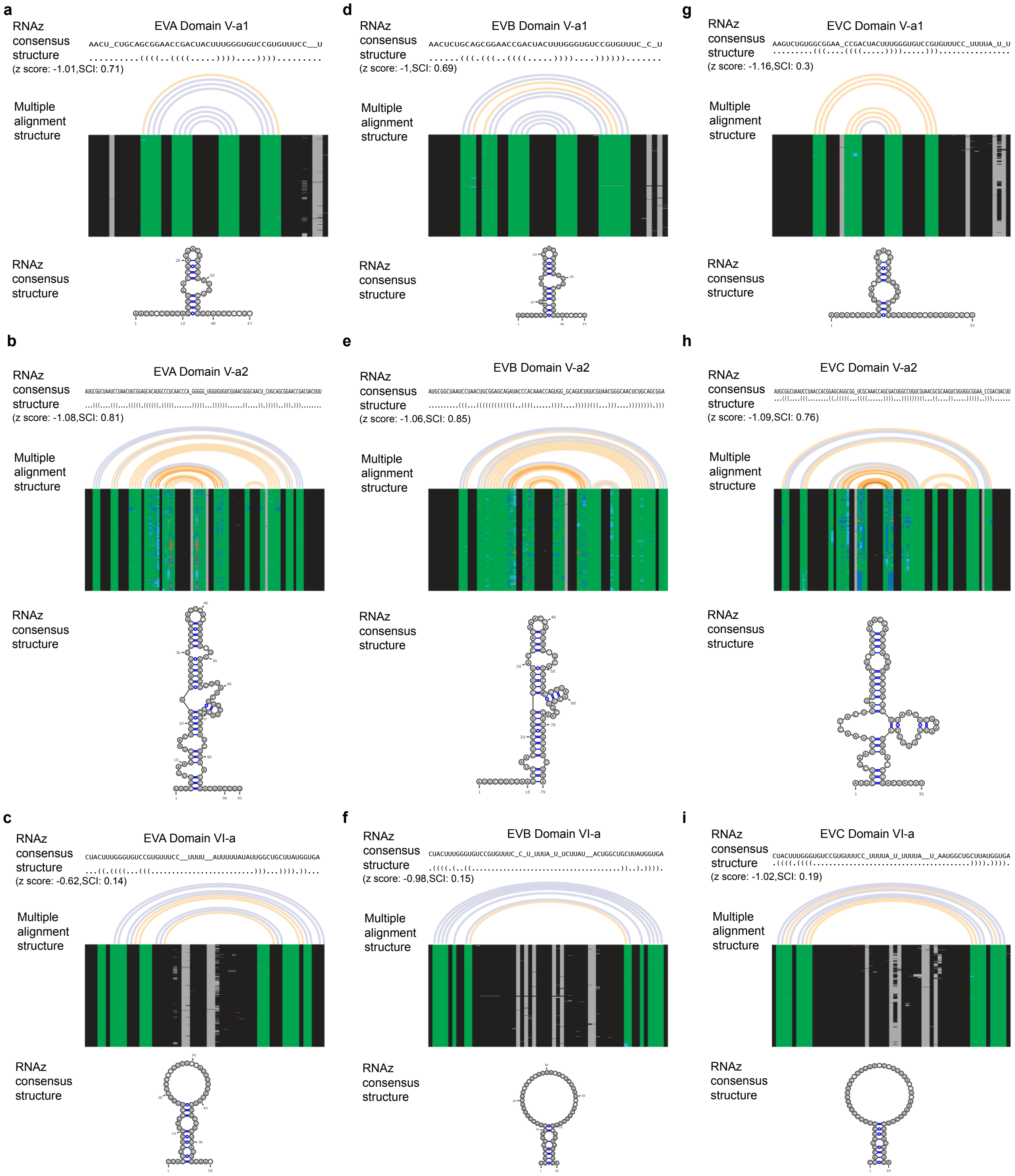

**Supplementary Figure 28. Conservation analysis of IRES alternative structures in enterovirus species A, B, and C using RNAz guided by US/MO/47 structures identified by PARIS2. a-c**. Alternative structures domain V-a1, V-a2, and VI-a of enterovirus A. **d-f**. Alternative structures domain V-a1, V-a2, and VI-a of enterovirus B. **g-i**. Alternative structures domain V-a1, V-a2, and VI-a of enterovirus C.

Fig. 4f

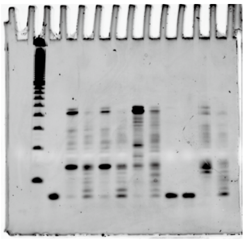

Fig. 4g

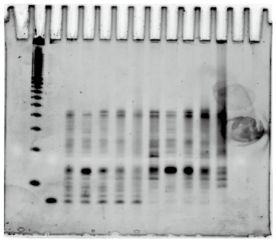

Supplementary Fig. 4d

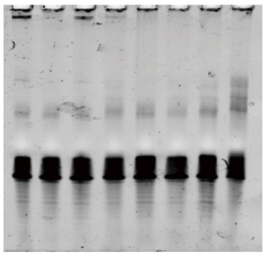

Supplementary Figure 10i

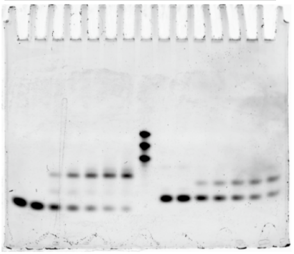

Supplementary Figure 10j

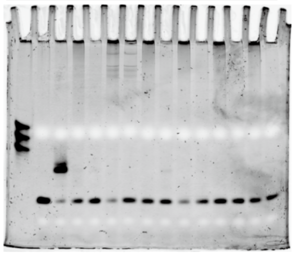

Supplementary Figure 10k

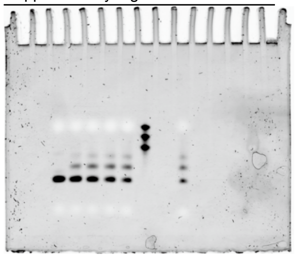

Supplementary Figure 10l

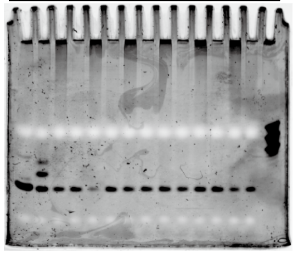

Supplementary Figure 12b

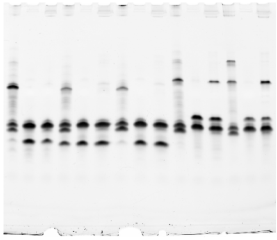

Supplementary Figure 12c

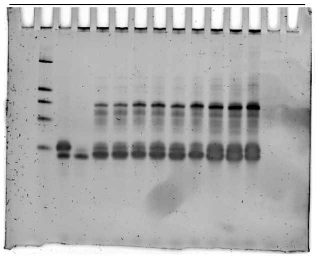

Supplementary Figure 12d

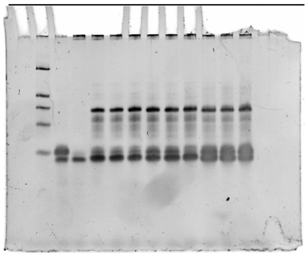

Supplementary Figure 12e

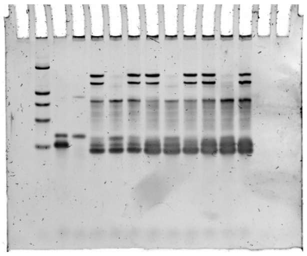

Supplementary Fig. 15f

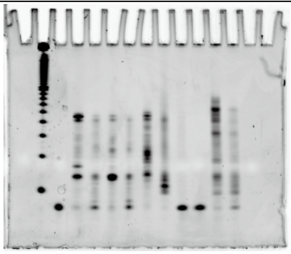

Supplementary Fig. 15g

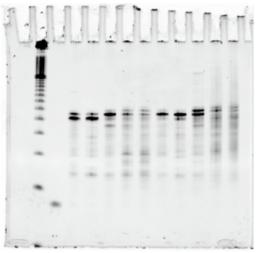

**Supplementary Figure 29. Original gel pictures**

| Sample preparation | Comparison | Average Ratio | Related figures |
|---|---|---|---|
| Amotosalen crosslinking<br>• 0.5 mg/ml amotosalen<br>• 2.0 mg/ml amotosalen<br>• 5.0 mg/ml amotosalen | 0.5 AMT | 3.42–6.95<br>• 3.42<br>• 5.13<br>• 7.95 | Fig. 2 |
| TNA method<br>• 0.5 mg/ml amotosalen<br>• 5.0 mg/ml amotosalen | TRIzol | 2.13–6.48<br>• 2.13<br>• 6.48 | Fig. 3<br>Supplementary Fig. 3 and 6 |
| DD2D gel purification | ND2D | 1.54 | Supplementary Fig. 8 |
| Total improvement ratio<br>• 0.5 mg/ml amotosalen<br>• 5.0 mg/ml amotosalen | PATIS1 | 11.21–69.35<br>• 11.21<br>• 69.35 | |

| Library preparation (after 2D gel) | Comparison | Average Ratio | Related figures |
|---|---|---|---|
| Adapter ligation<br>• SLRNA2<br>• SLRNA8 | Standard | 2.45–3.91<br>• 2.45<br>• 3.91 | Supplementary Fig. 9 |
| UVC protection (AO)<br>• based on GAPDH (70 bp)<br>• based on ACTB (184 bp) | no AO | 6.51 - 9.90<br>• 6.51<br>• 9.90 | Fig. 4<br>Supplementary Fig. 11 |
| PUVA bypass (SSIV) | SSIII | 3.32 | Fig. 4 |
| PUVA bypass (SSIV with $Mn^{2+}$ buffer)<br>• based on SONRD118 (96 bp)<br>• based on SNORD13 (63 bp)<br>• based on ACTB (184 bp) | $Mg^{2+}$ buffer | 2.48 - 23<br>• 2.48<br>• 2.62<br>• 23.03 | Fig. 4<br>Supplementary Fig. 15 |
| Total library yield | PARIS1 | 76 | |

**Supplementary Table 1. Summary of PARIS2 improvements.** The sample and library preparation improvements are listed separately. Sample preparation starts from crosslinking of cells to the 2D gel isolation of croslinked RNA. The library preparation starts from the 2D gel purified crosslinked RNA to the final DNA library ready for sequencing. See Supplementary Table 2 for detailed numbers of the improvements for every step. PUVA, psoralen plus UVA; AO, arcidine orange; SSIII, Superscript III; SSIV, Superscript IV. The TNA vs. TRIzol comparison is based on the improvement over all retrieved RNA from aqueous phase in TRIzol. Therefore the improvement of recovery for larger RNAs is even higher. The total library yield improvement of 76 fold was based on experimental results of library yield from the same amount of starting crosslinked RNA after 2D gel purification (not the same total RNA, see Supplementary Table 2). This number is close to the lower bound of the theoretical multiplied improvements for the library preparation steps (adapter ligation, UVC protection, PUVA bypass (SSIV) and PUVA bypass (SSIV with $Mn^{2+}$ buffer)), which is in the range of [131, 2956]. This is because the several steps could be bottlenecks at the same time and improving one step may not be sufficient to lift the yield for the entire pipeline. The prevention of UVC-induced damage and bypass of PUVA-induced damage is positively correlated to RNA length. Longer amplicons contain more damage sites, and therefore the

| **Step1. Amotosalen crosslinking** | | | | |
|---|---|---|---|---|
| Crosslinked RNA fraction | 0.5 AMT | 0.5 Amoto | 2.0 Amoto | 5.0 Amoto |
| Rep1 | 0.92% | 1.83% | 3.43% | 4.65% |
| Rep2 | 0.63% | 2.71% | | |
| Rep3 | 0.65% | 1.72% | | |
| Rep4 | 0.45% | 2.90% | | |
| Rep5 | 0.70% | | | |
| Average fraction | 0.67% | 2.29% | 3.43% | 4.65% |
| Ratio (to 0.5 AMT) | 1.00 | 3.42 | 5.13 | 6.95 |

| **Step2. TNA method** | | | | | | | | |
|---|---|---|---|---|---|---|---|---|
| RNA yield (μg) form 2 million cells | TRIzol method | | | | TNA method | | | |
| | Ctrl | 0.5 AMT | 0.5 Amoto | 5.0 Amoto | Ctrl | 0.5 AMT | 0.5 Amoto | 5.0 Amoto |
| Rep1 | 20.06 | 6.86 | 6.00 | 3.18 | 19.6 | 13.34 | 11.98 | 18.64 |
| Rep2 | 18.46 | 6.26 | 5.96 | 2.96 | 18.56 | 14.36 | 14.82 | 21.3 |
| Rep3 | NA | NA | NA | NA | 16.82 | 11.54 | 11.44 | 19.78 |
| Average yield | 19.26 | 6.56 | 5.98 | 3.07 | 18.33 | 13.08 | 12.75 | 19.91 |
| Riato (to TRIzol) | | | | | 0.95 | 1.99 | 2.13 | 6.48 |

| **Step3. DD2D** | | |
|---|---|---|
| Crosslinked RNA fraction | ND2D | DD2D |
| Rep1 | 0.47% | 0.70% |
| Rep2 | 0.35% | 0.63% |
| Rep3 | 0.46% | 0.65% |
| Average fraction | 0.43% | 0.66% |
| Ratio (to ND2D) | | 1.54 |
| Note: ND2D data is from Figs. 1e of Lu. *et al.* Cell 2016. | | |

| **Step4. Adapter ligation** | | | |
|---|---|---|---|
| | tested oligos | Control | New condition |
| Adj. Vol. (Int) | SLRNA2 | 40479770 | 158230455 |
| Adj. Vol. (Int) | SLRNA8 | 26086600 | 64037635 |
| Ratio | SLRNA2 | | 3.91 |
| Ratio | SLRNA8 | | 2.45 |
| Adj. Vol. (Int), gel quantification data by Image Lab 6.0 Software. | | | |

| **Step5. UVC protection (AO)** | | | | |
|---|---|---|---|---|
| Fold change of cDNA yield (normalized to control) | GAPDH | | ACTB | |
| | No protection | With protection | No protection | With protection |
| Rep1 | 0.09 | 0.63 | 0.04 | 0.91 |
| Rep2 | 0.08 | 0.66 | 0.04 | 0.99 |
| Rep3 | 0.15 | 1.41 | 0.04 | 2.28 |
| Rep4 | 0.12 | 1.09 | 0.35 | 3.48 |
| Rep5 | 0.09 | 1.09 | 0.14 | 1.07 |
| Rep6 | 0.28 | 1.08 | 0.54 | 0.87 |
| Rep7 | 0.27 | 1.52 | 0.29 | 2.07 |
| Rep8 | 0.30 | 1.37 | 0.13 | 2.14 |
| Rep9 | 0.18 | NA | 0.14 | 2.00 |
| Rep10 | 0.14 | NA | 0.13 | 2.37 |
| Average cDNA yield | 0.17 | 1.11 | 0.18 | 1.82 |
| Ratio (to no protection) | | 6.51 | | 9.90 |

| **Step6. PUVA bypass (SSIV)** | | |
|---|---|---|
| Fold change of cDNA yield (normalized to SSIII) | SSIII | SSIV |
| Rep1 | 1.00 | 2.29 |
| Rep2 | 1.00 | 3.01 |
| Average ratio (to SSIII) | | 2.65 |

| **Step7. PUVA bypass (SSIV with $Mn^{2+}$ buffer)** | | | |
|---|---|---|---|
| Fold change of cDNA yield (normalized to $Mg^{2+}$ buffer) | ACTB | SNORD118 | SNORD13 |
| Rep1 | 19.33 | 2.57 | 2.77 |
| Rep2 | 21.48 | 2.46 | 2.47 |
| Rep3 | 19.65 | 2.39 | |
| Rep4 | 31.62 | | |
| Average ratio (to $Mg^{2+}$) | 23.02 | 2.48 | 2.62 |

| **Total library yield** | | |
|---|---|---|
| Library yield (nmol) | PARIS1 | PARIS2 |
| Rep1 | 0.38 | 60.19 |
| Rep2 | 0.71 | 51.86 |
| Rep3 | 1.59 | 93.17 |
| Rep4 | 0.49 | 42.86 |
| Rep5 | NA | 58.85 |
| Rep6 | NA | 53.27 |
| Average yield | 0.79 | 60.03 |
| Ratio (to PARIS1) | | 75.75 |
| Note: Library products yield from 50 ng of crosslinked RNA. | | |

**Supplementary Table 2. PARIS2 improvements for each step.** The total library yield improvement of ~76 fold was based on experimental results ofrom 50ng starting crosslinked RNA after the 2D gel purification, not the same initial total RNA.

| | **HEK293T mRNA** | **Mice brain mRNA** | **HeLa EV-D68** |
|---|---|---|---|
| Total input reads | 42,236,572 | 25,215,538 | 624,544 |
| Primary alignments | 22,641,746 | 14,559,941 | 423,113 |
| continuous alignments | 20,184,157 | 12,652,537 | 347,625 |
| gap1 alignments | 1,179,743 | 1,087,852 | 48,658 |
| filtered gap1 alignments | 627,884 | 740,304 | 16,334 |
| gapm alignments | 18,578 | 17,880 | 2,709 |
| filtered gapm alignments | 11,436 | 13,746 | 1,376 |
| trans alignments | 1,193,628 | 775,401 | 21,470 |
| homotypic alignments | 4,342 | 4,278 | 71 |
| bad alignments | 61,298 | 21,993 | 2,580 |
| Filtered gap1+gapm+trans alignments | 1,832,948 | 1,529,451 | 39,180 |

**Supplementary Table 3. Library statistics.** The primary alignments were filtered to remove low-confidence segments, rearranged and classified into 6 types using gaptypes.py (https://github.com/zhipenglu/CRSSANT). Gap1: non-continuous alignments with one gap; gapm: non-continuous alignments with more than one gaps; trans: continuous alignments with the two arms on different strands or chromosomes; homotypic: non-continuous alignments with the two arms overlapping each other. Gap1 and gapm alignments containing splicing junctions and short 1-2 nt gaps were filtered out before further processing. Then filtered gap1 alignments, filtered gapm alignments and trans alignments were combined and used to analyze RNA structures and interactions.

| | | | |
|---|---|---|---|
| **Crosslinking oligos** | | | |
| Name | Sequence(5'-3') | Length | Related figures |
| ssDNA1-25mer | ACAGGGAAGGGTTATCCCACCTGAC | 25 nt | Supplementary Fig. 2 |
| ssDNA2-25mer | GTCAGGTGGGATAATCCTTACCTGT | 25 nt | Supplementary Fig. 2 |
| ssDNA3-25mer | ACAGGGAAGGGTTATGCCGCCTGAC | 25 nt | Supplementary Fig. 4 |
| ssDNA4-25mer | GTCAGGCGGCATAACCCTTCCCTGT | 25 nt | Supplementary Fig. 4 |
| ssDNA-8mer | CGGTACCG | 8 nt | Supplementary Fig. 11 |
| ssRNA-8mer | CGGUACCG | 8 nt | Supplementary Fig. 11 |
| **Primer extension oligos** | | | |
| Name | Sequence(5'-3') | Length | Related figures |
| RNA template | CUUGCUAGGCCCGGGUUCCUCCCGGGCCUAGCCCUG UCUGAGCGUCGC | 48 nt | Fig. 4 |
| DNA primer | GCGACGCTCAGACAGG | 16 nt | Fig. 4 |
| **Adapter ligation oligos** | | | |
| Name | Sequence(5'-3') | Length | Related figures |
| SLRNA1 | UUGGUCAACGCGAGUUGACC | 20 nt | Supplementary Fig. 9 |
| SLRNA2 | GGUCAACGCGAGUUGACCUU | 20 nt | Supplementary Fig. 9 |
| SLRNA3 | AGGUCAACGCGAGUUGACCU | 20 nt | Supplementary Fig. 9 |
| SLRNA7 | UUCCUUCCUCAACGCGAGUUGACC | 24 nt | Supplementary Fig. 9 |
| SLRNA8 | CCUCAACGCGAGUUGACCUUCCUU | 24 nt | Supplementary Fig. 9 |
| **Library generation oligos** | | | |
| Name | Sequence(5'-3') | Length | |
| Adapter | /5rApp/AGATCGGAAGAGCGGTTCAG/3ddC/ | 21 nt | |
| RT primer | /5phos/WWWNNNATCACGNNNNNTACCCTTCGCTTC ACACACAAG/iSp18/GGATCC/iSp18/TACTGAAC CGC | 56 nt | 6 nucleotides in red color are barcode. |
| P3Tall | GCATTCCTGCTGAACCGCTCTTCCGATCT | 29 nt | |
| P6Tall | TTTCCCCTTGTGTGTGAAGCGAAGGGTA | 28 nt | |
| P3Solexa | CAAGCAGAAGACGGCATACGAGATCGGTCTCGGCATT CCTGCTGAACCGCTCTTCCGATCT | 61 nt | |
| P6Solexa | AATGATACGGCGACCACCGAGATCTACACTCTTTCCC CTTGTGTGTGAAGCGAAGGGTA | 59 nt | |
| **EV-D68 antisense oligos** | | | |
| Name | Sequence(5'-3') | Length | |
| EV-D68_US47_428 | GAGGACTCTATAGTAGCTCA/3BioTEG/ | 20 nt | |
| EV-D68_US47_1647 | AAAGGTATGTTGGGACACCT/3BioTEG/ | 20 nt | |
| EV-D68_US47_2553 | AATTCTCCACTAGAGTCTCG/3BioTEG/ | 20 nt | |
| EV-D68_US47_3366 | CTGATTGCCAATCCACATAG/3BioTEG/ | 20 nt | |
| EV-D68_US47_4369 | CAAACCGGTTCAATGCGAGA/3BioTEG/ | 20 nt | |
| EV-D68_US47_5518 | GTCAAGTCTCTAAGTGCACA/3BioTEG/ | 20 nt | |
| EV-D68_US47_7042 | TTCATTGGCATCACTGGATG/3BioTEG/ | 20 nt | |
| **RT-qPCR primers** | | | |
| Name | Sequence(5'-3') | Length | |
| human-ACTB-F | AGAGCTACGAGCTGCCTGAC | 20 nt | |
| human-ACTB-R | AGCACTGTGTTGGCGTACAG | 20 nt | |
| human-GAPDH-F | CCATGAGAAGTATGACAACAGCC | 23 nt | |
| human-GAPDH-R | GGGTGCTAAGCAGTTGGTG | 19 nt | |
| human-SNORD118-F | TGGGATAATCCTTACCTGTTCCT | 23 nt | |
| human-SNORD118-R | TCCTGATTACGCAGAGACGTTA | 22 nt | |
| human-SNORD13-F | GTGATGATTGGGTGTTCATACG | 22 nt | |
| human-SNORD13-R | CACGTCGTAACAAGGTTCAAGG | 22 nt | |
| human-18SrRNA-F | CTTAGAGGGACAAGTGGCGTTC | 22 nt | |
| huamn-18SrRNA-R | ACGCTGAGCCAGTCAGTGTA | 20 nt | |
| EV-D68_US47_2C-F | GTGGAAGCAAAGAGGGTAGTAG | 22 nt | |
| EV-D68_US47_2C-R | GTTCCTGGAGAGCCATGTATTAT | 23 nt | |

**Supplementary Table 4. Oligos used in this study.**

### Methods

#### Synthesis and characterization of amotosalen HCl

Psoralen is the only class of reversible nucleic acid crosslinkers that can be used in mild physiological conditions, and AMT is the most commonly used one due to its relatively high solubility at 1mg/ml in aqueous solutions (~ 3mM). Nevertheless, crosslinking at 0.5mg/ml does not approach saturation and therefore the solubility still limits its efficiency (Calvet and Pederson 1979). It is likely that this limited solubility is responsible for the low crosslinking efficiency (0.2-0.5% crosslinked RNA from total RNA) (Lu, Zhang et al. 2016). In a related class of methods that analyzes nucleotide flexibility/accessibility, as exemplified by SHAPE and DMS-seq, the RNA-reactive compounds are typically used at much higher concentrations to merely obtain single hit kinetics (e.g. 100mM or higher for NAI-N3, and 650mM for DMS (Rouskin, Zubradt et al. 2014, Spitale, Flynn et al. 2015). In the chemical probing experiments, the reactions would destabilize RNA structures and therefore modifications should be limited to less than 1 per ~100nt. However, in the case of crosslinking, RNA structures are stabilized, and therefore higher crosslinking efficiency does not have adverse effects.

One way to improve PARIS is to use psoralen derivatives that are more water-soluble. Previous studies have shown that amotosalen (also known as S59 or S-59) is soluble at 50mg/ml in aqueous solutions (Lin, Cook et al. 1997, Wollowitz, Isaacs et al. 1997). Amotosalen (compound 2 in patent US5654443) was used at 50ug/ml, irradiated with $3 J/cm^2$ 365nm UV for inactivation of viruses and bacteria. The activity of amotosalen was slightly better than AMT at the same concentration (Wollowitz, Isaacs et al. 1997). The synthesis of amotosalen was described on page 44 of patent US5654443, but the procedure is unnecessarily complex. We synthesized amotosalen from trioxalen using a simplified three-step procedure as follows (see **Supplementary Figure 1**).

Trioxsalen + $ClCH_2OCH_3$ → CMT + methanol;
CMT + Boc-ethanolamine → Boc-amotosalen → amotosalen + Boc

*General.* All chemicals for synthesis were obtained from commercial sources and used as received unless stated otherwise. Solvents were reagent grade. Thin-layer chromatography (TLC) was performed using commercial Kieselgel 60, F254 silica gel plates. Flash chromatography was performed on silica gel (40-63 µm, 230-400 mesh). Drying of solutions was performed with $MgSO_4$ and solvents were removed with a rotary evaporator. Chemical shifts for NMR measurements were determined relative to the residual solvent peaks ($\delta_H$ 7.26 for $CHCl_3$ and 2.50 for DMSO, $\delta_C$ 77.0 for $CHCl_3$ and 40.0 for DMSO). The following abbreviations are used to indicate signal multiplicity: s, singlet; d, doublet; t, triplet; q, quartet; m, multiplet; brs, broad signal; appt, apparent triplet.

*3-(chloromethyl)-2,5,9-trimethyl-7H-furo[3,2-g]chromen-7-one* (**2**, CMT, or chloromethyl trioxalen, or 4'-chloromethyl-4,5',8-trimethyl psoralen). Compound **2** was synthesized as previously reported (Hearst, Rapoport et al. 1978). Trioxsalen (1.9 g, 4.4 mmol) was dissolved in AcOH by gently heating after which the solution was cooled back to room temperature. Chloromethyl methylether (16.0 g, 200 mmol) was added and the resulting reaction mixture was stirred at room temperature for 24 h. Next, more chloromethyl methylether (16.0 g, 200 mmol) was added and the solution was stirred at 35 °C. for 48 h. The reaction was cooled down to room temperature and allowed to stand for another 24 h. The formed precipitate was filtered off yielding 1.5 g (65%) of a white cotton-like solid. $^1H$ NMR (400 MHz, $CDCl_3$) δ 7.60 (s, 1H), 6.27 (s, 1H), 4.74 (s, 2H), 2.58 (s, 3H), 2.54 – 2.52 (m, 6H).

*2,2,2-trifluoro-N-(2-((2,5,9-trimethyl-7-oxo-7H-furo[3,2-g]chromen-3-yl)methoxy)ethyl)acetamide* (**3**, Boc-amotosalen). The conversion of CMT to amotosalen can be accomplished with a Williamson ether synthesis method. Compound **2** (1.5 g, 5.4 mmol) was mixed with *N*-(2-hydroxyethyl)trifluoroacetamide (3.0 g, 19.1 mmol) and heated for 1 h at 100 °C. The mixture was cooled down to room temperature and recrystallized from methanol yielding an off-white powder. $^1H$ NMR (400 MHz, DMSO) δ 8.30 (s, 1H), 7.70 (s, 1H), 6.31 (s, 1H), 4.62 (s, 2H), 3.52 (t, $J$ = 5.6 Hz, 2H), 3.35 (t, $J$ = 5.4 Hz, 2H), 2.46 (s, 6H), 2.43 (s, 3H).

*3-((2-aminoethoxy)methyl)-2,5,9-trimethyl-7H-furo[3,2-g]chromen-7-one hydrochloride* (Amotosalen HCl) (**1**). Compound **3** was dissolved in 0.5 M $Cs_2CO_3$ in methanol and stirred at room temperature for 16 h. The mixture was concentrated *in vacuo* and purified using flash chromatography (DCM:MeOH, 9:1) yielding yellow crystals. The product was dissolved in ethanol and the mixture was cooled on an ice bath. 1 M HCl in diethyl ether was added and the mixture was stirred for 4 h on ice. The white precipitate was collected by filtration yielding Amotosalen HCl. $^1H$ NMR (400 MHz, DMSO) δ 8.02 (s, 3H), 7.80 (s, 1H), 6.34 (s, 1H), 4.68 (s, 2H), 3.62 (t, $J$ = 5.1 Hz, 2H), 2.97 (d, $J$ = 4.9 Hz, 2H), 2.52 – 2.44 (m, 9H).

#### Measuring amotosalen solubility

Solubility of the newly synthesized amotosalen was tested in water, PBS and various other solutions. Amotosalen was previously reported to be soluble at least 50mg/ml in 0.9% NaCl (Lin, Cook et al. 1997). We dissolved amotosalen-HCl in water so that there was a large amount of insoluble solid and the solution was saturated. The saturated solution has a bright orange color. We diluted the solution 2500-fold and observed an absorbance of 7.29 at 250nm. This corresponds to 229mg/ml at room temperature, given the specific absorbance of 26,900 $M^{-1}cm^{-1}$ (similar to AMT, 25,000 $M^{-1}cm^{-1}$, and 8-MOP, 22,900 $M^{-1}cm^{-1}$) (Grass, Hei et al. 1998). We found that amotosalen-HCl is soluble in 1x PBS pH 7.4 above 100mg/ml (did not push it to the limit). However, amotosalen-HCl is partialy insoluble at 10mg/ml in the following solutions: 150mM NaCl without buffer, 100mM $CH_3COONa$ pH 5.2, and highly insoluble in 1% SDS. These tests suggest that amotosalen is incompatible with ionic solutions, except the 100mg/ml solution in PBS.

#### Cells and Animals

HEK293T and HeLa cells were purchased from ATCC and maintained in Dulbecco's modified Eagle's medium (DMEM, Gibco) + 10% fetal bovine serum (FBS, Gibco) + Pen/Strep antibiotic, in 37°C incubator with 5% CO2. Wild-type C57BL/6J mice were housed in a 12 h light/dark cycle. Mice between 4-6 weeks were used for experiments. All cell culture were handled according to protocols approved by the University of Southern California. All animals were used according to animal use protocols granted by the Institutional Animal Care and Use Committee at the University of Southern California.

**Crosslinking**

Crosslinking of cells. AMT (Sigma-Aldrich A4330) and Amotosalen were dissolved in pure water at a concentration of 1 mg/ml and 100 mg/ml, respectively. Cells cultured to 80% confluency in 10 cm dish were washed twice with 1x PBS, and then were treated with 0.5 mg/ml AMT, 0.5 mg/ml, 2.0 mg/ml or 5.0 mg/ml Amotosalen in 1x PBS for 15 min in 37°C incubator. Control cells were incubated in 1x PBS. The cells in crosslinking solution were placed on ice trays in Stratalink 2400 UV crosslinker and crosslinked for 30 min under UV365nm bulbs (Thompson and Hearst 1983). Swirl the plates every 10 min and make sure that plated are horizontal. Remove cross-linking solution after cross-linking and wash cells twice with 1x PBS.

Crosslinking of tissues. Four mice brain tissues were harvested and placed in ice-cold HBSS (Gibco, 14025076). The tissues were dissociated by passing through 5 ml pipet 20 times. After 3 times washing with 1x ice-cold HBSS, tissues were resuspended in 2 ml 0.5 mg/ml amotosalen and incubated for 15 min in dark. Tissues in crosslinking solution were placed on ice trays and crosslinked for 30 min under UV365nm bulbs.

Crosslinking of nucleic acid strands. DNA oligos, RNA oligos or total RNA samples were incubated with specific concentration of AMT or Amotosalen in 1x PBS for 5 min. Oligo or RNA samples in crosslinking solution were transferred to a clean surface with ice beneath it and placed in Stratalink 2400 UV crosslinker. Samples were crosslinked for 30 min under UV365nm bulbs.

**Extraction of crosslinked RNA**

TNA method. For each 10 cm dish cells, added 100 µl of 6 M GuSCN and lysed cells with vigorous manual shaking for 1 min. After cell were lysed into a nearly homogenous solution, cell lysate was added 12 µl of 500 mM EDTA, 60 µl of 10x PBS, and water to final volume of 600 µl. Then each sample was passed through a 25G or 26G needle about 20 times to further break the insoluble material. Proteinase K (PK) was added to final concentration of 1 mg/ml, and PK treatment was performed at 37ºC for 1 hour on a shaker at 600-900 RPM. After PK digestion, 60 µl of 3 M sodium acetate (pH 5.3), 600 µl of water-saturated phenol (pH 6.7), and 1 volume pure isopropanol were added to precipitate total nucleic acids by spinning at 15000 rpm for 20 min at 4ºC. After twice washing using 70% ethanol, total nucleic acids were resuspend in 300 µl of nuclease-free water.
For 100 µg of TNA samples, 50 units of TURBO™ DNase (Invitroge, AM2239) were added to remove DNA at 37ºC for 20 min. Then added 20 µl of 3 M sodium acetate (pH 5.3), equal volume of water-saturated phenol (pH 6.7), two volume of pure isopropanol to precipitate RNA sample by spinning 20 min at 12,000 x g at 4ºC. To compare the recovery efficiency, crosslinked RNA were also extracted using TRIzol reagent and RNeasy Mini™ kit (Qiagen, 74104) according to the manufacturer's instructions.

The PK digestion should clarify the solutions to some extent and greatly reduce turbidity. The addition of isopropanol should clarify the solution, resulting in obvious compact and stringy precipitates that contain both DNA and RNA, but little protein. Most of the TNA sample should be soluble. If there is still some insoluble material, spin down and remove it. The A260/A280 ratios of cross-linked TNA samples are usually in around 1.90, in the middle between the ratios for DNA and RNA. The A260/A230 ratios for the controls samples are usually above 2.1 and the ratios for crosslinked samples are usually below 1.9. The Tape Station profile for the TNA from cross-linked samples should show an obvious smear across the entire size range, while controls show three major peaks, namely the small RNAs, the 18S and 28S rRNAs. The controls should have a RIN number close to 10 while the cross-linked ones have a RIN number below 8.

**RNA fragmentation**

Crosslinked RNA were fragmented using ShortCut RNase III (NEB, M0245). Briefly, 10 µg of crosslinked RNA was fragmented using 10 µl of RNase III with 50 mM $MnCl_2$ and 1x supplied shortcut buffer at 37ºC for 5 min. After fragmentation, equal volume of phenol was immediately added to stop the reaction. Then one tenth volume of 3 M sodium acetate (pH 5.3), 3 µl of GlycoBlue (Invitrogen, AM9516), three volume of pure ethanol were added to precipitate RNA. Fragmented RNA was resuspended in RNase-free water and checked size distribution using Tape station. Different fragmentation condition also were tested in this study, such as different RNase III amount, different fragmentation time and different concentration of $MnCl_2$.

After 5 min of short cut digestion, reaction need to be stopped as soon as possible to get the optimal size distribution. Longer reaction time will reduce the RNA fragments size. The size distribution of fragmented crosslinked RNA can be analyzed by Bioanalyzer or TapeStation system. If using TapeStation, high sensitivity D1000 ScreenTape plus high sensitivity RNA sample buffer should be used because of short RNA size after fragmentation.

**DD2D purification of crosslinked RNA (dsRNA fragments).**

First dimension gel. Prepare 8% 1.5 mm thick denatured first dimension gel using the UreaGel system (National Diagnostics, EC-833). Loading dsRNA ladder (NEB, N0363S) as molecular weight marker. Run the first dimension gel at 30 W for 7~8 min in 0.5x TBE (Invitrogen, 15581044). After electrophoresis was finished, staining the gel with SYBR Gold in 0.5X TBE and excising each lane between 50 nt to topside from the first dimension gel. The second dimension gel can usually accommodate three gel splices.

Second dimension gel. Prepare the 16% 1.5 mm thick urea denatured second dimension gel using the UreaGel system (Lipson and Hearst 1988). Using prewarmed 0.5x TBE buffer to fill the electrophoresis chamber to facilitate denaturation of the cross-linked RNA. Run the second dimension at 30 W for 50 min to maintain high temperature and promote denaturation. Gel containing the cross-linked RNA above the diagonal from the 2D gel was excised and crushed for RNA extraction (Supplementary Note 3).

The different combination of 6%, 8%, 10% first dimension gel and 16%, 22.5% secondary dimension gel was also tested in this study.

15-well combs should be used for the first dimension gel so that each lane is narrower and the second dimension has a higher resolution. No more than 10 µg of fragmented RNA should be loaded to each line. 300 nm transillumination should be used to image the gel (254 nm epi-illumination will reverse the psoralen cross-linking). To make the second dimension gel, put the square plate horizontally and arrange gel slices in a "head-to-toe" manner with 2–5 mm gap between them. Apply 20–50 µl 0.5X TBE buffer on each gel slice to avoid air bubbles when placing the notched plate on top of the gel slices. Remove the excess TBE buffer after the cassette is assembled. Pour and gel solution from the bottom of the plates, while slightly tilting the plates to one side to avoid air bubbles building up between the plates. If there are air bubbles, use the thin loading tips to draw them out. During the second dimension gel running, the voltage started around 300 V and gradually increased to 500 V, while the current started around 100 mA and gradually decreased to 60 mA.

**Proximity Ligation**

Purified dsRNA fragments were proximity ligated by T4 RNA Ligase1 (NEB, M0437M). Briefly, 2 µl of 10x ligation buffer, 5 µl of T4 RNA Ligase, 1µl of SuperaseIn (Invitrogen, AM2696) and 1 µl of 0.1 mM ATP were added to 10 µl of purified dsRNA fragments. Ligation mixture was incubated at room temperature overnight. After ligation, the samples were boiled for 2 minutes to stop the reaction. After heat denaturation, samples were centrifuged to remove the precipitate and then precipitated by ethanol.

**Reverse crosslinking**

Proximity ligated RNA fragments were placed on a clean surface with ice beneath it. To protect RNA from UVC damage, 2 µL of 2.5 mM acridine orange was added to each sample (total volume 20 µL). Samples were irradiated with UV254nm for 30 min. After reverse crosslinking, RNA was purified with three volume of ethanol and 1 µl of GlycoBlue.

**Adapter Ligation**

Reverse crosslinked RNA were heated at 80°C for 90s, then snapped cooling on ice. To each sample, 3 µl of 10 µM ddc adapter, 1 µl of T4 RNA ligase 1, 2 µl of DMSO, 5 µl of PEG8000, 1 µl of 0.1 M DTT, 1 µl of SuperaseIn and 2 µl of 10x T4 RNA ligase buffer were added to perform adapter libation at room temperature for 3 hours. After adapter ligation, following reagents were added to remove free adapters: 3 µl of 10x RecJf buffer (NEBuffer 2, B7002S), 2 µl of RecJf (NEB, M0264S), 1 µl of 5'Deadenylase (NEB, M0331S), 1 µl of SuperaseIn, Reaction was incubated at 37°C for 1 h. Then 20 µl of water was added to each sample to make total volume of 50 µl and Zymo RNA clean and Concentrator-5 (Zymo Reasearch, R1013) was used to purify RNA.

**Reverse Transcription**

SuperScript IV (SSIV) (Invitrogen, 18090010) was used to performing reverse transcription. The reaction buffer was optimized $Mn^{2+}$ buffer (1x): 50 mM Tris-HCl (PH 8.3), 75 mM $CH_3COOK$ and 1.5 mM $MnCl_2$. 1 pmol of barcoded RT primer and 1 µl of 10 mM dNTP were added to RNA samples and heated at 65°C for 5 min in a PCR block, chill the samples one ice rapidly. Then 4 µl of 5x $Mn^{2+}$ buffer, 2 µl of 0.1 M DTT, 1 µl of SuperaseIn and 1 µl of SSIV were added to each sample. Mixed sample was incubated at 25°C for 15 min, 42°C for 10 hours, 80°C for 10 min; hold at 10°C. After reverse transcription, 1 µl RNase H and RNase A/T1 mix were added and incubated at 37°C for 30 min at 1000 rpm in a thermomixer to remove RNA. Synthesized cDNA were purified using Zymo DNA clean and Concentrator-5 (Zymo Reasearch, R41013).

**cDNA circularization and library generation**

1 µl of CircLigase™ II ssDNA Ligase (Lucigen, CL9021K), 1 µl of 50 mM $MnCl_2$ and 10x CircLigaseII™ buffer were added to cDNA sample and performed circularization at 60°C for 100 min. 80°C treatment for 10 min was followed to stop the reaction. The circularised cDNA products were directly used to library PCR. Library PCR preparation was done as described in ref. (Flynn, Zhang et al. 2016). PCR products were run on 6% native TBE gel. Gel containing DNA products from 175 bp and topside (corresponding to > 40 bp insert) was excised and crushed for DNA extraction.

**UVC damage prevention**

200 ng of RNA and cDNA were irradiated with UV254nm for 10 min and 30 min to introduce the UVC damage. cDNA sample is generated from total RNA of HEK293T cells. UVC damages were determined by ct value of RT-qPCR. Different concentration of Acridine Orange (AO) (Fisher Scientific, AC300911000), Ethidium Bromide (EB) (Invitrogen, 15585011), Proflavine (PF) (Sigma, P2508-1G), Acetone (Sigma, 650501-1L) and SYBR Gold (Invitrogen, S11494) were added to each sample to test their UVC prevention efficiency. Other conditions were also tested in this study, such as high salt concentration (1 M NaCl), denaturing agents 4M Urea and 50% formamide (Thermo Scientific, 17899).

**Effects of antioxidants on PUVA damages**

Following antioxidants were used to test RNA protection from PUVA damages. $O_2$•- scavenger: Tiron (Sigma, 172553) and MnTBAP (Sigma, 475870); •OH scavenger: Mannitol (Sigma, M4125), DMSO (Sigma, D2650) and Glycerol (Sigma, G5516); $^1O_2$ scavenger: $NaN_3$ (Sigma, S2002); General radical scavenger: Vitamin C (VC) (Sigma, 11140). Cells cultured to 70% confluency were treated with normal culture media with 0.5 mg/ml AMT and different concentration of antioxidants for 15 min in dark. After incubation, the media was replaced with 0.5 mg/ml AMT plus different antioxidants. Control cells were incubated with 1x PBS. The plates in crosslinking solution were placed on ice bed in UV crosslinker for 30 min crosslinking. After crosslinking, total RNA was extracted by TNA method. Crosslinking efficiency was analyzed by DD2D gel system. PUVA damage was determined by ct value of RT-qPCR.

**Primer extension**

48-mer RNA oligo (5'-CUUGCUAGGCCCGGGUUCCUCCCGGGCCUAGCCCUGUCUGAGCGUCGC-3') was crosslinked by 0.5 mg/ml AMT with UV365nm for 30 min, and were reverse crosslinked by UV254nm for 30 min with the protection of acridine orange. Synthesis was primed by DNA primer (5'-GCGACGCTCAGACAGG-3') annealed to the 3'-end of RNA template. Unless otherwise specified, reverse transcription reactions were performed in 20 μl volumes. Before all primer extension assay, samples were treated by heating in 10 μl of solution containing 1 pmol of RNA template, 1 pmol of DNA primer and 0.5 mM dNTP (no dNTP for TGIRT™-III Enzyme) at 65ºC for 5 min, then snap cooling on ice at least for 1 min. Following enzymes were used to extension. SuperScript (SS) II (Invitrogen, 18064022), SSIII (Invitrogen, 18080093), SSIV, TGIRT™-III Enzyme (TGIRT) (Ingex, TGIRT50), HIV recombinant reverse transcriptase (HIV) (Worthington, LS05006).

SSII: to each was added 4 μl of 5x standard reaction buffer (250 mM Tris-HCl pH 8.3, 375 mM KCl, 15 mM $MgCl_2$) or 5x $Mn^{2+}$ buffer (250 mM Tris-HCl pH 8.3, 375 mM KCl, 15 mM $MnCl_2$), 2 μl of 0.1 M DTT, 1 μl of SuperaseIn and 50 units of SSII. Samples were mixed and incubated at 42°C for 60 min, followed by 80°C for 10 min to stop the reaction.

SSIII: hybridized primer-template was added 4 μl of 5x standard reaction buffer (250 mM Tris-HCl pH 8.3, 375 mM KCl, 15 mM $MgCl_2$) or 5x $Mn^{2+}$ buffer (250 mM Tris-HCl pH 8.3, 375 mM KCl, 15 mM $MnCl_2$), 2 μl of 0.1 M DTT, 1 μl of SuperaseIn and 50 units of SSIII. Then mixed samples were incubated at 42°C for 5 min, 50°C for 60 min, followed by 80°C for 10 min to stop the reaction.

SSIV: to each was added 4 μl of 5x commercial buffer or 5x $Mn^{2+}$ buffer (250 mM Tris-HCl pH 8.3, 375 mM KCl, 2.5 mM / 7.5 mM / 15 mM $MnCl_2$), 2 μl of 0.1 M DTT, 1 μl of SuperaseIn and 50 units of SSIV. Samples were incubated at 42°C 5 min, 55°C for 60 min, followed by 80°C for 10 min to stop the reaction.

TGIRT: primer-template sample was added 4 μl of 5x standard reaction buffer (100 mM Tris-HCl pH 7.5, 250mM NaCl, 50 mM MgCl2) or 5x $Mn^{2+}$ buffer (100 mM Tris-HCl pH 7.5, 250mM NaCl, 15 mM $MnCl_2$), 2 μl of 0.1 M DTT, 1 μl of SuperaseIn and 50 units of TGIRT. Sample were mixed and incubated at 42°C for 30 min. Then 2.5 μl of 10 mM dNTPs were added to reaction and incubated at 60ºC for 2 hour (Mohr, Ghanem et al. 2013).

HIV: to each was added 4 μl of 5x standard buffer (250 mM Tris-HCl pH 8.3, 375 mM $CH_3COOK$, 250 mM $MgCl_2$) or 5x $Mn^{2+}$ buffer (250 mM Tris-HCl pH 8.3, 375 mM $CH_3COOK$, 15 mM $MnCl_2$), 2 μl of 0.1 M DTT, 1 μl of SuperaseIn and 50 units of HIV. Samples were incubated at 42°C 5 min, 55°C for 60 min, followed by 80°C for 10 min to stop the reaction.

After extension, 1 μl of 5 M NaOH was added to each tube and incubate the tubes for 3 min at 95°C. Then samples were neutralized with 1 μl of 5 M HCl. After purification with ethanol precipitation, cDNA products were separated on 20% denaturing polyacrylamide gels. The Gene Ruler was 10-bp DNA ladder (Invitrogen, 10821015).

**Virus stock preparation**

EV-D68 (US/MO/14-18947 strain, US/MO/47 for short in the paper, ATCC, VR-1823) stocks were prepared by infecting HeLa cells at 33ºC in 5% CO2 for 3-4 days until obvious CPE (cell rounding and sloughing) was observed. Then infected cells were subjected to three freezing-thawing cycles followed by centrifugation to remove cell debris. Virus titers were determined by 50% tissue culture infective dose (TCID50) assay and calculated by the Reed and Muench method. The virus stocks were stored at -80ºC for use.

**Immunostaining**

HeLa cells were grown to 50 to 70% confluence in a 24-well plate. For studies involving EV-D68 infection, cells were infected with EV-D68 at an MOI of 1 for 18 h at 33ºC. Samples were fixed with 4% paraformaldehyde (PFA) and stored at room temperature for 10 min and washed with PBS three times followed by 0.1% Triton-X incubation at 4ºC overnight. Cells were washed three times in 1x PBS before blocking with 2% bovine serum albumin in PBS (2% BSA) for 1 h. Then cells were incubated with rabbit polyclonal anti-VP1 of EV-D68 (GeneTex, GTX132313) at a final concentration of 1 μg/mL at room temperature for 1 h. Wash three times with PBS. Then a secondary goat anti-rabbit rhodamine red-X (Thermo Fisher, R-6394) at a final concentration of 1 μg/mL was added into cells for 1 h incubation. To visualize nuclei, DAPI stain (1:1000) was added for 3-5 min incubation followed by three times washing. At last, Images were taken with on a fluorescence microscope using DAPI and rhodamine filters.

**Target RNA enrichment**

mRNA enrichment. mRNA was enriched from total crosslinked RNA using Poly(A)Purist™ MAG Kit (Invitrogen, AM1922) according to the manufacturer's instructions.

Viral RNA enrichment. Viral RNA was enriched by several antisense oligos. Briefly, 200 μg total RNA extracted from EV-D68 infected cells were mixed with 200 pmol of seven biotinylated DNA oligos cocktail, which was maintained at 37℃ overnight with rotation in the hybridization buffer from ref (Chu, Quinn et al. 2012). At the end of the hybridization, 100 μL of MyOne Streptavidin C1 Dynabeads (Invitrogen, 65002) were added into the RNA-probe hybridization solution for additional 4 hour rotation at 37ºC. After five times washing, beads were resuspended in 0.2 units/μL Turbo DNase at 37°C for 20 min to degrade DNA probes followed by 80°C 90 s treatment to release all target RNA as much as possible. Released RNA was separated from beads and purified with ethanol precipitation. To test the intactness and purity of enriched RNA, we performed gel electrophoresis on a 1% agarose gel, using ssRNA (NEB, N0362S) as the ladder. Additionally, viral RNA profile was tested using TapeStation.

**Data analysis**

Preparing the masked genome indices. In order to accurately and easily analyze PARIS data, pseudogenes and multicopy genes from gencode, refGene and Dfam were masked from hg38/mm10 genome. And then single copy of them was added back as a separated "chromosome". For example, multicopy of snRNAs were masked from the basic hg38/mm10 assembly genome, and 9 snRNAs (U1, U2, U4, U5, U6, U11, U12, U4atac and U6atac) were concatenated into one reference, separated by 100nt "N"s, was added back. The curated hg38/mm10 genome contained 25 reference sequences, or "chromosomes", masked the multicopy genes and added back single copies. This reference is best suited for the PARIS analysis. The adjusted genome reference was used for mapping reads and IGV visualization. The EV_D68 viral genome (GenBank, KM851225.1) was downloaded from NCBI and manually corrected based on our viral sequencing data. After mutation identifying using GATK software, three variant sites on EV_D68 genome were corrected (2023:G->A; 2647:G->A; 3242:A->G). The curated EV_D68 genome was added to hg38 refence as an independent chromosome.

Mapping. Sequencing data were preprocessed to remove adapters form the 3'end using Trimmomatic. PCR duplicates were removed using readCollapse script from the icSHAPE pipeline (Flynn, Zhang et al. 2016). Then the library were split based on the barcodes using splitFastqLibrary from icSHAPE pipeline. 5' header were removed using Trimmomatic.
After primary preprocessing, reads were mapped to manually curated hg38 or mm10 genome using STAR program(Dobin, Davis et al. 2013). The parameters used are as follows: STAR --runThreadN 8 --runMode alignReads --genomeDir OuputPath --readFilesIn SampleFastq --outFileNamePrefix Outprefix --genomeLoad NoSharedMemory outReadsUnmapped Fastx --outFilterMultimapNmax 10 --outFilterScoreMinOverLread 0 --outSAMattributes All --outSAMtype BAM Unsorted SortedByCoordinate --alignIntronMin 1 --scoreGap 0 --scoreGapNoncan 0 --scoreGapGCAG 0 --scoreGapATAC 0 --scoreGenomicLengthLog2scale -1 --chimOutType WithinBAM HardClip --chimSegmentMin 5 --chimJunctionOverhangMin 5 --chimScoreJunctionNonGTAG 0 --chimScoreDropMax 80 --chimNonchimScoreDropMin 20.

Classify alignments. The primary mapping alignments were extracted from SampleAligned.sortedByCoord.out.bam. Then the primary mapping alignments were filtered to remove low-confidence segments, rearranged and classified into six different types using gaptypes.py (https://github.com/zhipenglu/CRSSANT). cont.sam, continuous alignments; gap1.sam, non-continuous alignments with one gap; gapm.sam, non-continuous alignments with more than one gaps; trans.sam, non-continuous alignments with the two arms on different strands or chromosomes; homo.sam, non-continuous alignments with the two arms overlapping each other; bad.sam, non-continuous alignments with complex combinations of indels and gaps. Gap1. and gapm alignments containing splicing junctions and short 1-2 nt gaps were filtered out using gapfilter.py (https://github.com/zhipenglu/CRSSANT) before further processing. Then filtered gap1.sam, filtered gapm.sam and trans.sam were used to analyze RNA structures and interactions.

Cluster alignments to groups. Filtering alignments were assembled to DGs and NGs using the crssant.py script (https://github.com/zhipenglu/CRSSANT).

**Global profiling of ribosome small subunit (SSU) analysis**

mRNA-rRNA interaction chimeric alignments were extracted using extractChimeAlign.py (https://github.com/minjiezhang-usc). The mRNA-rRNA interaction alignment can be directly loaded to IGV to visualize the binding sites of mRNAs on the 45S unit. Upstream and downstream 200 nt windows of transcription start site and transcription termination site were extracted and used to analyze the binding sites of h18 and h26 on the meta mRNA (mRNAmegaCoverage.py script).

**Global profiling of spliceosomal snRNP binding sites**

snRNA-target interaction alignments were extracted using awk command. snRNA-target interaction alignments with at least 15 nt matches for the snRNA targets were filtered using filterchimera.py. 200 nt windows around splice sites was extracted for gencode gtf file using gtf2splice.py. Chimera connecting specific snRNA regions were further extracted using sam2chimera.py script. Coverage along the 200nt windows was calculated using bedtools coverage. Meta-analysis for all windows around start 5' and 3' splice sites were performed with windowmeta.py. The Output.bedgraph can be loaded to IGV for visualization.

**Analysis of EV-D68 RNA structure conservation**

508 complete genomic sequences of EV-D68 strains were retrieved from the NIAID Virus Pathogen Database and Analysis Resource (ViPR) (http://www.viprbrc.org/). After removing duplicate sequences, 491 unique genomic sequences were remained. Manually curated US/MO/47 genome plus above 491 unique genomic sequences were used for alignment and further analysis. To analyze the structure conservation of EV-D68 RNA structures obtained from PARIS data, corresponding region of proposed structure from each sequence were extracted to perform multiple sequence alignment (MSA) using MSCULE.

The conservation of RNA secondary structure within each data set was evaluated using RNAz 2.0 (Gruber et al., 2010) by calculating the z score and the Structure Conservation Index (SCI). The following parameters were applied: --both-strands --no-shuffle --cutoff=0.5. The scoring results and consensus structure were visualized by R-chie (https://www.e-rna.org/r-chie/). Also, the phylogenetic tree of corresponding alignment was obtained by applying FastMe algorithm. For alignments with high sequence identity that cannot be analyzed by RNAz program, RNAalifold was used to predict their consensus structure.

R-scape was also used to study conserved RNA structure by measuring pairwise covariations observed in multiple sequence alignment (491 sequences plus curated US/MO/47). The following parameters were applied: --fold, the default E value 0.05. To further analyze covariation among all the enterovirus genomes, 3633 complete genomic sequences of EV strains were retrieved from ViPR after and used to evaluate covariation of proposed secondary structures.

**Analysis of structure conservation in the 5'UTR in EVA/EVB/EVC**

For EVA genomes, 1484 sequences were aligned and 499 sequences (maximum sequence number for RNAz) were selected by applying a script called rnazSelectSeqs.pl involved in RNAz program, which were analyzed by running RNAz.

For EVB genomes, 379 sequences were aligned and analyzed using RNAz.

For EVC genomes, 741 sequences were aligned and 499 sequences (maximum sequence number for RNAz) were selected by applying a script called rnazSelectSeqs.pl involved in RNAz program, which were analyzed by running RNAz.

rnazSelectSeqs.pl is one of the scripts involved in RNAz program, which is used for optimization of mean pairwise identity (MPI) for alignment (default:80). Additionally, too similar sequences (more than 99%) are removed.

**Data availability**

The raw and processed PARIS sequencing data was deposited to NCBI GEO with accession number GSE149493 (https://www.ncbi.nlm.nih.gov/geo/query/acc.cgi?acc=GSE149493) with the accession code 'sfitmsyejlsbnsv'.

**Code availability**

PARIS2 analysis scripts are available on GitHub partly at https://github.com/zhipenglu/CRSSANT, partly at https://github.com/minjiezhang-usc/PARIS2. The engineered genome reference of hg38/mm10 are available at https://drive.google.com/open?id=1wHSC-mf1jNNClXrVqMugqVmDVT4Crxzz.

**References**

Calvet, J. P. and T. Pederson (1979). "Heterogeneous nuclear RNA double-stranded regions probed in living HeLa cells by crosslinking with the psoralen derivative aminomethyltrioxsalen." Proc Natl Acad Sci U S A **76**(2): 755-759.
Chu, C., J. Quinn and H. Y. Chang (2012). "Chromatin isolation by RNA purification (ChIRP)." J Vis Exp(61).
Dobin, A., C. A. Davis, F. Schlesinger, J. Drenkow, C. Zaleski, S. Jha, P. Batut, M. Chaisson and T. R. Gingeras (2013). "STAR: ultrafast universal RNA-seq aligner." Bioinformatics **29**(1): 15-21.
Flynn, R. A., Q. C. Zhang, R. C. Spitale, B. Lee, M. R. Mumbach and H. Y. Chang (2016). "Transcriptome-wide interrogation of RNA secondary structure in living cells with icSHAPE." Nat Protoc **11**(2): 273-290.
Grass, J. A., D. J. Hei, K. Metchette, G. D. Cimino, G. P. Wiesehahn, L. Corash and L. Lin (1998). "Inactivation of leukocytes in platelet concentrates by photochemical treatment with psoralen plus UVA." Blood **91**(6): 2180-2188.
Hearst, J. E., H. Rapoport, S. Isaacs and C.-K. J. Shen (1978). Psoralens, Google Patents.
Lin, L., D. N. Cook, G. P. Wiesehahn, R. Alfonso, B. Behrman, G. D. Cimino, L. Corten, P. B. Damonte, R. Dikeman, K. Dupuis, Y. M. Fang, C. V. Hanson, J. E. Hearst, C. Y. Lin, H. F. Londe, K. Metchette, A. T. Nerio, J. T. Pu, A. A. Reames, M. Rheinschmidt, J. Tessman, S. T. Isaacs, S. Wollowitz and L. Corash (1997). "Photochemical inactivation of viruses and bacteria in platelet concentrates by use of a novel psoralen and long-wavelength ultraviolet light." Transfusion **37**(4): 423-435.
Lipson, S. E. and J. E. Hearst (1988). "Psoralen Cross-Linking of Ribosomal-Rna." Methods in Enzymology **164**: 330-341.
Lu, Z., Q. C. Zhang, B. Lee, R. A. Flynn, M. A. Smith, J. T. Robinson, C. Davidovich, A. R. Gooding, K. J. Goodrich, J. S. Mattick, J. P. Mesirov, T. R. Cech and H. Y. Chang (2016). "RNA Duplex Map in Living Cells Reveals Higher-Order Transcriptome Structure." Cell **165**(5): 1267-1279.
Mohr, S., E. Ghanem, W. Smith, D. Sheeter, Y. Qin, O. King, D. Polioudakis, V. R. Iyer, S. Hunicke-Smith, S. Swamy, S. Kuersten and A. M. Lambowitz (2013). "Thermostable group II intron reverse transcriptase fusion proteins and their use in cDNA synthesis and next-generation RNA sequencing." RNA **19**(7): 958-970.
Rouskin, S., M. Zubradt, S. Washietl, M. Kellis and J. S. Weissman (2014). "Genome-wide probing of RNA structure reveals active unfolding of mRNA structures in vivo." Nature **505**(7485): 701-705.
Spitale, R. C., R. A. Flynn, Q. C. Zhang, P. Crisalli, B. Lee, J. W. Jung, H. Y. Kuchelmeister, P. J. Batista, E. A. Torre, E. T. Kool and H. Y. Chang (2015). "Structural imprints in vivo decode RNA regulatory mechanisms." Nature **519**(7544): 486-490.
Sutherland, B. M. and J. C. Sutherland (1969). "Mechanisms of inhibition of pyrimidine dimer formation in deoxyribonucleic acid by acridine dyes." Biophys J **9**(3): 292-302.
Thompson, J. F. and J. E. Hearst (1983). "Structure of E. coli 16S RNA elucidated by psoralen crosslinking." Cell **32**(4): 1355-1365.
Wollowitz, S., S. T. Isaacs, H. Rapoport and H. P. Spielmann (1997). Compounds for the photo decontamination of pathogens in blood, Google Patents.

**PARIS2 Protocol**

**1. Psoralen (AMT) Crosslinking:**

1) Wash 10 cm dish cells with 1X PBS twice;
2) Add 200 μL 2X PBS, 200 μL 1 mg/mL AMT to each dish;
3) Put cells at 37ºC for 15 mins;
4) Place ice trays in the cross-linker and put cell dish on ice. Irradiate cells with 365 nm UV for 30 mins. Swirl the plates every 10 mins and make sure that they are horizontal.
5) Remove cross-linking solution after cross-linking and wash cells twice with 1x PBS. (*see* **Note 1**)

**2. TNA (total nucleic acid) extraction from psoralen crosslinked cells:**

6) For each 10 cm dish cells, add 100 μL of 6 M GuSCN, lyse cells with vigorous manual shaking for 1 min. The cells should be lysed into a nearly homogenous solution, which may not be entirely clear. Be careful, as the 6 M GuSCN is highly corrosive.
7) Then to each tube add 12 μL of 500 mM EDTA, 60 μL of 10x PBS, and bring the volume to 600 μL with water. This dilution of the sample will lead to some insoluble material. Then pass the sample through a 25G or 26G needle about 20 times to further break the insoluble material.
8) Add proteinase K to 1 mg/ml (30 μL from the 20 mg/mL stock), mix well and incubate at 37 ºC for 1 hour on a shaker (eg: Thermomixer C), at 600-900 RPM. Manually shake the tubes a few times during the incubation to facilitate mixing.
9) After PK digestion, add 60 μL of 3 M sodium acetate (pH 5.3), 600 μL of water-saturated phenol (pH 6.7), mix well divide into two tubes and then to each tube add 600 μL of pure isopropanol. (*see* **Note 2**)
10) Spin down the precipitate at 15000 rpm for 20 min at 4 ºC and remove supernantant (dispose of phenol waster properly).
11) Wash the precipitate with 70% ethanol twice to remove residual phenol and other contaminants. In each wash, mix well and shake vigorously before spinning down.
12) Combine the TNA pellets from two tubes and resuspend in 300 μL od nuclease-free water for each 10 cm plate of cells.
13) Determine the concentration and quality of the TNA sample using Nanodrop and Tape station. (*see* **Note 3**)

**3. DNase I Treatment:**

14) Transfer 100 μg of TNA samples to a new tube. Add 20 μL of 10X TURBO™ DNase Buffer, 25 μL of TURBO™ DNase (2 Units/μL). Bring each sample to a final reaction volume of 200 μL using $H_2O$.
15) Incubate samples at 37ºC for 20 min.
16) Add 20 μL of 3 M sodium acetate (pH 5.3), 220 μL of water-saturated phenol (pH 6.7), 450 μL of pure isopropanol, mix well. Spin 20 mins at 12,000 x g at 4 ºC. Wash pellet twice with 70% Ethanol. (*see* **Note 4**)
17) Resuspend RNA samples in 50 μL of RNase-free water.

**4. Shortcut Digestion:**

18) Transfer 10 μg of DNase treated RNA sample to a new tube.
19) Add ShortCut mix (tabulated below) to each sample and incubate at 37ºC for 5 mins;

| Component | Amount (μL) | Final Concentration |
|---|---|---|
| 10x ShortCut buffer, | 4 μL | 1x |
| 50 mM $MnCl_2$ | 4 μL | 5 mM |
| ShortCut RNase III | 10 μL | 0.5 U/μL |
| RNase-free water | Up to 40 μL | |

20) Add 4 μL of 3 M sodium acetate (pH 5.3), 3 μL of GlycoBlue, 60 μL of phenol, 360 μL of pure ethanol, mix well. Spin 20 mins at 12,000 x g at 4 ºC. Wash pellet twice with 70% Ethanol. (*see* **Note 5**)
21) Resuspend RNA in 10 μL of RNase-free water. Determine concentration of the samples by spectrophotometer and analyze size distribution using Tape station. (*see* **Note 6**)

**5. 2D gel purification:**

**5.1 First dimension gel:**

22) Prepare the 8% 1.5 mm thick denatured first dimension gel using the UreaGel system. For 10 mL gel solution, use 3.2 mL of UreaGel concentrate, 5.8 mL of UreaGel diluent, 1 mL of UreaGel buffer, 4 µL of TEMED, and 80 µL of 10% APS. Add TEMED and APS right before pouring the gel.
23) Use 15-well combs so that each lane is narrower and the second dimension has a higher resolution.
24) To each 10 µL sample add 10 µL GBLII loading dye. Load 200 ng dsRNA ladder as molecular weight marker. Run the first dimension gel at 30 W for 7~8 mins in 0.5X TBE.
25) After electrophoresis finishes, stain the gel with 2 µL of SYBR Gold in 20 mL 0.5X TBE, incubate for 5 min. Image the gel using 300 nm transillumination (not the 254 nm epi-illumination, which reverses the psoralen cross-linking). Excise each lane between 50 nt to topside from the first dimension gel. The second dimension gel can usually accommodate three gel splices.

**5.2 Second dimension gel:**

26) Prepare the 16% 1.5 mm thick urea denatured second dimension gel using the UreaGel system. For 20 mL gel solution, use 12.8 mL UreaGel concentrate, 5.2 mL UreaGel diluent, 2 mL UreaGel buffer, 8 µL TEMED, and 160 µL 10% APS.
27) To make the second dimension gel, put the square plate horizontally and arrange gel slices in a “head-to-toe” manner with 2–5 mm gap between them. Leave 1 cm space at the top of the notched plate so that the second dimension gel would completely encapsulate the first dimension gel slices.
28) Apply 20–50 µL 0.5X TBE buffer on each gel slice to avoid air bubbles when placing the notched plate on top of the gel slices.
29) Remove the excess TBE buffer after the cassette is assembled, and leave 2 mm space at the bottom of the notched plate to facilitate pouring the second dimension gel.
30) Pour and gel solution from the bottom of the plates, while slightly tilting the plates to one side to avoid air bubbles building up between the plates. If there are air bubbles, use the thin loading tips to draw them out.
31) Use ~60ºC prewarmed 0.5X TBE buffer to fill the electrophoresis chamber to facilitate denaturation of the cross-linked RNA. Run the second dimension at 30 W for 50 min to maintain high temperature and promote denaturation. The voltage starts around 300 V and gradually increases to 500 V, while the current starts around 100 mA and gradually decreases to 60 mA.
32) After electrophoresis, stain the gel with SYBR Gold the same as the first dimension gel.

**5.3 Purification:**

33) Excise the gel containing the cross-linked RNA from the 2D gel and transfer it to a new 10 cm cell culture dish. Crush the gel by grinding with the cap of a 15 mL tube.
34) Add 300 µL crushing buffer to gel debris. Transfer the gel slurry to a 15 mL tube by shoveling with a cell scraper.
35) Add additional 1.2 mL crushing buffer and rotate at room temperature overnight.
36) Transfer ~0.5 mL gel slurry to Spin-X 0.45 µm column. Spin at room temperature, 3400X g for 1 min. Continue until all gel slurry is filtered.
37) Aliquot 500 µL of the filtered RNA sample to an Amicon 10 k 0.5 mL column. Spin at 12,000 X g for 5 min. Repeat until all of the filtered RNA sample flowed through the column.
38) Wash the column with 300 µL water and spin the column at 12,000X g for 5 min.
39) Invert and place the column in a new collection tube, and spin at 6000 X g for 5 min. Recover ~85 µL RNA from each column (~170 µL total from two columns).
40) Precipitate the RNA using the standard ethanol precipitation method, with glycogen as a carrier. Alternatively, the RNA can be purified using the Zymo RNA clean and concentrator-5 columns.
41) Reconstitute RNA in 11 µL water and dilute 1 µL RNA sample for Bioanalyzer analysis. The RNA sample should have a broad size distribution between 40 and 150 nt in the Bioanalyzer trace. The yield is typically 0.1–0.5% from 10 µg input RNA.

**6. Proximity Ligation:**

42) Add 10 µL of proximity ligation to 10 µL of RNA, mix well and incubate at 65ºC for 20 mins.

| Component | Amount (µL) | Final Concentration |
|---|---|---|
| 10x 5' DNA Adenylation Reaction Buffer | 2 µL | 1x |
| Mth RNA Ligase | 2 µL | 100 pmol |
| SUPERase In | 1 µL | 1 U/µL |
| RNase-free water | 5 µL | |

43) Inactivate the enzyme by incubation at 85°C for 5 minutes.

44) Add Proteinase K to 1 mg/mL, incubate at 37°C for 30 minutes. (*see* **Note 7**)

45) Add 2 µL of 3 M sodium acetate (pH 5.3), 2 µL of GlycoBlue, 25 µL of phenol, 60 µL of isopropanol, mix well. Spin 20 mins at 12,000 x g at 4 ºC. Wash pellet twice with 70% ethanol. Resuspend RNA in 8 µL of RNase-free water.

**7. Reverse crosslinking:**

46) To reverse the AMT cross-linking, put the samples on a clean surface with ice beneath it. Add 2 µL of 25 mM acridine orange and mix well. (*see* **Note 8**)

47) Irradiate with 254 nm UV for 30 min.

48) Transfer reverse crosslinked sample to a new tube. Add 190 µL of RNase-free water, 20 µL of 3 M sodium acetate (pH 5.3), 3 µL of GlycoBlue, 600 µL of pure ethanol, mix well. Spin 20 mins at 12,000 x g at 4 ºC. Wash pellet twice with 70% ethanol. Resuspend RNA in 6 µL of RNase-free water.

**8. Adapter Ligation**

49) Heat reverse crosslinked RNA at 80ºC for 90s, then snap cooling on ice.

50) Add 14 µL of adapter ligation mixture to 6 µL RNA and perform the adapter ligation reaction for 3 h at room temperature. (*see* **Note 9**)

| Adapter ligation mixture | Amount (µL) | Final Concentration |
|---|---|---|
| 10x T4 RNA ligase buffer | 2.0 µL | 1x |
| 0.1 M DTT | 1.0 µL | 5 mM |
| 50 v/v % PEG8000 | 5.0 µL | 12.5 % v/v |
| DMSO | 2.0 µL | 5% |
| 10 µM ddc RNA adapter | 3.0 µL | 1.5 µM |
| High Concentration T4 RNA ligase 1 | 1.0 µL | 1.5 U/µL |

43) After adapter ligation add the following reagents to remove free adapters: 3 µL of 10X RecJf buffer (NEBuffer™ 2, B7002S), 2 µL of RecJf, 1 µL of 5' deadenylase, 1 µL of SuperaseIn, and 3 of µL water. Incubate at 37 ºC for 1 h.

44) Add 20 µL of water to each sample (total volume of 50 µL) and purify RNA with Zymo RNA clean and Concentrator-5 or ethanol precipitation. Reconstitute RNA in 11 µL of RNase-free water (elute in 6 µL of water, use same 6 µL twice).

**9. Reverse Transcription**

45) To the purified RNA add 2 µL of custom RT primer (with barcode) and 1 µL of 10mM dNTPs.

46) Heat the samples to 65ºC for 5 min in a PCR block, chill the samples one ice rapidly.

47) Add 7.5 µL of reverse transcriptase mix to the RNA and heat the samples at 25 ºC for 15 min, 42 ºC for 10 hours, 80 ºC for 10 min; hold at 10 ºC.

| SuperScript IV RT Master | Amount (µL) | Final Concentration |
|---|---|---|
| 5x SSIV Mn2+ Buffer | 4.0 µL | 1x |
| 100 mM DTT | 2.0 µL | 10 mM |
| SUPERaseIn | 1.0 µL | 1 U/µL |
| SuperScript IV | 1.0 µL | 5 U/µL |

| 5x SSIV Mn2+ Buffer | Final Concentration |
|---|---|

| | |
|---|---|
| Tris-HCl (PH 8.3) | 250 mM |
| $CH_3COOK$ | 375 mM |
| $MnCl_2$ | 7.5 mM |

48) Add 1 μL RNase H and RNase A/T1 mix and incubate at 37 ºC for 30 min at 1000 rpm in a thermomixer.

49) Purify the cDNA using SPRI DNA beads. Add 2x volume of SPRI DNA beads, equal volume of isopropanol, mix well; Incubate for 5 min at RT. Let the beads settle on the magnet for 5 min. Remove the supernatant and wash the beads once with 80% ethanol (200 μL) at RT. Dry 2min. Elute twice with 8.5 μL water (recover ~16 μL).

*Or:*

Using DNA Zymo concentrator-5 columns (add 7x Binding Buffer, then equal volume (8x original) of 100% EtOH to bind, wash normal, elute in 2 x 8.5 μL of water).

**10. cDNA Circularization, Library PCR, and Sequencing**

50) Add 4 μL circularization reaction mix to the cDNA sample and incubate at 60 ºC for 100 min, followed by 80 ºC for 10 min.

| CircμLarization Mix (4 μL) | Amount (μL) | Final Concentration |
|---|---|---|
| 10x CircLigaseII Buffer | 2.0 μL | 1x |
| CircLigase II Enzyme | 1.0 μL | 5 U/μL |
| 50 mM $MnCl_2$ | 1.0 μL | 2.5 mM |

51) Add 21.4 μL of PCR Tall mix and run PCR program until exponential amplification confirmed. Transfer cDNA to optical PCR tubes (each tube should be separate so that individual tubes can be taken out of the qPCR machine when the fluorescence signal reaches a defined point).

| 2x Phusion HF mix (100 μL) | Amount (μL) |
|---|---|
| 5x HF buffer | 40.0 μL |
| 10 mM dNTP | 4.0 μL |
| Phusion | 2.0 μL |
| Water | 54.0 μL |

| PCR Tall Mix (21 μL) | Amount (μL) |
|---|---|
| P3/P6 Tall (20 μM) | 1.0 μL |
| Phusion HF 2x | 20.0 μL |
| 25x SYBR Green I | 0.4 μL |

52) Set up the following qPCR program. Choose SYBR, initial 98 ºC, 2 mins, 10 cycles of: 98 ºC, 15 s; 65 ºC, 30 s; 72 ºC, 45s, detect fluorescence at extension step (a set of nine cycles). Take sample out once amplification reaches exponential phase.

53) Transfer PCR product to 1.5 mL tube. Purify the DNA using SPRI DNA beads. Add 2x volume of SPRI DNA beads, mix well. Let the beads settle on the magnet for 5 min. Remove the supernatant and wash the beads once with 80% ethanol (200 μL) at RT. Dry 2min. Elute twice with 10.5 μL water (recover ~20 μL).

54) Repeat SPRI DNA beads purification one more time.

55) Pool elute and add 21 μL 2X PCR Solexa mix.

| PCR Solexa Mix (21 μL) | Amount (μL) |
|---|---|
| P3/P6 Solexa (20 μM) | 1.0 μL |
| Phusion HF 2x | 20.0 μL |

56) Run PCR reaction (98 ºC, 2 mins; 3 cycles of 98 ºC, 15 s; 70 ºC, 30 s; 72 ºC, 45 s; and 4 ºC on hold).

57) Purify reaction by standard Zymo concentrator-5 column protocol. Elute with 2x 8.5 μL of water and add 3 μL of Orange G loading dye.

58) Run a 6% native TBE gel at 200 V for 30 min, until the dye just ran off the gel. Loadind 50 bp ladder (NEB).

59) Stain gel in SYBR Gold for 3 min. Image gel at 0.5, 1, and 2 s exposure times. Cut out the DNA from 175 bp and above (corresponding to > 40 bp insert).

60) Use a syringe needle to punch a hole in the bottom of a 0.65 mL tube.
61) Transfer the gel slice to 0.65 mL tube and insert into a 2 mL collection tube. Spin at room temperature, 16,000X g for 5 min. The gel slice gets sheared into slurry by passing through the hole.
62) Remove the 0.65 mL tube and add 300 μL Gel elute buffer to the slurry. Shake at 55 ºC, 1000 rpm overnight in a thermomixer.
63) Pass the gel slurry through a Spin-X 0.45 μm column to recover the DNA library.
64) Add 5x volume of Zymo DNA binding buffer and flow-through Zymo concentrator-5 column. Wash with 200 μL Washing buffer once and elute twice with 8 μL water (recover ~15 μL library). Quantify library by a high sensitivity Bioanalyzer assay.
65) Barcoded libraries can be pooled together for sequencing if necessary.
66) Sequence the libraries on an Illumina sequencer using standard conditions and the P6_Custom_seqPrimer. Usually, a 70 nt single end sequencing reaction is enough for PARIS. The multiplexing and random barcodes are sequenced together with the insert.

**Notes**

1. AMT cross-linked cell pellets should have a darker color than the non-cross-linked ones.
2. The PK digestion should clarify the solutions to some extent and greatly reduce turbidity. The addition of isopropanol should clarify the solution, resulting in obvious compact and stringy precipitates that contain both DNA and RNA, but little protein.
3. TNA sample: Most of the TNA sample should be soluble. If there is still some insoluble material, spin down and remove it. The cross-linked samples yield 60-70% of TNA compared to controls, and the A260/A280 ratios are usually in around 1.90, in the middle between the ratios for DNA and RNA. The A260/A230 ratios for the controls samples are usually above 2.1 and the ratios for crosslinked samples are usually below 1.9.
The Tape Station profile for the TNA from cross-linked samples should show an obvious smear across the entire size range, while controls show three major peaks, namely the small RNAs, the 18S and 28S rRNAs. The controls should have a RIN number close to 10 while the cross-linked ones have a RIN number below 8. Alternatively use bioanalyzer to check size distribution.
4. Purification of RNA using Trizol/chloroform will lose ~50% of cross-linked RNAs. It is better to extract RNA directly using phenol/isopropanol precipitation method.
5. After 5 mins of short cut digestion, reaction need to be stopped as soon as possible. Longer reaction time will reduce the RNA fragments size.
6. Typically, the AMT cross-linked samples have a stronger tail above 100 nt than the control samples.
7. Mth RNA ligases will tightly bind target RNAs, affecting RNA recovery efficiency. Proteinase K treatment will remove Mth RNA ligases before purification, and increase the recovery efficiency.
8. UV irradiation will produce heavy RNA damage, such as cyclobutene pyrimidine dimer (CPD) and (6-4) lesion. Absorption of UV photons produce RNA singlet and triplet excited states. In this respect, the characterization of the RNA singlet states are mainly responsible for the formation of pyrimidine photoproducts. Triplet excited states only play a limited role (less than 10%) (1-2). Acridine dyes bind to double strand RNA by intercalating between adjacent base pairs or by exterior ionic bonding, and inhibit the pyrimidine dimer formation. Energy transfer from RNA to acridine is important in the reduction of dimer yields. And singlet states of RNA are responsible for this transfer (3).
9. Denature treatment and 10% of DMSO will unfold the RNA duplexes and enhance the adapter ligation efficiency.

**Reference:**

1. Banyasz , T. Douki , R. Improta , T. Gustavsson , D. Onidas , I. Vaya , M. Perron and D. Markovitsi , Electronic excited states responsible for dimer formation upon UV absorption directly by thymine strands: joint experimental and theoretical study, J. Am. Chem. Soc., 2012, 134 , 14834 —14845
2. L. Liu , B. M. Pilles , J. Gontcharov , D. B. Bucher and W. Zinth , Quantum yield of cyclobutane pyrimidine dimer formation via the triplet channel determined by photosensitization, J. Phys. Chem. B, 2016, 120 , 292 —298
3. Mechanisms of inhibition of pyrimidine dimer formation in deoxyribonucleic acid by acridine dyes. Sutherland BM, Sutherland JC. Biophys J. 1969 Mar;9(3):292-302.

**Gel Elution Buffer**

| Reagent | Quantity (for 500 mL) | Final concentration |
|---|---|---|
| Tris–HCl (1 M, pH 7.5) | 1 mL | 20 mM |
| Sodium acetate (2.5M, prepared without pH adjustment) | 5 mL | 0.25 M |
| EDTA (0.5 M, pH 8.0) | 0.1 mL | 1 mM |
| SDS (10%, w/v) | 1.25 mL | 0.25% |
| H2O | 42.65 mL | |

If the SDS precipitates, warm to 37°C until the precipitate disappears. Store indefinitely at room temperature.
